# Targeted degradation of the oncogenic phosphatase SHP2

**DOI:** 10.1101/2021.06.02.446786

**Authors:** Vidyasiri Vemulapalli, Katherine A. Donovan, Tom C.M. Seegar, Julia M. Rogers, Munhyung Bae, Ryan J. Lumpkin, Ruili Cao, Matthew T. Henke, Eric S. Fischer, Gregory D. Cuny, Stephen C. Blacklow

**Affiliations:** Department of Biological Chemistry and Molecular Pharmacology, Harvard Medical School, Boston MA, 02115; Department of Cancer Biology, Dana Farber Cancer Institute, Boston MA 02215; Department of Molecular Genetics, Biochemistry and Microbiology, University of Cincinnati College of Medicine, Cincinnati OH, 45267; Department of Pharmacological and Pharmaceutical Sciences, University of Houston, Houston, TX 77204

## Abstract

SHP2 is a protein tyrosine phosphatase that plays a critical role in the full activation of the Ras/MAPK pathway upon stimulation of receptor tyrosine kinases (RTKs), which are frequently amplified or mutationally activated in human cancer. In addition, activating mutations in SHP2 result in developmental disorders and hematologic malignancies. Several allosteric inhibitors have been developed for SHP2 and are currently in clinical trials. Here, we report the development and evaluation of a SHP2 PROTAC created by conjugating RMC-4550 with pomalidomide using a PEG linker. This molecule is highly selective for SHP2, induces degradation of SHP2 in leukemic cells at sub-micromolar concentration, inhibits MAPK signaling, and suppresses cancer cell growth. SHP2 PROTACs serve as an alternative strategy for targeting ERK-dependent cancers and are useful tools alongside allosteric inhibitors for dissecting the mechanisms by which SHP2 exerts its oncogenic activity.

---

The proto-oncogene PTPN11 encodes a cytoplasmic protein tyrosine phosphatase, SHP2, which is required for normal development. SHP2 acts downstream of multiple receptor tyrosine kinases (RTKs) to exert sustained activation of the RAS-MAPK signaling cascade. The first oncogenic phosphatase to be identified, SHP2 is dysregulated in multiple human diseases, where germline mutations cause the developmental disorders of Noonan and LEOPARD syndromes. Somatic mutations of SHP2 are found in about 35% of cases of juvenile myelomonocytic leukemia (JMML), and are seen recurrently in myelodysplastic syndrome, ALL, AML, and even in solid tumors ^1, 2^. Oncogenic mutations in SHP2 destabilize the “off” or “auto-inhibited” state of the enzyme and boost basal activity through shifting the conformational equilibrium towards a more open state ^3^, which leads to uncontrolled MAPK activation. Reduction of SHP2 activity through genetic knockdown or allosteric inhibition suppresses RAS-ERK signalling and inhibits tumor growth, validating SHP2 as a target for cancer therapy ^4^. Moreover, because SHP2 lies downstream of the T cell immunoinhibitory receptor PD-1, SHP2 inhibition may also be a viable strategy for cancer immunotherapy in combination with PD-1 blockade or with other immunomodulatory agents ^5–7^.

Structurally, SHP2 is composed of two tandem Src homology 2 (SH2) domains, N-SH2 and C-SH2, followed by a catalytic protein tyrosine phosphatase (PTP) domain and an unstructured C-terminal tail. In the basal state, the N-SH2 domain packs against and sterically occludes the active site of the PTP domain by inserting a loop into the cleft that inhibits substrate access. Upon engagement of the N-SH2 and C-SH2 domains of SHP2 with tyrosine-phosphorylated signaling proteins, SHP2 activity is induced, presumably due to an induced conformational opening that alleviates N-SH2 autoinhibition of the PTP domain active site ^8^. Cancer mutations typically occur at the interface between the N-SH2 and PTP domains and, in most cases, activate the phosphatase ^8^.

Given the importance of SHP2 in cancer therapy, there have been a number of efforts to develop SHP2-selective inhibitors. Early reports described active site-directed competitive inhibitors that had poor selectivity ^9, 10^. More recently, research groups at Novartis and Revolution Medicines developed allosteric inhibitors that are highly selective for SHP2, called SHP099 ^4, 11^ and RMC4550 ^12^, which were both shown to be effective tool compounds with nanomolar potency and pre-clinical activity in RTK- and RAS-driven cancers ^4, 12^. Currently, several SHP2 allosteric inhibitors (TNO155, JAB-3312, RMC-4630, RLY-1971, ERAS-601) are in phase I/II clinical trials for the treatment of advanced or metastatic solid tumors.

Although allosteric SHP2 inhibitors show clinical promise, recent preclinical studies highlight the potential for the emergence of non-mutational mechanisms of resistance ^13^. By acutely depleting the target protein, proteolysis targeting chimeras (PROTACs) have the potential to overcome such resistance mechanisms. By degrading the target protein, they have the additional benefit of eliminating any residual activity of the target protein associated with the inhibitor-bound state.

Here, we report the development and characterization of a proteolysis targeting chimera (PROTAC) highly selective for degradation of SHP2. Our lead compound consists of a SHP2-binding warhead (RMC-4550) tethered to an IMiD (immunomodulatory drug) derivative using a PEG linker. By increasing the linker length of our first generation PROTAC, we identified a lead compound that carries out highly selective SHP2 degradation with a low nanomolar DC50, suppresses MAPK signaling, and inhibits cancer cell growth. This SHP2-targeting PROTAC will be a valuable tool for acute depletion of SHP2 in functional studies and will be a starting point for further development of a SHP2-targeting PROTAC therapeutic.

## Materials and Methods

### Chemical Synthesis

The compounds reported in this manuscript were synthesized as described below.

**Scheme 1.**
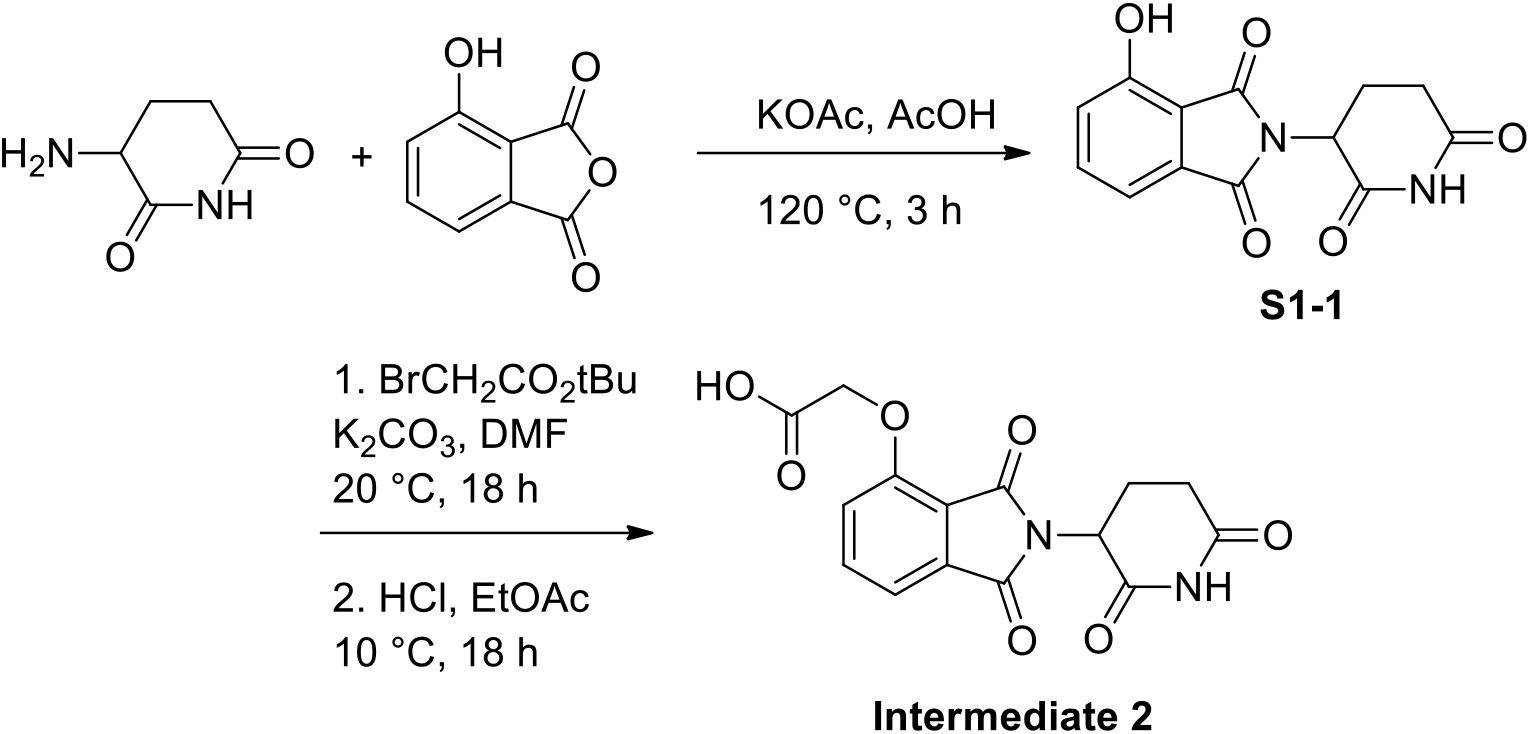

**2-(2,6-Dioxopiperidin-3-yl)-4-hydroxyisoindoline-1,3-dione (S1-1)**: A mixture of 3-aminopiperidine-2,6-dione (19.2 g, 116 mmol, 1.00 eq), 4-hydroxyisobenzofuran-1,3-dione (21.0 g, 128 mmol, 1.10 eq) and KOAc (34.3 g, 349 mmol, 3.00 eq) in AcOH (200 mL). The mixture was stirred at 120 °C for 3 h. The reaction mixture was concentrated under reduced pressure to remove solvent. The residue was diluted with water (250 mL), filtered, washed with water, and concentrated under reduced pressure to give 2-(2,6-dioxopiperidin-3-yl)-4-hydroxyisoindoline-1,3-dione (28.0 g, 101 mmol, 86.7% yield, 98.8% purity) was obtained as a gray solid.

**2-((2-(2,6-Dioxopiperidin-3-yl)-1,3-dioxoisoindolin-4-yl)oxy)acetic acid (Intermediate 2)**: To a solution of 2-(2,6-dioxopiperidin-3-yl)-4-hydroxyisoindoline-1,3-dione (28.0 g, 102 mmol, 1.00 eq) in DMF (200 mL) was added *tert*-butyl 2-bromoacetate (19.9 g, 102 mmol, 15.1 mL, 1.00 eq), KI (1.69 g, 10.2 mmol, 0.100 eq) and K_2_CO_3_ (21.2 g, 153 mmol, 1.50 eq). The mixture was stirred at 20 °C for 18 h. The reaction mixture was diluted with water (250 mL) and extracted with EtOAc (150 mL × 3). The combined organic layers were washed with saturated NaCl (200 mL), dried over anhydrous Na_2_SO_4_, filtered and concentrated under reduced pressure to give a residue. The crude product was triturated with MTBE (200 mL) at 10 °C for 3 h, and then the product was triturated with n-heptane (100 mL) at 105 °C for 3 h to give the product *tert*-butyl 2-((2-(2,6-dioxopiperidin-3-yl)-1,3-dioxoisoindolin-4-yl)oxy)acetate (36.0 g, 90.1 mmol, 88.7% yield, 97.7% purity) as a white solid.

To a solution of *tert*-butyl 2-((2-(2,6-dioxopiperidin-3-yl)-1,3-dioxoisoindolin-4-yl)oxy)acetate (5.00 g, 12.9 mmol, 1.00 eq) in EtOAc (2.00 mL) was added HCl/EtOAc (4 M, 45.0 mL, 14.0 eq). The mixture was stirred at 10 °C for 18 h. The reaction mixture was concentrated under reduced pressure to remove the EtOAc. The crude product was triturated with MTBE (200 mL) at 10 °C for 1 h, filtered and concentrated under reduced pressure. To the solid was added deionized water (100 mL). The residual aqueous solution was lyophilized to give **Intermediate 2** (4.00 g, 11.6 mmol, 90.3% yield, 96.6% purity) as a white solid.

**Scheme 2.**
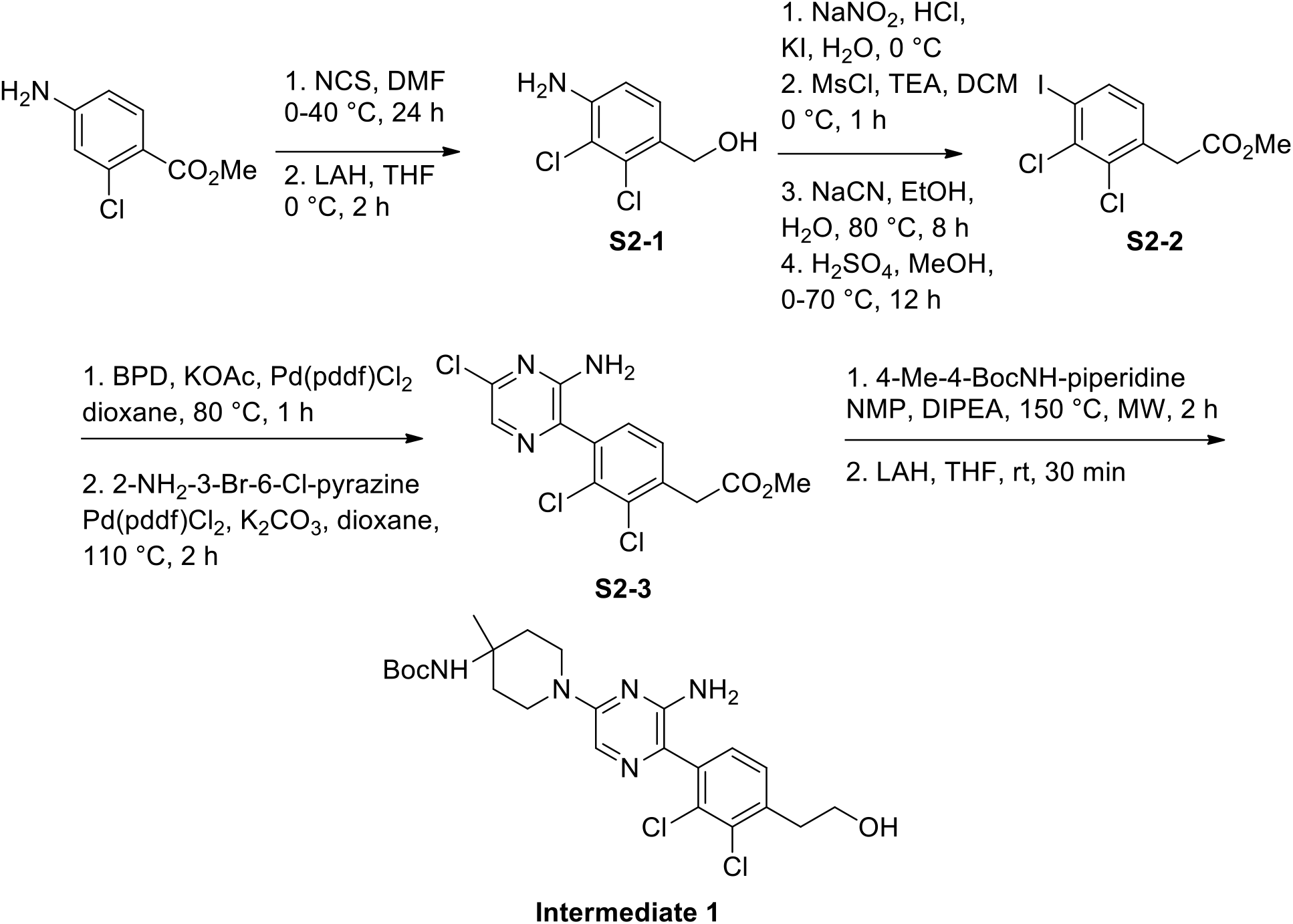

**(4-Amino-2,3-dichlorophenyl)methanol (S2-1)**: To a solution of methyl 4-amino-2-chlorobenzoate (72.0 g, 387 mmol, 1.00 eq) in DMF (360 mL) was added NCS (55.0 g, 411 mmol, 1.06 eq) at 0 °C and the mixture was stirred at 40 °C for 24 h. The reaction mixture was poured into water (500 mL) and a precipitate was formed, filtered and collected. The crude product was purified by re-crystallization from EtOH (250 mL) at 80 °C to give methyl 4-amino-2,3-dichlorobenzoate (48.0 g, 56.2% yield) as a yellow solid.

To a solution of 4-amino-2,3-dichlorobenzoate (50.0 g, 227 mmol, 1.00 eq) in THF (500 mL) was added LiAlH_4_ (18.0 g, 474 mmol, 2.09 eq) at 0 °C and the mixture was stirred at 0 °C for 2 h. To the mixture was slowly added water (23 mL), 15 % NaOH (23 mL), and MgSO_4_ (150 g) at 0°C, the mixture was stirred at 10 °C for 15 min, filtered and concentrated under reduced pressure to give a residue. The mixture was triturated with petroleum ether/EtOAc (6/1, 70 mL) at 15 °C for 12 h to give **S2-1** (36.0 g, 75.9% yield, 92.1% purity) as a light yellow solid.

**Methyl 2-(2,3-dichloro-4-iodophenyl)acetate (S2-2)**: To a solution of **S2-1** (34.0 g, 177 mmol, 1.00 eq) in H_2_O (340 mL) and HCl (12 M, 340 mL, 23.0 eq) was added a solution of NaNO_2_ (18.3 g, 265 mmol, 1.50 eq) in water (34 mL) drop-wise at 0 °C. The mixture was stirred at 0 °C for 30 min. Then a solution of KI (146 g, 885 mmol, 5.00 eq) in water (340 mL) was added drop-wise at 0°C and the resulting mixture was stirred at 0 °C for 30 min. The mixture was extracted with EtOAc (600 mL × 3). The combined organic phase was washed with brine and dried over anhydrous Na_2_SO_4_, filtered and concentrated under vacuum to give a residue. The residue was purified by column chromatography (SiO_2_, petroleum ether/EtOAc = 100/1∼20/1, R_f_ = 0.56) to give (2,3-dichloro-4-iodophenyl)methanol (25.0 g, 46.6% yield) as a yellow solid.

To a solution of (2,3-dichloro-4-iodophenyl)methanol (26.0 g, 85.8 mmol, 1.00 eq) in DCM (500 mL) and TEA (26.0 g, 257 mmol, 35.8 mL, 3.00 eq) was added MsCl (13.2 g, 115 mmol, 8.95 mL, 1.35 eq) at 0 °C. The mixture was stirred at 0 °C for 1 h. The mixture was washed with brine (300 mL × 3) and dried over anhydrous Na_2_SO_4_, filtered and concentrated under vacuum to give the corresponding mesylate (31.0 g, 94.7% yield) as a brown solid. To this material (31.0 g, 81.3 mmol, 1.00 eq) in EtOH (620 mL) was added a solution of NaCN (6.04 g, 123 mmol, 1.51 eq) in water (155 mL). The mixture was stirred at 80 °C for 8 h. The mixture was concentrated under vacuum to remove EtOH and diluted with water (150 mL), extracted with EtOAc (100 mL × 3). The combined organic phase was washed with brine (200 mL) and dried over anhydrous Na_2_SO_4_, filtered and concentrated under vacuum to give 2-(2,3-dichloro-4-iodophenyl)acetonitrile (18.0 g, 70.9% yield) as a light yellow solid.

To a solution of 2-(2,3-dichloro-4-iodophenyl)acetonitrile (17.0 g, 54.5 mmol, 1.00 eq) in MeOH (150 mL) was added H_2_SO_4_ (88.3 g, 900 mmol, 48.0 mL, 16.5 eq) at 0 °C. The mixture was stirred at 70 °C for 12 h. The reaction mixture was concentrated under reduced pressure to remove MeOH. The residue was diluted with water (150 mL) and extracted with EtOAc (100 mL × 2). The combined organic layers were washed with brine (200 mL), dried over anhydrous Na_2_SO_4_, filtered and concentrated under reduced pressure to give **S2-2** (15.0 g, 79.7% yield) as a light yellow solid.

**Methyl 2-(4-(3-amino-5-chloropyrazin-2-yl)-2,3-dichlorophenyl)acetate (S2-3):** A mixture of **S2-2** (6.00 g, 17.3 mmol, 1.00 eq), BPD (4.64 g, 18.2 mmol, 1.05 eq), AcOK (8.54 g, 86.9 mmol, 5.00 eq) and Pd(dppf)Cl_2_ (600 mg, 920 umol, 5.29x10^-2^ eq) in 1,4-dioxane (60 mL) was stirred at 80 °C for 1 h under N_2_. The mixture was filtered and the filtrate was concentrated under vacuum to give a residue. The residue was purified by column chromatography (SiO_2_, petroleum ether/EtOAc = 30/1∼10/1, R_f_ = 0.54, I_2_) to give methyl 2-(2,3-dichloro-4-(4,4,5,5-tetramethyl-1,3,2-dioxaborolan-2-yl)phenyl)acetate (5.50 g, crude) as light yellow oil.

A mixture of 3-bromo-6-chloropyrazin-2-amine (1.00 g, 4.80 mmol, 1.00 eq), methyl 2-(2,3-dichloro-4-(4,4,5,5-tetramethyl-1,3,2-dioxaborolan-2-yl)phenyl)acetate (4.97 g, 14.3 mmol, 3.00 eq), Pd(dppf)Cl_2_ (175 mg, 239 umol, 0.05 eq) and K_2_CO_3_ (663 mg, 4.80 mmol, 1.00 eq) in 1,4-dioxane (40 mL) was stirred at 110 °C for 2 h. The mixture was filtered through a pad of celite and the filtrate was concentrated under vacuum to give a residue. The residue was purified by column chromatography (SiO_2_, petroleum ether/EtOAc = 30/1∼1/1, R_f_ = 0.23) to give **S2-3** (900 mg, 51.3% yield, 94.8% purity) as a yellow solid.

*Tert*-butyl (1-(6-amino-5-(2,3-dichloro-4-(2-hydroxyethyl)phenyl)pyrazin-2-yl)-4-methylpiperidin-4-yl)carbamate (Intermediate 1): S2-3 (900 mg, 2.46 mmol, 1.00 eq), *tert*-butyl (4-methylpiperidin-4-yl)carbamate (800 mg, 3.73 mmol, 1.52 eq) and DIPEA (1.59 g, 12.3 mmol, 2.14 mL, 5.00 eq) were taken up into a microwave tube in NMP (10 mL). The sealed tube was heated at 150 °C for 2 h under microwave. The mixture was diluted with water (30 mL) and extracted with EtOAc (30 mL × 3). The combined organic phase was washed with brine (50 mL) and dried over anhydrous Na_2_SO_4_, filtered and concentrated under vacuum to give a residue. The residue was purified by column chromatography (SiO_2_, petroleum ether/EtOAc = 10/1∼1/1, R_f_ = 0.33) to give methyl 2-(4-(3-amino-5-(4-((*tert*-butoxycarbonyl)amino)-4-methylpiperidin-1-yl)pyrazin-2-yl)-2,3-dichlorophenyl)acetate (600 mg, 40.5% yield, 87.3% purity) as yellow oil.

To a solution of methyl 2-(4-(3-amino-5-(4-((*tert*-butoxycarbonyl)amino)-4-methylpiperidin-1-yl)pyrazin-2-yl)-2,3-dichlorophenyl)acetate (600 mg, 998 umol, 1.00 eq) in THF (1.5 mL) was added LiAlH_4_ (78.5 mg, 2.07 mmol, 2.07 eq). The mixture was stirred at 20 °C for 0.5 h. Next, 0.08 mL water and 0.08 mL 10% aqueous NaOH was added to the mixture at 0 °C, followed by addition 0.24 mL water and 3.00 g MgSO_4_. The mixture was stirred at 15 °C for 15 min, filtered and the filtrate was concentrated under vacuum to give a residue. The residue was purified by column chromatography (SiO_2_, petroleum ether/EtOAc = 10/1∼1/1, R_f_ = 0.23) to give **Intermediate 1** (400 mg, 74.0% yield, 91.8% purity) as yellow oil.

**Scheme 3.**
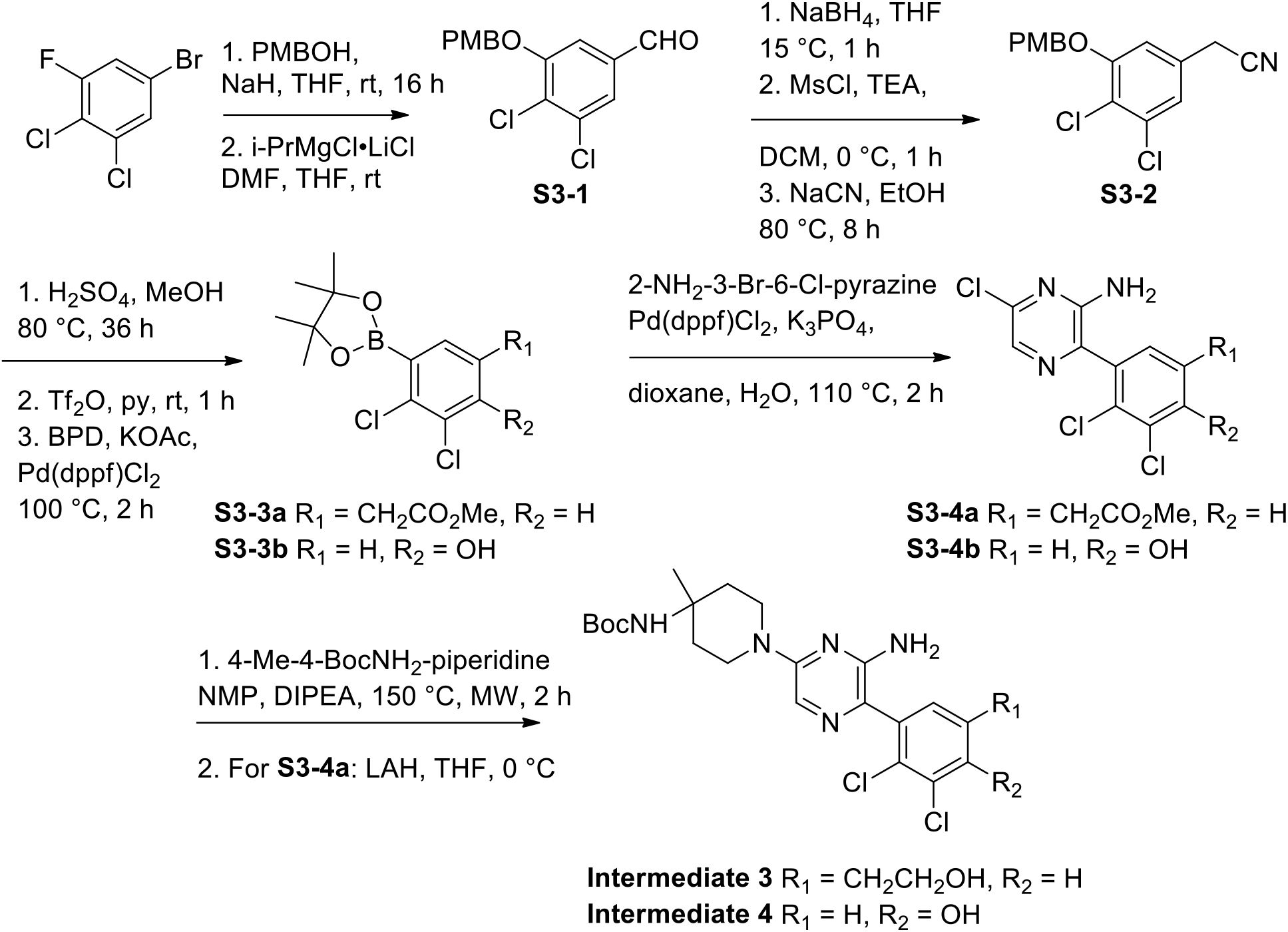

**3,4-Dichloro-5-((4-methoxybenzyl)oxy)benzaldehyde (S3-1):** To a solution of 4-methoxybenzyl alcohol (PMBOH, 30.5 g, 221 mmol, 27.5 mL, 1.20 eq) in THF (450 mL) was added NaH (10.3 g, 258 mmol, 60% purity, 1.40 eq) at 25 °C. The mixture was stirred at 25 °C for 0.5 h. Then 5-bromo-1,2-dichloro-3-fluorobenzene (45.0 g, 184 mmol, 1.00 eq) was added and the resulting mixture was stirred at 25 °C for 16 h. The mixture was quenched with 600 mL saturated aqueous NH_4_Cl and extracted with EtOAc (500 mL × 3). The combined organic phase was washed with 800 mL brine and dried over anhydrous Na_2_SO_4_, filtered and concentrated under vacuum to give a residue. The residue was purified by flash silica gel chromatography (ISCO®; 80.0 g SepaFlash® Silica Flash Column, Eluent of 0∼4% EtOAc/petroleum ether gradient) to give 5-bromo-1,2-dichloro-3-((4-methoxybenzyl)oxy)benzene (60.0 g, 89.8% yield) as a white solid.

To a solution of 5-bromo-1,2-dichloro-3-((4-methoxybenzyl)oxy)benzene (59.0 g, 162 mmol, 1.00 eq) in THF (600 mL) was added i-PrMgCl•LiCl (1.3 M, 250 mL, 2.00 eq) at 0 °C. The mixture was stirred at 25 °C for 2 h. Then DMF (35.7 g, 488 mmol, 37.6 mL, 3.00 eq) was added drop-wise and the resulting mixture was stirred at 25 °C for 1 h. The reaction mixture was quenched with 500 mL saturated aqueous NH_4_Cl and extracted with EtOAc (400 mL × 3). The combined organic phase was washed with 500 mL brine and dried over anhydrous Na_2_SO_4_, filtered and concentrated under vacuum to give a residue. The residue was triturated with 150 mL petroleum ether at 25 °C for 8 h, and then filtered to collect **S3-1** (48.0 g, 94.6% yield) as a white solid.

**2-(3,4-Dichloro-5-((4-methoxybenzyl)oxy)phenyl)acetonitrile (S3-2):** To a solution of **S3-1** (48.0 g, 154 mmol, 1.00 eq) in THF (500 mL) was added NaBH_4_ (11.6 g, 307 mmol, 2.00 eq) at 0 °C and the mixture was stirred at 15 °C for 1 h. The mixture was quenched with 400 mL water and extracted with EtOAc (300 mL × 3). The combined organic phase was washed with 500 mL brine and dried over anhydrous Na_2_SO_4_, filtered and concentrated under vacuum to give (3,4-dichloro-5-((4-methoxybenzyl)oxy)phenyl)methanol (46.0 g, 95.2% yield) as a light yellow solid.

To a solution of (3,4-dichloro-5-((4-methoxybenzyl)oxy)phenyl)methanol (46.0 g, 146 mmol, 1.00 eq) in Et_3_N (44.5 g, 440 mmol, 61.3 mL, 3.00 eq) and DCM (500 mL) was added MsCl (23.7 g, 207 mmol, 16.0 mL, 1.41 eq) at 0 °C. Then the mixture was stirred at 0 °C for 1 h. The reaction mixture was washed with brine (400 mL × 3) and dried over anhydrous Na_2_SO_4_, filtered and concentrated under vacuum to give the corresponding mesylate (37.0 g, 64.3% yield) as yellow oil. To a solution of the mesylate (37.0 g, 94.5 mmol, 1.00 eq) in EtOH (740 mL) was added a solution of NaCN (8.59 g, 175 mmol, 1.85 eq) in water (185 mL) at 25 °C. The mixture was stirred at 80 °C for 8 h. The reaction mixture was concentrated under vacuum to remove EtOH and extracted with EtOAc (200 mL × 3). The combined organic phase was washed with 300 mL brine and dried over anhydrous Na_2_SO_4_, filtered and concentrated under vacuum to give a residue. The residue was purified by column chromatography (SiO_2_, petroleum ether/EtOAc = 20/1∼3/1, R_f_ = 0.38) to give **S3-2** (16.5 g, 54.1% yield) as a white solid.

**Methyl 2-(3,4-dichloro-5-(4,4,5,5-tetramethyl-1,3,2-dioxaborolan-2-yl)phenyl)acetate (S3-3a)**: To a solution of 2-(3,4-dichloro-5-((4-methoxybenzyl)oxy)phenyl)acetonitrile 16.0 g, 49.6 mmol, 1.00 eq) in MeOH (192 mL) was added H_2_SO_4_ (117 g, 1.20 mol, 64.0 mL, 24.1 eq) at 0 °C. The mixture was stirred at 80 °C for 36 h. The mixture was poured into 600 mL saturate NaHCO_3_ and extracted with EtOAc (300 mL × 3). The combined organic phase was washed with 500 mL brine and dried over anhydrous Na_2_SO_4_, filtered and concentrated under vacuum to give a residue. The residue was purified by flash silica gel chromatography (ISCO®; 18.0 g SepaFlash® Silica Flash Column, Eluent of 0∼30% EtOAc/petroleum ether gradient, R_f_ = 0.23) to give methyl 2-(3,4-dichloro-5-hydroxyphenyl)acetate (9.30 g, 79.6% yield) as a white solid.

To a solution of methyl 2-(3,4-dichloro-5-hydroxyphenyl)acetate (8.20 g, 34.8 mmol, 1.00 eq) in pyridine (50 mL) was added Tf_2_O (10.8 g, 38.3 mmol, 6.33 mL, 1.10 eq) at 0 °C. The mixture was stirred at 20 °C for 1 h. The reaction mixture was diluted with 150 mL EtOAc and washed with 1 N HCl (100 mL × 2). The organic phase was washed with brine (100 mL × 2) and dried over anhydrous Na_2_SO_4_, filtered and concentrated under vacuum to give the triflate (12.5 g, crude) as yellow oil. To a solution of the triflate (12.5 g, 34.0 mmol, 1.00 eq) in 1,4-dioxane (130 mL) were added bis(pinacolato)diboron (BPD, 8.65 g, 34.0 mmol, 1.00 eq), Pd(dppf)Cl_2_ (2.50 g, 3.42 mmol, 0.10 eq) and KOAc (10.0 g, 102 mmol, 3.00 eq) under N_2_. The mixture was stirred at 100 °C for 2 h. The mixture was filtered through a pad of celite and the filtrate was concentrated under vacuum to give a residue. The residue was purified by column (SiO_2_, petroleum ether/EtOAc = 20/1∼4/1, R_f_ = 0.54) to give **S3-3a** (15.0 g, crude) as yellow oil.

**Methyl 2-(3-(3-amino-5-chloropyrazin-2-yl)-4,5-dichlorophenyl)acetate (S3-4a)**: To a solution of **S3-3a** (14.9 g, 43.1 mmol, 1.50 eq) in 1,4-dioxane (160 mL) were added 3-bromo-6-chloropyrazin-2-amine (6.00 g, 28.7 mmol, 1.00 eq), Pd(dppf)Cl_2_ (2.11 g, 2.88 mmol, 0.10 eq) and K_3_PO_4_ (13.0 g, 61.2 mmol, 2.13 eq) under N_2_. The mixture was stirred at 110 °C for 2 h. The reaction mixture was filtered through a pad of celite and the filtrate was concentrated under vacuum to give a residue. The residue was purified by column chromatography (SiO_2_, petroleum ether/EtOAc = 15/1∼1/1, TLC: petroleum ether/EtOAc = 2/1, R_f_ = 0.45) to give **S3-4a** (9.00 g, 90.2% yield) as a yellow solid.

*Tert*-butyl (1-(6-amino-5-(2,3-dichloro-5-(2-hydroxyethyl)phenyl)pyrazin-2-yl)-4-methylpiperidin-4-yl)carbamate (Intermediate 3): To a solution of **S3-4** (1.20 g, 3.46 mmol, 1.00 eq) in NMP (10 mL) were added *tert*-butyl (4-methylpiperidin-4-yl)carbamate (840 mg, 3.92 mmol, 1.13 eq) and DIPEA (2.24 g, 17.3 mmol, 3.02 mL, 5.00 eq). The suspension was degassed and purged with N_2_ 3 times and stirred at 150 °C for 2 h under microwave. The reaction mixture was diluted with 100 mL water and extracted with EtOAc (60 mL × 3). The combined organic phase was washed with brine (100 mL × 3) and dried over anhydrous Na_2_SO_4_, filtered and concentrated under vacuum to give a residue. The residue was purified by flash silica gel chromatography (ISCO®; 12.0 g SepaFlash® Silica Flash Column, Eluent of 0∼50% EtOAc/petroleum ether gradient, R_f_ = 0.38) to give methyl 2-(3-(3-amino-5-(4-((*tert*-butoxycarbonyl)amino)-4-methylpiperidin-1-yl)pyrazin-2-yl)-4,5-dichlorophenyl)acetate (2.60 g, crude) as a yellow solid.

To a solution of methyl 2-(3-(3-amino-5-(4-((*tert*-butoxycarbonyl)amino)-4-methylpiperidin-1-yl)pyrazin-2-yl)-4,5-dichlorophenyl)acetate (2.50 g, 4.77 mmol, 1.00 eq) in THF (30 mL) was added LiAlH_4_ (181 mg, 4.77 mmol, 1.00 eq) at 0 °C and the mixture was stirred at 0 °C for 0.5 h. The mixture was quenched with 0.15 mL water, 0.15 mL 15% NaOH and 0.3 mL water at 0 °C. Next, 2.00 g MgSO_4_ was added. The mixture was stirred at 25 °C for 30 min. Then filtered through a pad of celite and the filtrate was concentrated under vacuum to give a residue. The residue was purified by reversed-phase HPLC (0.1% formic acid condition) to give **Intermediate 3** (650 mg, 27.4% yield) as a yellow foam.

**2,3-Dichloro-4-(4,4,5,5-tetramethyl-1,3,2-dioxaborolan-2-yl)phenol (S3-3b)**: To a solution of 2,3-dichlorophenol (40.0 g, 245 mmol, 1.00 eq) in DCM (200 mL) was added Br_2_ (43.1 g, 270 mmol, 13.9 mL, 1.10 eq) over 30 mins at 0 °C. The mixture was warmed to 15 °C for 16 h. The mixture was washed with 10% aqueous Na_2_SO_3_ (240 mL) and then washed with brine (120 mL). The combined water phase was washed with DCM (150 mL). The combined organic phase was dried over anhydrous Na_2_SO_4_, filtered and concentrated under reduced pressure to give a residue. The residue was purified by column chromatography (SiO_2_, petroleum ether: EtOAc = 100: 1 to 0:1) to five 4-bromo-2,3-dichlorophenol (25.7 g, 106 mmol, 43.3% yield, 100% purity) as an off-white solid.

To a solution of 4-bromo-2,3-dichlorophenol (4.00 g, 16.5 mmol, 1.00 eq), BPD (5.46 g, 21.5 mmol, 2.53 mL, 1.30 eq) and AcOK (4.87 g, 49.6 mmol, 3.00 eq) in 1,4-dioxane (40.0 mL), was added Pd(dppf)Cl_2_•DCM (1.08 g, 1.32 mmol, 0.08 eq) under N_2_. The mixture was stirred at 90 °C for 15 h. The reaction mixture was filtered, and the residue was washed with EtOAc (100 mL × 4) and the filtrate was concentrated. The residue was purified by column chromatography (SiO_2_, petroleum ether : EtOAc = 100: 1 to 10: 1, TLC: petroleum ether : EtOAc = 5: 1, R_f_ = 0.49). to give **S3-3b** (2.50 g, 8.10 mmol, 49.0% yield, 93.6% purity) as a white solid.

**4-(3-Amino-5-chloropyrazin-2-yl)-2,3-dichlorophenol (S3-4b)**: The following reactions were done two times. To a solution of 3-bromo-6-chloropyrazin-2-amine (731 mg, 3.51 mmol, 1.00 eq) and **S3-3b** (1.30 g, 4.21 mmol, 1.20 eq) in water (2.00 mL) and 1,4-dioxane (8.00 mL) was added K_3_PO_4_ (2.61 g, 12.3 mmol, 3.50 eq) and Pd (dppf)Cl_2_ (128 mg, 175 umol, 0.05 eq). The reaction mixture was stirred at 110 °C for 1.5 h. The reaction mixture was filtered and concentrated under reduced pressure to give a residue. The residue was diluted with water (10.0 mL) and extracted with EtAcO (10.0 mL × 2). The combined organic layers were washed with brine (20.0 mL), dried over anhydrous Na_2_SO_4_, filtered and concentrated under reduced pressure to give a residue. The two batches were combined to purify by prep-HPLC (neutral condition) to give **S3-4b** (1.03 g, 3.51 mmol, 78.5% yield, 99.0% purity) as a yellow solid.

***Tert*-butyl (1-(6-amino-5-(2,3-dichloro-4-hydroxyphenyl)pyrazin-2-yl)-4-methylpiperidin-4-yl)carbamate (Intermediate 4)**: **S3-4b** (1.03 g, 3.51 mmol, 1.00 eq), *tert*-butyl (4-methylpiperidin-4-yl)carbamate (1.13 g, 5.26 mmol, 1.50 eq) and DIPEA (2.27 g, 17.6 mmol, 3.06 mL, 5.00 eq) were taken up into a microwave tube in NMP (10.0 mL). The sealed tube was heated at 150 °C for 2 h under microwave. The mixture was partitioned between water (20.0 mL) and EtOAc (20.0 mL). The water phase was separated, washed with EtOAc (20.0 mL × 4). The combined organic phase was washed with brine (10.0 mL), dried over anhydrous Na_2_SO_4_, filtered and concentrated under reduced pressure to give a residue. The residue was purified by prep-HPLC (neutral condition) to give **Intermediate 4** (700 mg, 1.39 mmol, 39.7% yield, 93.3% purity as a brown solid.

**Scheme 4.**
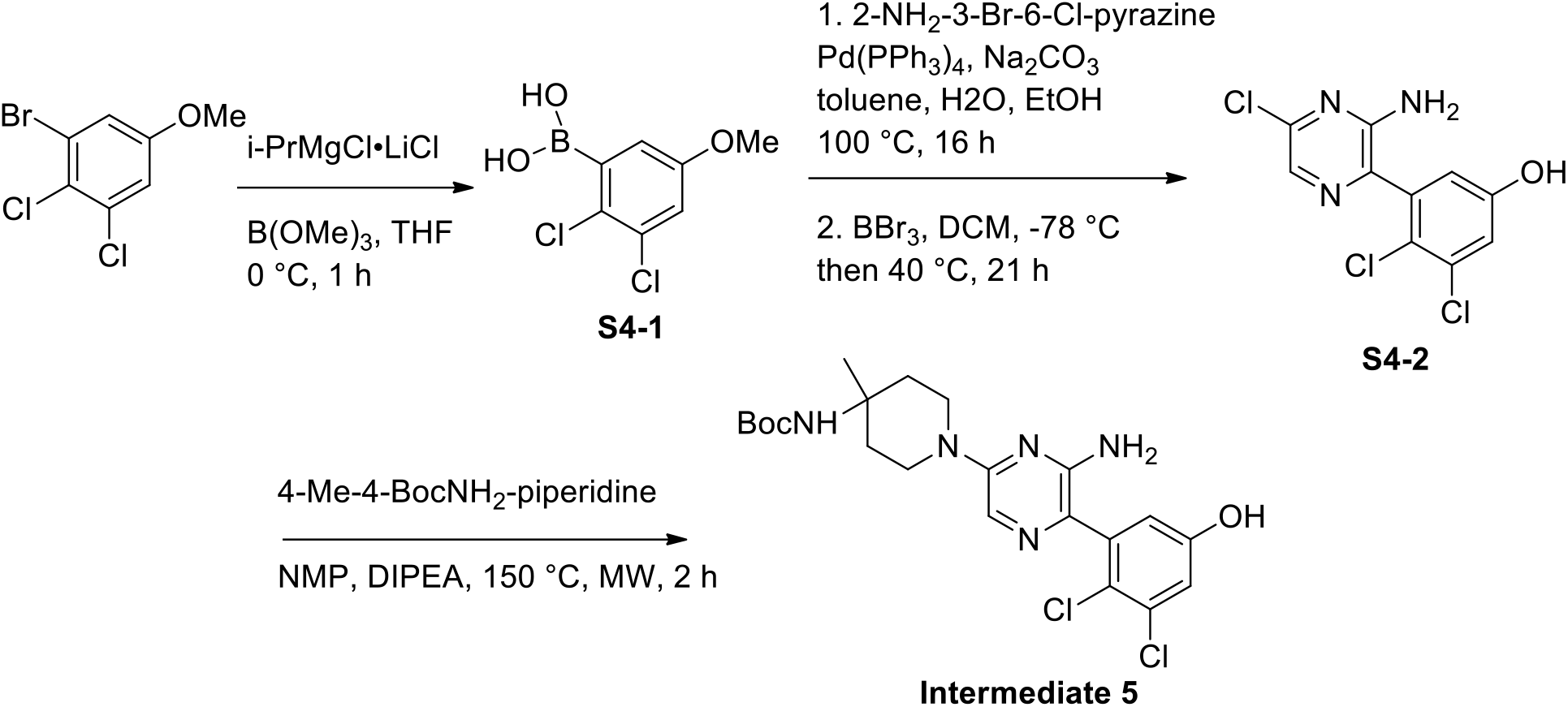

**(2,3-Dichloro-5-methoxyphenyl)boronic acid (S4-1)**: To a solution of 1-bromo-2,3-dichloro-5-methoxybenzene (5.00 g, 19.2 mmol, 1.00 eq) in THF (100 mL) was added dropwise i-PrMgCl•LiCl (1.30 M, 23.9 mL, 1.62 eq) at 0 °C under N_2_. The mixture was stirred at 0 °C for 1 h. Then trimethyl borate (9.95 g, 95.8 mmol, 10.8 mL, 5.00 eq) was added at 0 °C and the mixture stirred for 1 h. Next, HCl (1.00 M, 51.5 mL, 2.69 eq) was added and stirred at 0 °C for 1 h. The organic phase was separated, filtered and concentrated under reduced pressure to give a residue **S4-1** (4.90 g, 17.9 mmol, 93.6% yield, 80.8% purity) as a white solid.

**3-(3-Amino-5-chloropyrazin-2-yl)-4,5-dichlorophenol (S4-2)**: A mixture of **S4-1** (2.00 g, 7.32 mmol, 1.20 eq), 3-bromo-6-chloropyrazin-2-amine (1.27 g, 6.10 mmol, 1.00 eq), Pd (PPh_3_)_4_ (564 mg, 488 umol, 0.080 eq) and Na_2_CO_3_ (1.29 g, 12.20 mmol, 2.00 eq) in toluene (10.0 mL) was added water (2.00 mL) and EtOH (4.00 mL). Then the mixture was degassed and purged with N_2_ three times. The mixture was stirred at 100 °C for 16 h under a N_2_ atmosphere. The reaction mixture was partitioned between ethyl aceate (100 mL) and water (80.0 mL). The water phase was separated, washed with ethyl aceate (100 mL × 3). The combined organic phase was washed with brine (60.0 mL × 2), dried over anhydrous Na_2_SO_4_, filtered and concentrated under reduced pressure to give a residue. The residue was purified by column chromatography (SiO_2_, petroleum ether : EtOAc = 20: 1 to 3: 1, TLC: petroleum ether : EtOAc = 3: 1, R_f_ = 0.62) to give 6-chloro-3-(2,3-dichloro-5-methoxyphenyl)pyrazin-2-amine (1.70 g, 5.24 mmol, 86.0% yield, 93.9% purity) was obtained as a yellow solid.

To a solution of 6-chloro-3-(2,3-dichloro-5-methoxyphenyl)pyrazin-2-amine (1.70 g, 5.24 mmol, 1.00 eq) in DCM (85.0 mL) was added dropwise BBr_3_ (1 M, 10.5 mL, 2.00 eq) at -78 °C over 1h. After the addition, the resulting mixture was stirred at 40 °C for 21 h. The mixture was poured into ice water (150 mL) and washed with DCM (100 mL × 3). The combined organic phase was washed with brine (100 mL), dried over anhydrous Na_2_SO_4_, filtered and concentrated under reduced pressure to give a residue. The residue was purified by prep-HPLC (TFA condition) to give **S4-2** (370 mg, 1.19 mmol, 22.7% yield, 93.5% purity) as a yellow solid.

***Tert*-butyl (1-(6-amino-5-(2,3-dichloro-5-hydroxyphenyl)pyrazin-2-yl)-4-methylpiperidin-4-yl)carbamate (Intermediate 5)**: **S4-2** (520 mg, 1.79 mmol, 1.00 eq), *tert*-butyl (4-methylpiperidin-4-yl)carbamate (575 mg, 2.68 mmol, 1.50 eq) and DIPEA (1.16 g, 8.95 mmol, 1.56 mL, 5.00 eq) were taken up into a microwave tube in NMP (5.00 mL). The sealed tube was heated at 150 °C for 2 h under microwave. The combined mixture was partitioned between water (20.0 mL) and EtOAc (20.0 mL). The water phase was separated, washed with EtOAc (20.0 mL × 4). The combined organic phase was washed with brine (10.0 mL × 3), dried over anhydrous Na_2_SO_4_, filtered and concentrated under reduced pressure to give a residue. The residue was purified by column chromatography (SiO_2_, petroleum ether : EtOAc = 10: 1 to 1: 1, TLC: petroleum ether : EtOAc = 1: 1, R_f_ = 0.40) to give **Intermediate 5** (450 mg, 888 umol, 49.6% yield, 92.4% purity) as a yellow oil.

**Scheme 5.**
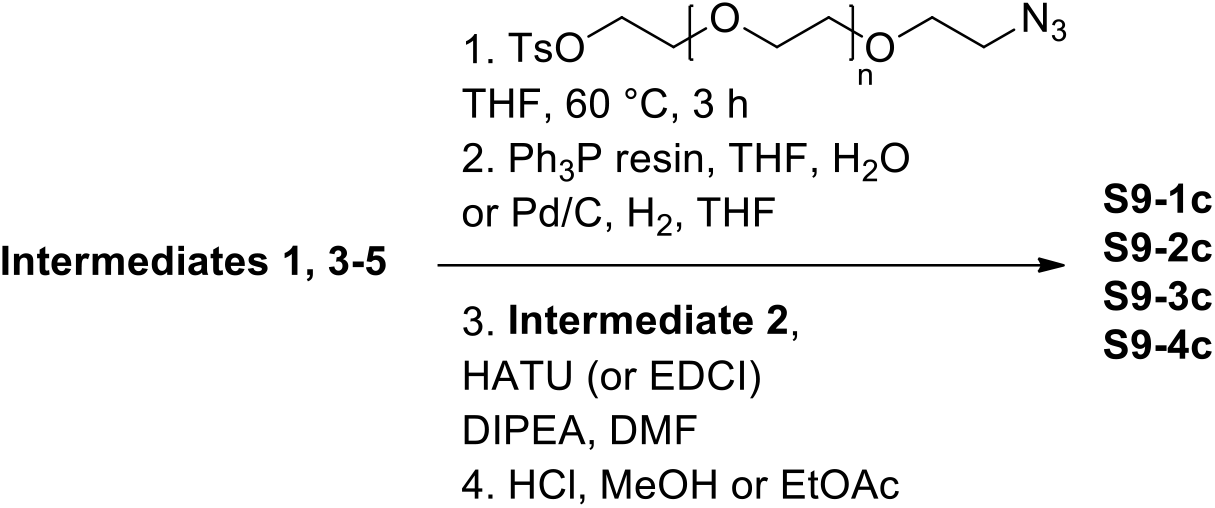

**Synthesis of S9-1c**: To a solution of **Intermediate 1** (100 mg, 184 umol, 1.00 eq) in THF (1.5 mL) was added NaH (15.0 mg, 375 umol, 60% purity, 2.03 eq) at 0 °C, followed by addition 14-azido-3,6,9,12-tetraoxatetradecyl 4-methylbenzenesulfonate (100 mg, 239 umol, 1.30 eq). The mixture was stirred at 60 °C for 3 h. The mixture was quenched with (2 mL) water and extracted with EtOAc (2 mL × 3). The combined organic phase was concentrated under vacuum to give a residue. The residue was purified by prep. TLC (dichloromethane/methanol = 10/1, TLC: dichloromethane/methanol = 10/1, R_f_ = 0.55) to give *tert*-butyl (1-(6-amino-5-(4-(17-azido-3,6,9,12,15-pentaoxaheptadecyl)-2,3-dichlorophenyl)pyrazin-2-yl)-4-methylpiperidin-4-yl)carbamate (50.0 mg, 34.0% yield, 93.4% purity) as yellow oil.

To a solution of *tert*-butyl (1-(6-amino-5-(4-(17-azido-3,6,9,12,15-pentaoxaheptadecyl)-2,3-dichlorophenyl)pyrazin-2-yl)-4-methylpiperidin-4-yl)carbamate (40.0 mg, 50.3 umol, 1.00 eq) in THF (1 mL) and water (0.3 mL) was added PPh_3_ (13.2 mg, 50.3 umol, 1.00 eq). The mixture was stirred at 60 °C for 3 h. The mixture was diluted with 5 mL EtOAc and washed with 5 mL brine. The organic phase was concentrated under vacuum to give a residue. The residue was purified by prep. TLC (dichloromethane/methanol = 10/1, TLC: dichloromethane/methanol = 10/1, R_f_ = 0.11) to give *tert*-butyl (1-(6-amino-5-(4-(17-amino-3,6,9,12,15-pentaoxaheptadecyl)-2,3-dichlorophenyl)pyrazin-2-yl)-4-methylpiperidin-4-yl)carbamate (25.0 mg, 69.3% yield) as yellow oil.

A mixture of *tert*-butyl (1-(6-amino-5-(4-(17-amino-3,6,9,12,15-pentaoxaheptadecyl)-2,3-dichlorophenyl)pyrazin-2-yl)-4-methylpiperidin-4-yl)carbamate (25.0 mg, 34.9 umol, 1.00 eq), **Intermediate 2** (19.3 mg, 52.4 umol, 1.50 eq), HATU (19.9 mg, 52.4 umol, 1.50 eq) and DIPEA (13.5 mg, 104 umol, 18.2 uL, 3.00 eq) in DMF (1 mL) was stirred at 25 °C for 0.5 h. The mixture was acidified to pH∼6 with 1M HCl and purified by prep. HPLC (column: Phenomenex Synergi C18 150*25*10um;mobile phase: [water (0.05% HCl)-acetonitrile];B%: 48%-68%, 10 min) and lyophilized to give *tert*-butyl (1-(6-amino-5-(2,3-dichloro-4-(1-((2-(2,6-dioxopiperidin-3-yl)-1,3-dioxoisoindolin-4-yl)oxy)-2-oxo-6,9,12,15,18-pentaoxa-3-azaicosan-20-yl)phenyl)pyrazin-2-yl)-4-methylpiperidin-4-yl)carbamate (15.0 mg, 41.6% yield) as a yellow solid.

A mixture of *tert*-butyl (1-(6-amino-5-(2,3-dichloro-4-(1-((2-(2,6-dioxopiperidin-3-yl)-1,3-dioxoisoindolin-4-yl)oxy)-2-oxo-6,9,12,15,18-pentaoxa-3-azaicosan-20-yl)phenyl)pyrazin-2-yl)-4-methylpiperidin-4-yl)carbamate (15.0 mg, 14.5 umol, 1.00 eq) in HCl-1,4-dioxane (4 M, 0.2 mL, 54.9 eq) and EtOAc (0.2 mL) was stirred at 20 °C for 0.5 h. The mixture was concentrated under vacuum to give a residue. The residue was purified by prep. HPLC (column: Phenomenex Synergi C18 150*25*10um; mobile phase: [water (0.05% HCl)-acetonitrile]; B%: 22%-42%, 9.5 min) and lyophilized to give **S9-1c** (2.89 mg, 20.3% yield, 95.3% purity) as a yellow solid.

**Synthesis of S9-2c**: The following reactions were done three times. To a solution of **Intermediate 3** (50.0 mg, 100 umol, 1.00 eq) in THF (1 mL) was added NaH (8.06 mg, 201 umol, 60% purity, 2.00 eq) at 0 °C and the mixture was stirred at 40 °C for 1 h. 14-Azido-3,6,9,12-tetraoxatetradecyl 4-methylbenzenesulfonate (45.1 mg, 120 umol, 1.20 eq) was added to the reaction mixture at 0 °C. The resulting mixture was stirred at 55 °C for 1 h. The mixture was quenched with 10 mL sat. NH_4_Cl. The three reaction batches were combined and the resulting mixture was extracted with EtOAc (10 mL × 3). The combined organic phase was washed with 30 mL brine and dried over Na_2_SO_4_, filtered and concentrated under vacuum to give a residue.

The residue was purified by reversed-phase HPLC (0.1% FA condition) to give *tert*-butyl (1-(6-amino-5-(5-(17-azido-3,6,9,12,15-pentaoxaheptadecyl)-2,3-dichlorophenyl)pyrazin-2-yl)-4-methylpiperidin-4-yl)carbamate (60.0 mg, 75.0% purity) as a yellow oil.

A mixture of *tert*-butyl (1-(6-amino-5-(5-(17-azido-3,6,9,12,15-pentaoxaheptadecyl)-2,3-dichlorophenyl)pyrazin-2-yl)-4-methylpiperidin-4-yl)carbamate (58.0 mg, 78.2 umol, 1.00 eq) and PPh_3_ resin (61.2 mg, 234 umol, 3.00 eq) in THF (1 mL) and water (0.3 mL) was stirred at 50 °C for 4 h. The reaction mixture was filtered and the filtrate was concentrated under vacuum to give *tert*-butyl (1-(6-amino-5-(5-(17-amino-3,6,9,12,15-pentaoxaheptadecyl)-2,3-dichlorophenyl)pyrazin-2-yl)-4-methylpiperidin-4-yl)carbamate (60.0 mg, crude) as a yellow oil.

A mixture of *tert*-butyl (1-(6-amino-5-(5-(17-amino-3,6,9,12,15-pentaoxaheptadecyl)-2,3-dichlorophenyl)pyrazin-2-yl)-4-methylpiperidin-4-yl)carbamate (60.0 mg, 83.8 umol, 1.00 eq), **Intermediate 2** (42.0 mg, 126 umol, 1.51 eq), EDCI (28.0 mg, 146 umol, 1.74 eq), HOBt (21.0 mg, 155umol, 1.85 eq) and DIPEA (43.0 mg, 332 umol, 3.97 eq) in DMF (1 mL) was stirred at 25 °C for 1 h. The reaction mixture was neutralized with 1M HCl (in DMF) and purified by prep. HPLC (column: Phenomenex Synergi C18 150*25*10um; mobile phase: [water (0.05% HCl)-acetonitrile]; B%: 43%-63%, 8 min) to give *tert*-butyl (1-(6-amino-5-(2,3-dichloro-5-(1-((2-(2,6-dioxopiperidin-3-yl)-1,3-dioxoisoindolin-4-yl)oxy)-2-oxo-6,9,12,15,18-pentaoxa-3-azaicosan-20-yl)phenyl)pyrazin-2-yl)-4-methylpiperidin-4-yl)carbamate (45.0 mg, 52.1% yield) as a yellow solid.

A mixture of *tert*-butyl (1-(6-amino-5-(2,3-dichloro-5-(1-((2-(2,6-dioxopiperidin-3-yl)-1,3-dioxoisoindolin-4-yl)oxy)-2-oxo-6,9,12,15,18-pentaoxa-3-azaicosan-20-yl)phenyl)pyrazin-2-yl)-4-methylpiperidin-4-yl)carbamate (45.0 mg, 43.6 umol, 1.00 eq) in MeOH (0.5 mL) and HCl-1,4-dioxane (4 M, 0.5 mL, 45.7 eq) was stirred at 25 °C for 0.5 h. The mixture was concentrated under vacuum to give a residue. The residue was purified by prep. HPLC (column: Phenomenex Synergi C18 150*25*10um; mobile phase: [water (0.05% HCl)-acetonitrile]; B%: 19%-39%, 6.5 min) to give **S9-2c** (32.08 mg, 77.2% yield, 97.8% purity) as a yellow solid.

**Synthesis of S9-3c**: To a solution of compound **Intermediate 4** (100 mg, 199 umol, 1.00 eq) and 17-azido-3,6,9,12,15-pentaoxaheptadecyl 4-methylbenzenesulfonate (123 mg, 239 umol, 1.20 eq) in DMF (0.500 mL) was added K_2_CO_3_ (55.1 mg, 398 umol, 2.00 eq). The mixture was stirred at 60 °C for 3 h. The reaction mixture was partitioned between water (2.00 mL) and EtOAc (2.00 mL). The water phase was separated, washed with EtOAc (2.00 mL × 2). The combined organic phase was washed with brine (2.00 mL × 2), dried over anhydrous Na_2_SO_4_, filtered and concentrated under reduced pressure to give a residue. The residue was purified by prep-HPLC (neutral condition: column: Waters Xbridge 150*25mm*5um; mobile phase: [water (10 mM NH_4_HCO_3_)-acetonitrile]; B%: 40%-70%, 10 min) to give *tert*-butyl (1-(6-amino-5-(4-((17-azido-3,6,9,12,15-pentaoxaheptadecyl)oxy)-2,3-dichlorophenyl)pyrazin-2-yl)-4-methylpiperidin-4-yl)carbamate (66.0 mg, 87.1 umol, 43.7% yield) as a yellow oil.

To a solution of *tert*-butyl (1-(6-amino-5-(4-((17-azido-3,6,9,12,15-pentaoxaheptadecyl)oxy)-2,3-dichlorophenyl)pyrazin-2-yl)-4-methylpiperidin-4-yl)carbamate (66.0 mg, 87.1 umol, 1.00 eq) in THF (1.00 mL) was added 10% Pd/C (7.00 mg, 87.1 umol, 1.00 eq) under N_2_. The suspension was degassed under vacuum and purged with H_2_ several times.

The mixture was stirred under H_2_ (15 psi) at 15 °C for 2 h. The mixture was filtered and the filtrate was concentrated under reduced pressure to afford *tert*-butyl (1-(6-amino-5-(4-((17-amino-3,6,9,12,15-pentaoxaheptadecyl)oxy)-2,3-dichlorophenyl)pyrazin-2-yl)-4-methylpiperidin-4-yl)carbamate (63.0 mg, 86.1 umol, 98.9% yield) as a brown oil, which was used without further purification.

To a solution of *tert*-butyl (1-(6-amino-5-(4-((17-amino-3,6,9,12,15-pentaoxaheptadecyl)oxy)-2,3-dichlorophenyl)pyrazin-2-yl)-4-methylpiperidin-4-yl)carbamate (63.0 mg, 86.1 umol, 1.00 eq) and **Intermediate 2** (49.3 mg, 129 umol, 1.50 eq, HCl) in DMF (1.00 mL) was added HATU (39.3 mg, 103 umol, 1.20 eq), and DIPEA (50.1 mg, 387 umol, 67.5 uL, 4.50 eq). Then the mixture was stirred at 15 °C for 13 h. The residue was purified by prep-HPLC (neutral condition: column: Waters Xbridge 150*25mm*5um; mobile phase: [water (10 mM NH_4_HCO_3_)-acetonitrile]; B%: 32%-62%, 10 min) to give *tert*-butyl (1-(6-amino-5-(2,3-dichloro-4-((1-((2-(2,6-dioxopiperidin-3-yl)-1,3-dioxoisoindolin-4-yl)oxy)-2-oxo-6,9,12,15,18-pentaoxa-3-azaicosan-20-yl)oxy)phenyl)pyrazin-2-yl)-4-methylpiperidin-4-yl)carbamate (77.0 mg, 73.6 umol, 85.5% yield) as a yellow solid.

To a solution of compound *tert*-butyl (1-(6-amino-5-(2,3-dichloro-4-((1-((2-(2,6-dioxopiperidin-3-yl)-1,3-dioxoisoindolin-4-yl)oxy)-2-oxo-6,9,12,15,18-pentaoxa-3-azaicosan-20-yl)oxy)phenyl)pyrazin-2-yl)-4-methylpiperidin-4-yl)carbamate (77.0 mg, 73.6 umol, 1.00 eq) in EtOAc (0.700 mL) was added HCl/EtOAc (4 M, 184 uL, 10.0 eq), then the mixture was stirred at 15 °C for 2 h. The mixture was concentrated under reduced pressure to give a residue. The residue was purified by prep-HPLC (HCl condition: column: Phenomenex Synergi C18 150*30mm*4um; mobile phase: [water (0.05% HCl)-acetonitrile]; B%: 6%-36%, 10 min) to give **S9-3c** (29.5 mg, 29.5 umol, 40.1% yield, 98.2% purity, HCl) as a yellow solid.

**Synthesis of S9-4c**: To a solution of **Intermediate 5** (100 mg, 197 umol, 1.00 eq) and 17-azido-3,6,9,12,15-pentaoxaheptadecyl 4-methylbenzenesulfonate (122 mg, 237 umol, 1.20 eq) in DMF (1.00 mL) was added K_2_CO_3_ (54.5 mg, 395 umol, 2.00 eq). The mixture was stirred at 60 °C for 3 h. The reaction mixture was partitioned between water (2.00 mL) and EtOAc (2.00 mL). The water phase was separated, washed with ethyl aceate (2.00 mL × 2). The combined organic phase was washed with brine (2.00 mL × 2), dried over anhydrous Na_2_SO_4_, filtered and concentrated under reduced pressure to give a residue. The residue was purified by prep-HPLC (neutral condition: column: Waters Xbridge 150*25mm* 5um; mobile phase: [water (10mM NH_4_HCO_3_)-acetonitrile]; B%: 44%-74%, 10 min). The product *tert*-butyl (1-(6-amino-5-(5-((17-azido-3,6,9,12,15-pentaoxaheptadecyl)oxy)-2,3-dichlorophenyl)pyrazin-2-yl)-4-methylpiperidin-4-yl)carbamate (79.0 mg, 104 umol, 52.9% yield) was obtained as a yellow oil.

To a solution of *tert*-butyl (1-(6-amino-5-(5-((17-azido-3,6,9,12,15-pentaoxaheptadecyl)oxy)-2,3-dichlorophenyl)pyrazin-2-yl)-4-methylpiperidin-4-yl)carbamate (79.0 mg, 104 umol, 1.00 eq) in THF (1.00 mL) was added 10% Pd/C (8.00 mg, 10.4 umol, 0.100 eq) under N_2_. The suspension was degassed under vacuum and purged with H_2_ several times. The mixture was stirred under H_2_ (15 psi) at 15 °C for 3 h. The reaction mixture was filtered and concentrated under reduced pressure to give the product *tert*-butyl (1-(6-amino-5-(5-((17-amino-3,6,9,12,15-pentaoxaheptadecyl)oxy)-2,3-dichlorophenyl)pyrazin-2-yl)-4-methylpiperidin-4-yl)carbamate (80.0 mg, 93.9 umol, 90.1% yield, 85.9% purity) as a yellow oil.

To a solution of *tert*-butyl (1-(6-amino-5-(5-((17-amino-3,6,9,12,15-pentaoxaheptadecyl)oxy)-2,3-dichlorophenyl)pyrazin-2-yl)-4-methylpiperidin-4-yl)carbamate (80.0 mg, 93.9 umol, 1.00 eq) and **Intermediate 2** (53.8 mg, 141 umol, 1.50 eq, HCl) in DMF (1.00 mL) was added HATU (42.9 mg, 113 umol, 1.20 eq) and DIPEA (54.6 mg, 423 umol, 73.6 uL, 4.50 eq). The mixture was stirred at 15 °C for 16 h. The residue was purified by prep-HPLC (neutral condition: column: Phenomenex luna C18 150*25mm*10um; mobile phase: [water (10 mM NH_4_HCO_3_)-acetonitrile]; B%: 32%-62%, 10 min). The product *tert*-butyl (1-(6-amino-5-(2,3-dichloro-5-((1-((2-(2,6-dioxopiperidin-3-yl)-1,3-dioxoisoindolin-4-yl)oxy)-2-oxo-6,9,12,15,18-pentaoxa-3-azaicosan-20-yl)oxy)phenyl)pyrazin-2-yl)-4-methylpiperidin-4-yl)carbamate (77.0 mg, 73.6 umol, 78.4% yield) was obtained as a yellow solid.

To a solution of *tert*-butyl (1-(6-amino-5-(2,3-dichloro-5-((1-((2-(2,6-dioxopiperidin-3-yl)-1,3-dioxoisoindolin-4-yl)oxy)-2-oxo-6,9,12,15,18-pentaoxa-3-azaicosan-20-yl)oxy)phenyl)pyrazin-2-yl)-4-methylpiperidin-4-yl)carbamate (77.0 mg, 73.6 umol, 1.00 eq) in EtOAc (1.00 mL) was added HCl/EtOAc (4 M, 184 uL, 10.0 eq). The mixture was stirred at 15 °C for 2 h. The reaction mixture was filtered and concentrated under reduced pressure to give a residue. The residue was purified by prep-HPLC (HCl condition: column: Phenomenex Synergi C18 150*25*10um; mobile phase: [water (0.05% HCl)-acetonitrile]; B%: 23%-43%, 11 min). The product compound **S9-4c** (33.6 mg, 33.9 umol, 46.1% yield, 99.1% purity, HCl) was obtained as a yellow solid.

**Scheme 6.**
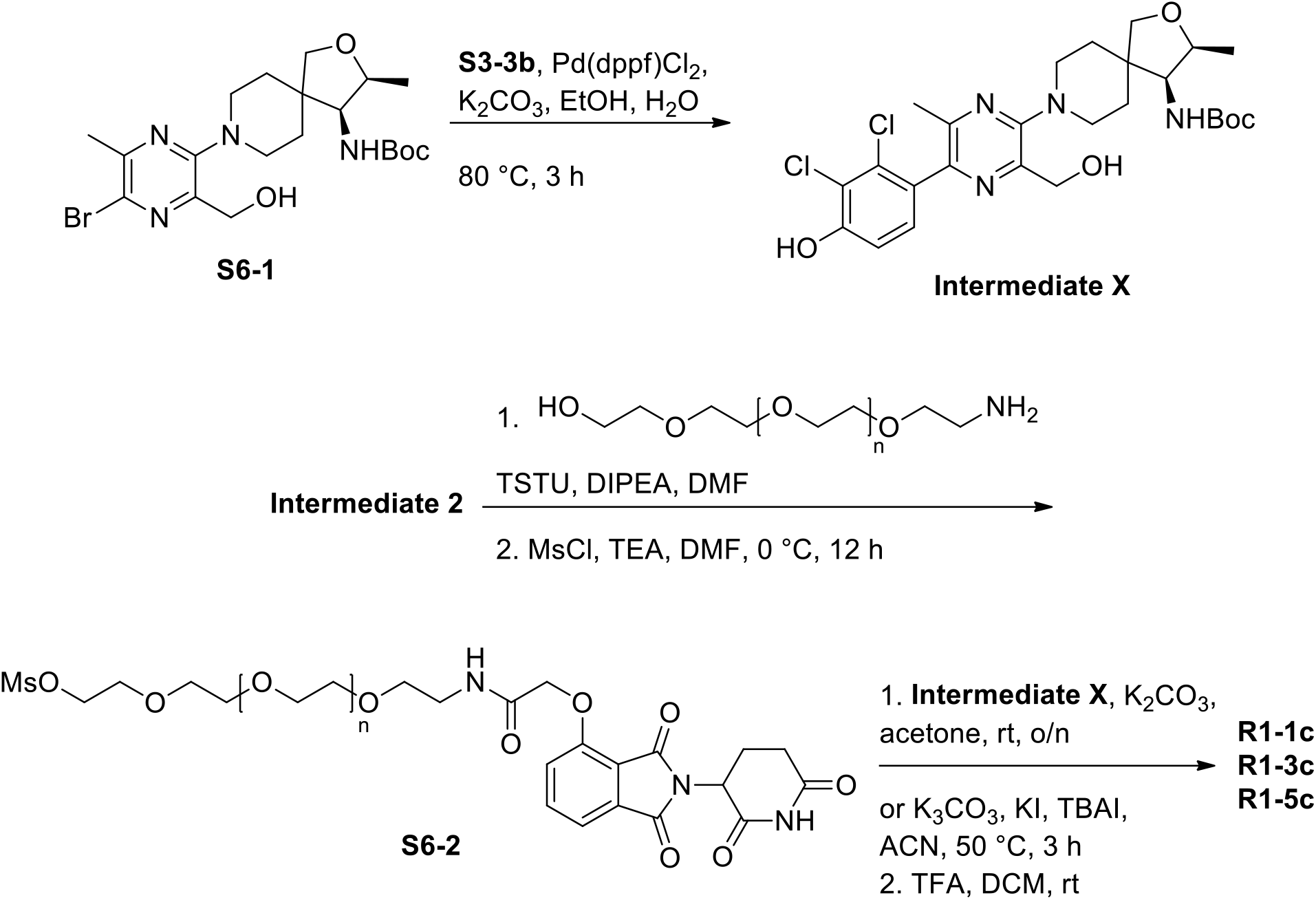

***Tert*-butyl ((3S,4S)-8-(5-(2,3-dichloro-4-hydroxyphenyl)-3-(hydroxymethyl)-6-methylpyrazin-2-yl)-3-methyl-2-oxa-8-azaspiro[4.5]decan-4-yl)carbamate (Intermediate X):** The mixture of previously reported^14^ *tert*-butyl N-[(3S,4S)-8-[5-bromo-3-(hydroxymethyl)-6-methylpyrazin-2-yl]-3-methyl-2-oxa-8-azaspiro[4.5]decan-4-yl]carbamate (480 mg, 1.02 mmol, 1.00 equiv), 2,3-dichloro-4-(4,4,5,5-tetramethyl-1,3,2-dioxaborolan-2-yl)phenol (359 mg, 1.24 mmol, 1.22 equiv), K_2_CO_3_ (432 mg, 3.13 mmol, 3.07 equiv) and Pd(dppf)Cl_2_ (97 mg, 0.132 mmol, 0.13 equiv) in EtOH (12 mL) and water (2.4 mL) was stirred for 3 h at 80 °C under a nitrogen atmosphere. The mixture was allowed to cool to room temperature. The reaction was quenched by the addition of water (30 mL). The resulting mixture was extracted with DCM (3 x 30 mL). The combined organic layers were washed with brine (30 mL), dried over anhydrous Na_2_SO_4_. After filtration, the filtrate was concentrated under reduced pressure. The residue was purified by silica gel column chromatography, eluted with DCM / MeOH (9:1) to afford *tert*-butyl N-[(3S,4S)-8-[5-(2,3-dichloro-4-hydroxyphenyl)-3-(hydroxymethyl)-6-methylpyrazin-2-yl]-3-methyl-2-oxa-8-azaspiro[4.5]decan-4-yl]carbamate (**Intermediate X**, 400 mg, 60%) as a brown solid.

**General synthesis of S6-2. Exemplified for n=1**. To a stirred mixture of **Intermediate 2** (6.50 g, 19.56 mmol, 1.00 equiv) and TSTU (8.8 g, 29.34 mmol, 1.50 equiv) in DMF (65 mL) was added DIPEA (5.0 g, 39.12 mmol, 2.00 equiv) dropwise at 0 °C under a nitrogen atmosphere. The resulting mixture was stirred for overnight at room temperature. The reaction was quenched with water/ice at room temperature. The resulting mixture was extracted with EtOAc (3 x 100 mL). The combined organic layers were washed with brine (100 mL), dried over anhydrous Na2SO4. After filtration, the filtrate was concentrated under reduced pressure. This resulted in 2,5-dioxopyrrolidin-1-yl 2-[[2-(2,6-dioxopiperidin-3-yl)-1,3-dioxoisoindol-4-yl]oxy]acetate (4.0 g, 38%) as a white solid.

To a stirred mixture of 2,5-dioxopyrrolidin-1-yl 2-[[2-(2,6-dioxopiperidin-3-yl)-1,3-dioxoisoindol-4-yl]oxy]acetate (2.00 g, 4.65 mmol, 1.00 equiv) and 2-[2-[2-(2-aminoethoxy)ethoxy]ethoxy]ethanol (0.90 g, 4.65 mmol, 1.00 equiv) in DMF (20 mL) was added DIEA (1.20 g, 9.27 mmol, 1.99 equiv) dropwise at 0 °C under a nitrogen atmosphere. The resulting mixture was stirred overnight at room temperature. The reaction mixture was purified by reverse flash chromatography [column, C18 silica gel; mobile phase, acetonitrile in water (0.1% formic acid), 0% to 70% gradient in 10 min; detector, UV 254 nm] to give 2-[[2-(2,6-dioxopiperidin-3-yl)-1,3-dioxoisoindol-4-yl]oxy]-N-(2-[2-[2-(2-hydroxyethoxy)ethoxy]ethoxy]ethyl)acetamide (1.1 g, 46%) as a light yellow oil.

To a stirred mixture of 2-[[2-(2,6-dioxopiperidin-3-yl)-1,3-dioxoisoindol-4-yl]oxy]-N-(2-[2-[2-(2-hydroxyethoxy)ethoxy]ethoxy]ethyl)acetamide (1.60 g, 0.31 mmol, 1.00 equiv) and TEA (0.96 g, 0.95 mmol, 3.0 equiv) in DMF (16 mL) was added MsCl (0.55 g, 0.46 mmol, 1.51 equiv) dropwise at 0 °C. The resulting mixture was stirred for 12 h at room temperature. The residue was purified by reverse flash chromatography [column, C18 silica gel; mobile phase, acetonitrile in water (0.1% formic acid), 0% to 70% gradient in 40 min; detector, UV 254 nm] to give 2-(2-[2-[2-(2-[[2-(2,6-dioxopiperidin-3-yl)-1,3-dioxoisoindol-4-yl]oxy]acetamido)ethoxy]ethoxy]ethoxy)ethyl methanesulfonate (**S6-2 n = 1**, 840 mg, 45%) as a light yellow oil.

**General synthesis of R1-1c, R1-3c and R1-5c. Exemplified for R1-1c**. To a stirred mixture of **Intermediate X** (100 mg, 0.18 mmol, 1.00 equiv) and **S6-2 n = 1** (106 mg, 0.18 mmol, 1.00 equiv) in acetone (10 mL) were added K2CO3 (75 mg, 0.54 mmol, 3.0 equiv) in portions at room temperature under a nitrogen atmosphere. The resulting mixture was stirred overnight at room temperature under a nitrogen atmosphere. The resulting mixture was purified by reverse flash chromatography [column, C18 silica gel; mobile phase, acetonitrile in water (0.1% formic acid), 10% to 80% gradient in 30 min; detector, UV 254 nm]. The collected fraction was lyophilized to afford *tert*-butyl N-[(3S,4S)-8-(5-[2,3-dichloro-4-[2-(2-[2-[2-(2-[[2-(2,6-dioxopiperidin-3-yl)-1,3-dioxoisoindol-4-yl]oxy]acetamido)ethoxy]ethoxy]ethoxy)ethoxy]phenyl]-3-(hydroxymethyl)-6-methylpyrazin-2-yl)-3-methyl-2-oxa-8-azaspiro[4.5]decan-4-yl]carbamate (84 mg, 22%) as a white solid.

To a stirred mixture of *tert*-butyl N-[(3S,4S)-8-(5-[2,3-dichloro-4-[2-(2-[2-[2-(2-[[2-(2,6-dioxopiperidin-3-yl)-1,3-dioxoisoindol-4-yl]oxy]acetamido)ethoxy]ethoxy]ethoxy)ethoxy]phenyl]-3-(hydroxymethyl)-6-methylpyrazin-2-yl)-3-methyl-2-oxa-8-azaspiro[4.5]decan-4-yl]carbamate (40 mg, mmol, 1.00 equiv) in DCM (4 mL) were added TFA (0.80 mL) in portions at room temperature under a nitrogen atmosphere. The resulting mixture was stirred for 3 h at room temperature under a nitrogen atmosphere. The resulting mixture was concentrated and the residue was purified by reverse flash chromatography [column, C18 silica gel; mobile phase, acetonitrile in water (0.1% formic acid), 10% to 50% gradient in 30 min; detector, UV 254 nm]. The crude product (40 mg) was purified by Prep-HPLC [column: Xselect CSH OBD Column 30x150mm 5um, n; Mobile Phase A: water (0.1% formic acid), Mobile Phase B: acetonitrile; flow rate: 60 mL/min; Gradient: 20 B to 34 B in 7 min; 220 nm]. The collected fraction was lyophilized to afford **R1-1c** (21 mg, 56%) as a white solid.

### NMR spectroscopy

All ^1^H NMR spectra were acquired at 600 MHz at 25 °C and chemical shifts are represented in δ scale. Residual protium in the NMR solvent (MeOH-*d*4, δ 3.31) was used to reference chemical shifts. Data are represented as follows: assignment, chemical shift, integration, multiplicity (s, singlet; d, doublet; t, triplet; m, multiplet), and coupling constant in Hertz. All ^13^C NMR spectra were obtained at 150 MHz at 25 °C and chemical shifts are represented in δ scale. The carbon resonances of the NMR solvent (MeOH-*d*4, δ 49.15) are used to reference chemical shifts. Full assignment of protons and carbons were completed on the basis of the following two-dimensional NMR spectroscopy experiments: ^1^H-^1^H correlation spectroscopy (COSY), ^1^H-^13^C heteronuclear single quantum coherence non-uniform sampling (HSQC-NUS), ^1^H-^13^C heteronuclear multiple bond connectivity non-uniform sampling (HMBC-NUS) **(Tables S2-S9 and Figures S11-S50)**.

### LC-MS

Samples were separated over a Kinetex C18 column with water +0.1% formic acid (A) and acetonitrile +0.1% formic acid (B) as mobile phases. After an initial equilibration period 3 minutes at 5% B, a linear gradient to 100% was run over 13 minutes. The LC stream was inline with an Agilent 6530 qTOF with standard parameters **(Table S10 and Figures S51-S57)**.

### Antibodies and compounds

Antibodies used in this study were obtained commercially from the following sources: SHP2 (Bethyl, #A301-544A), Phospho-Thr202/Tyr204-Erk1/2 (CST, #9101), Cereblon (CST, #71810), β-actin (Millipore-Sigma, #A1978), β -tubulin (CST, #2146), GAPDH (CST, #5174). SHP099 and RMC-4550 were purchased from DC chemicals (Catalog No. DC9737) and ProbeChem Biochemicals (Catalog No. PC-35116), respectively. 6,8-Difluoro-4-Methylumbelliferyl Phosphate (DiFMUP, #D22065) and 6,8-difluoro-7-hydroxy-4-methylcoumarin (DiFMU, #D6566) were purchased from Life Technologies.

### Cell culture

The MV-4-11 cell line were purchased from the American Type Culture Collection (ATCC). The KYSE-520 and MOLM-13 cell lines were purchased from DSMZ (Braunschweig, Germany). Parental and CRBN knockout MOLT4 cells were a gift from Dr. Nathanael Gray (Dana-Farber Cancer Institute). All cell lines were cultured in RPMI-1640 media containing 10% FBS and 1% penicillin-streptomycin at 37°C with 5% CO2.

### Immunoblotting analysis

Cells were treated with PROTAC compounds as indicated and lysed in RIPA lysis buffer (25 mM Tris-HCl (pH 7.6), 150 mM NaCl, 1% Nonidet P-40, 1% Sodium deoxycholate, 0.1% Sodium dodecyl sulfate, 2 mM EDTA) with protease inhibitors (cOmplete Mini, Roche). Lysates were resolved on SDS-PAGE gels (Novex Tris-Glycine, ThermoFisher Scientific) and transferred to PVDF membranes using the XCell II wet blotting system (ThermoFisher Scientific). Membranes were blocked with 5% BSA diluted in TBST (Tris-buffered saline, 0.1% Tween 20) buffer and incubated with primary antibodies overnight. The blots were rinsed three times for 10 minutes each in 15 ml of TBST buffer and incubated with secondary antibodies diluted with 5% BSA for 1 h followed by three final washes. Specific protein bands were detected using SuperSignal™ West Pico PLUS Chemiluminescent Substrate (ThermoFisher Scientific).

### Measurement of inhibitory activity of compounds using DiFMUP

The phosphatase activities of SHP2-F285S and the isolated PTP domain were measured using the fluorogenic small molecule substrate, DiFMUP. Compounds were dissolved in DMSO at 10 mM concentration, diluted 1:10 in assay buffer (60 mM Hepes pH 7.2, 75 mM KCl, 75 mM NaCl, 1 mM EDTA, 0.05% Tween-20, and 2 mM DTT) and added to 96-well plates in a 3-fold dilution (concentration range 200 μM – 3.3 nM) in triplicate. Serially diluted compound was mixed with 0.5 nM SHP2-F285S mutant or isolated PTP domain proteins and incubated at room temperature for 1 h, after which 400 μM DiFMUP was added to each well. 1 h after DiFMUP addition, fluorescence was measured on a SpectraMax M5 plate reader (Molecular Devices) using excitation and emission wavelengths of 340nm and 450nm and the inhibitor dose−response curves were analyzed using non-linear regression curve fitting with control based normalization.

### Cell viability analysis

MV-4-11 cells were seeded at 1000 cells per ml of media in 10 cm plates in triplicate and treated with 100 nM R1-5C, 100 nM R1-1C, 100 nM RMC-4550 or DMSO carrier. The cells were replenished with compounds every 24 h and counted on days 1-9 using automated cell counter (Bio-Rad). KYSE-520 cells were plated onto 96-well plates (500 cells per well) in 100 μl medium and treated with 100 nM R1-5C, 100 nM RMC-4550 or DMSO carrier. The cells were replenished with compounds every 24 h and cell viability was assessed on days 1, 3, 5, 7, 9 by adding 20 μl CellTiter-Blue reagent (Promega) to wells and measuring luminescent signal using GloMax discover microplate reader (Promega).

### TMT-based LC-MS3 proteomics

MV-4-11 cells (10 M) were treated with DMSO carrier or 100 nM R1-5C, 100 nM R1-3C, 100 nM R1-1C, 100 nM RMC-4550, 1 μM pomalidomide in biological duplicate or singlicate for the indicated time periods. MOLT4 cells (5 M) were treated with 1 μM R1-5C for 5 h in biological singlicate. Cells were then harvested by centrifugation and cell pellets snap frozen in liquid Nitrogen. Sample preparation for TMT LC-MS3 mass spectrometry were performed as described previously^15^.

### Expression and purification of wildtype SHP2, F285S-SHP2 and isolated PTP domain

Human wild-type SHP2 (1-525 aa, UniProtKB: Q06124) was inserted into a modified pGEX6P1 vector with an N-terminal GST tag, followed by a PreScission cleavage site. Recombinant GST-SHP2 protein was overexpressed in *E.coli* BL21 (DE3) cells induced by 0.2 mM isopropyl-1-thio-D-galactopyranoside (IPTG) at 16°C overnight. Cells were harvested by centrifugation, resuspended in lysis buffer (25 mM Tris 7.5, 150 mM NaCl, 2 mM MgCl2, 2 mM TCEP and an EDTA-free protease inhibitor cocktail tablet (cOmplete, Roche)) and lysed by sonication. After centrifugation, the recombinant GST-SHP2 in supernatant was affinity-purified by Pierce glutathione agarose (Thermo Fisher Scientific) and eluted with lysis buffer containing 20 mM GSH. After GST tag cleavage with the recombinant HRV 3C protease, the SHP2 protein was further purified by HiTrap Heparin HP column (GE Healthcare). SHP2 containing fractions were pooled and finally polished over a Superdex200 10/300 GL size exclusion column (GE Healthcare) in buffer containing 25 mM Tris 7.5, 100 mM NaCl, 2 mM MgCl2, 2 mM TCEP. Protein sample concentration was determined based on the UV absorbance at 280 nm. Wildtype SHP2 protein was concentrated to 18 mg/ml for crystallization. SHP2-F285S and isolated PTP domain proteins were expressed and purified as described previously^8^.

### Crystallization and structure determination

The hanging-drop vapor diffusion method was used for co-crystallization of wild-type SHP2 in complex with RMC-4550. Wild-type SHP2 was incubated with RMC-4550 at a molar ratio of 1:2 and crystals were grown at 18°C by mixing equal volumes of the protein sample and reservoir solution (0.1 M Sodium Chloride, 0.1 M BIS-TRIS propane pH 8.5, 11% PEG 1500). Diffraction quality crystals were cryoprotected by supplementing reservoir solution with 20% glycerol and flash frozen in liquid nitrogen. X-Ray diffraction collection was performed at the Advanced Photon Source NE-CAT beamline 24 ID-E at 100 K using a wavelength of 0.979 Å. The diffraction images from single crystals were processed and scaled using XDS^16^. To obtain phases, molecular replacement for both copies of SHP2 in the unit cell was performed in Phenix with Phaser using chain B of a SHP2 crystal structure (PDB ID 5EHR) as the search model. Iterative model building was performed in COOT^17^. Reciprocal space refinement was performed in phenix.refine, using reciprocal space optimization of xyz coordinates, individual atomic B-factors, NCS restraints, optimization for X-ray/stereochemistry weights, and optimization for X-ray/ADP weights^18^. The RMC-4550 ligand coordinate and restraint file were generated using eLBOW^19^. In the final cycles of model building, NCS restraints were removed during refinement and overall model quality was assessed using MolProbity^20^. All crystallographic data processing, refinement, and analysis software was compiled and supported by the SBGrid Consortium^21^.

### Real-time quantitative PCR

1 million cells were treated with R1-5C at [add concentration] or DMSO for 2 or 16 hours. Cells were harvested after treatment in TRIzol (Thermo Fisher), and RNA was purified by phenol-chloroform extraction using MaXtract high density tubes (Qiagen). RNA was treated with Turbo DNase (Thermo Fisher) for 30 minutes at 37C, then re-extracted with chloroform isoamyl alcohol using MaXtract high density tubes. 1μg of RNA for each sample was used to make cDNA using the iScript cDNA synthesis kit (Bio-Rad). qPCR was performed using PowerUp SYBR Green Master Mix (Thermo Fisher) in a 10 μL total reaction with 0.25 μM forward and reverse primers, with two technical replicates per experiment. Primer sequences are shown in table XX Expression was normalized to the average of GAPDH and β-Actin expression. Significant differences were identified using Welch’s t-test in PRISM 9.0.2.

Primer sequence table:

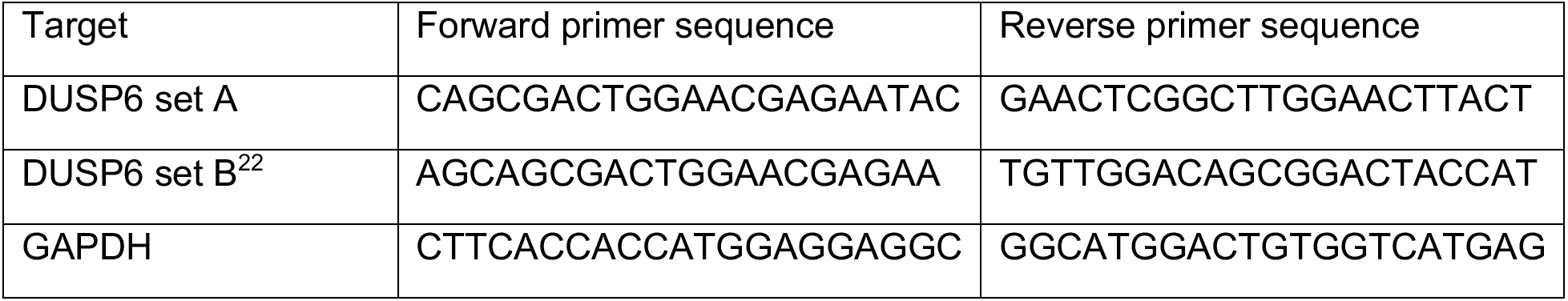
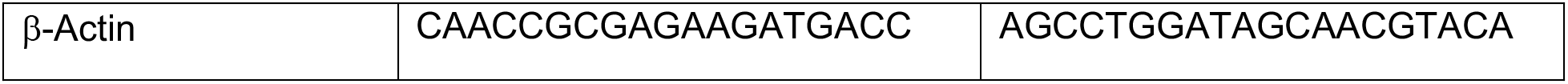

## Results

The CRBN E3 ligase ligands (pomalidomide, lenalidomide and thalidomide, collectively referred to as IMiDs) have been successfully employed in the design of a wide range of PROTACs to degrade various proteins ^23–26^. In order to develop an effective SHP2 PROTAC, therefore, we first designed a series of compounds designed to tether SHP099 to the IMiD pomalidomide. We chose pomalidomide as our E3 ligase ligand because it has higher cellular stability compared to other IMiDs ^27^.

To guide the sites for attachment of the linker to the SHP099 warhead, we relied on the X-ray structure of SHP2-SHP099 complex (PDB code 5EHR), which shows that there is an exit path for the linker when coupled at either of two positions on the dichloro-substituted aromatic ring **(Figure S1A)**. Two different linker strategies were therefore employed, using either aliphatic or ether attachments to the SHP099 warhead **(Figures S1B-E)**.

We tested whether any of these compounds catalyzed degradation of SHP2 in the MV4-11 acute myeloid leukemia cell line. When MV4-11 cells were treated with these compounds at doses ranging from 0.01 μM to 50 μM, there was no evidence of SHP2 degradation for any of the compounds, as judged by western blot **(Figure S2A)**.

Because none of these first-generation compounds resulted in SHP2 degradation, we determined whether the conjugation of pomalidomide to the SHP099 warhead affected the inhibitory potency of the parent SHP099 compound. We compared the inhibitory activity of SHP099 with the various SHP099-pomalidomide conjugates in enzymatic assays with purified protein, using the fluorogenic substrate DiFMUP and the weakly active SHP2 mutant F285S.

Upon titration of SHP099, the F285S enzyme showed a dose-dependent inhibition of phosphatase activity with a half maximal inhibitory concentration (IC50) of 62 nM, similar to our previous findings^28^ for its IC50, and similar to the KD values for wild-type SHP2 ^4^. In contrast, the inhibitory profiles of the SHP099-IMiD conjugates are right-shifted, with IC50 values ranging from 0.5 μM to 3.7 μM **(Figure S2B)**, indicating that the coupling of the linker to the warhead reduces inhibitory potency between roughly 10-100 fold, depending on the site of attachment and the nature of the linkage. In fact, simply installing a PEG chain on SHP099 caused a 10-fold decrease in inhibitory activity *in vitro* and in cells **(Figures S3A-S3C)**. Together, these findings revealed that coupling the linker to the warhead results in a minimum ∼10-fold reduction in potency, and suggested that an active PROTAC would require a more potent warhead to offset this consequence of linker coupling.

RMC-4550 is an allosteric inhibitor of SHP2 **(Figure 1A)** with a 50-fold higher potency than SHP099 *in vitro* ^12^. To determine whether a similar exit path for the linker exists for RMC-4550 bound to SHP2, we determined the structure of RMC-4550 bound to wild-type SHP2 by x-ray crystallography to 1.8 Å resolution **(Figure 1A**, **Table S1)**. The structure shows that the dichlorophenyl ring of RMC-4550 adopts virtually the identical pose as that of SHP099 when bound, exposing the same tethering sites to solvent for the design of PROTAC compounds **(Figure S4A,B)**.

**Figure 1.**
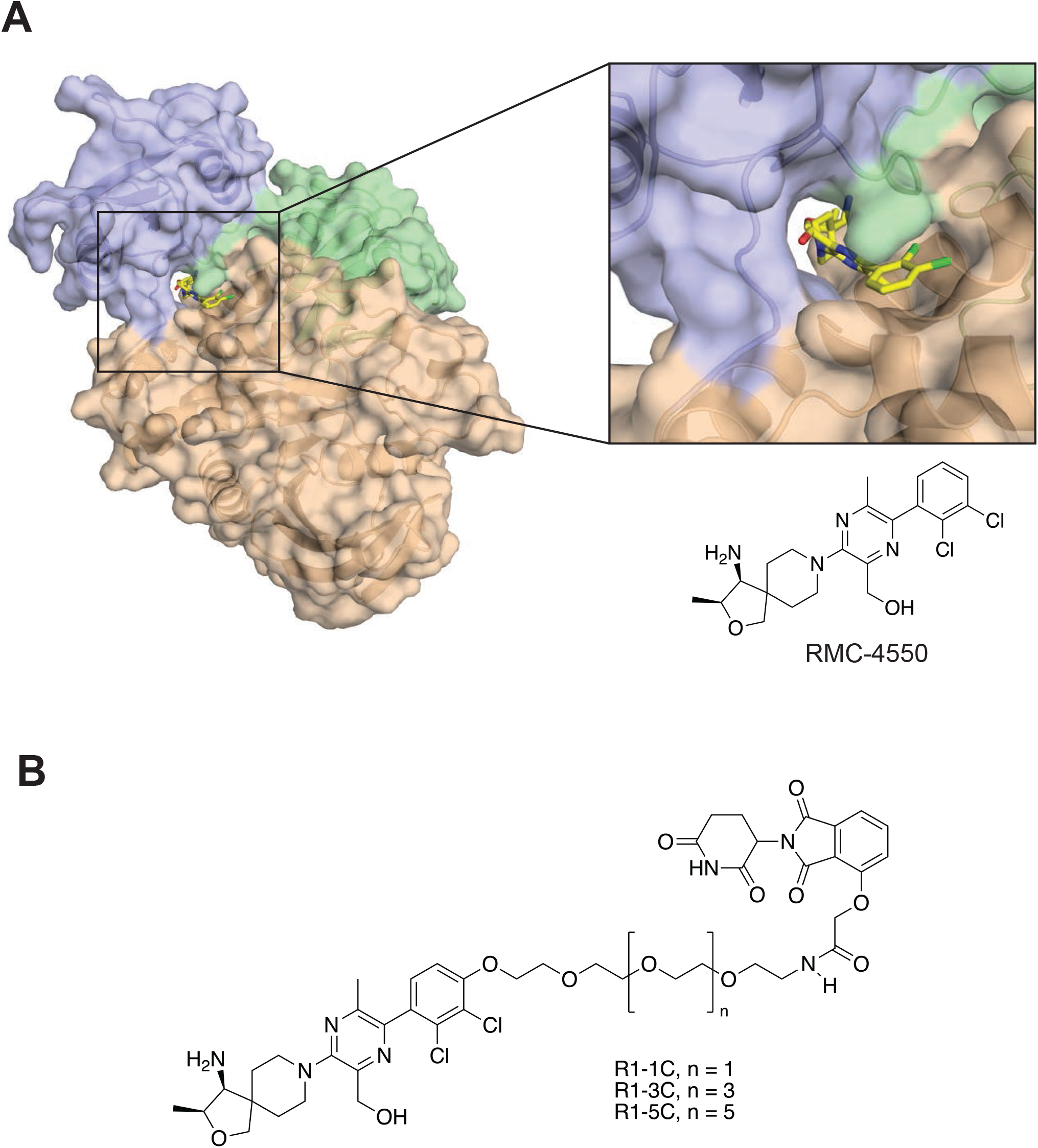
Structure of SHP2 protein in complex with RMC-4550. (A) Chemical structure of RMC-4550 and X-ray crystal structure of SHP2 in complex with RMC-4550 (PDB code XXX). Surface representation of SHP2 in complex with RMC-4550 bound in the central tunnel formed at the interface of N-SH2 (green), C-SH2 (blue) and PTP (wheat) domains. (B) Chemical structures of RMC-4550-based PROTAC candidates, R1-1C, R1-3C and R1-5C.

Since S9-3C is the most potent of all SHP099-IMiD conjugates, we substituted SHP099 with RMC-4550 in this design to create R1-3C **(Figure 1B)**. Upon titration of R1-3C, the SHP2 mutant F285S showed a dose-dependent inhibition of phosphatase activity with an IC50 of 14 nM **(Figure 2A)**, a roughly 10-fold reduction compared to RMC-4550 (IC50 = 1.5 nM). As expected, this compound displayed no inhibitory activity towards the free catalytic domain (PTP).

**Figure 2.**
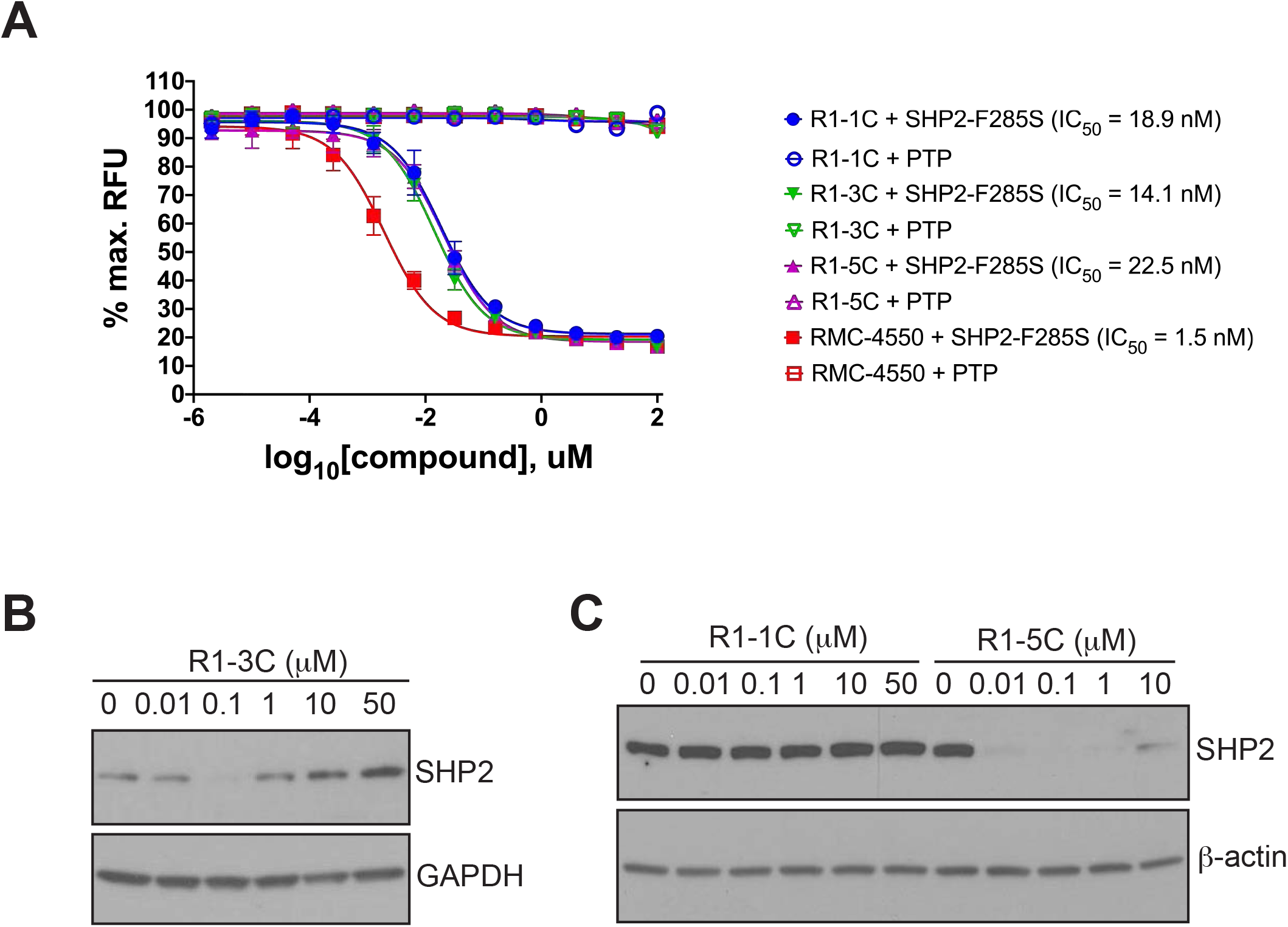
SHP2 degradation induced by RMC-4550-based PROTACs. (A) Inhibition of SHP2-F285S- or PTP-mediated DIFMUP dephosphorylation by R1-1C, R1-3C, R1-5C and RMC-4550. MV4;11 cells were treated with increasing doses of R1-3C (B), R1-1C or R1-5C (C) for 24 h and subjected to Western blotting using SHP2, GAPDH and β-actin antibodies.

We treated MV4;11 cells with increasing doses of R1-3C or with DMSO carrier as a control for 24 h and measured SHP2 protein levels by Western blotting. We observed a dose-dependent decrease in SHP2 levels with maximal degradation at 100 nM **(Figure 2B)**. The loss of activity observed at higher R1-3C concentrations (1 µM, 10 µM and 50 µM), referred to as a hook effect, is a signature trait of PROTACs.

Because the length of the linker plays a key role in the potency of PROTACs ^29–32^, we varied the linker lengths between RMC-4550 and pomalidomide to search for a PROTAC with higher potency. Extension of the linker with two additional PEG units **(R1-5C in Figures 1B, 2C, S5A)** resulted in a PROTAC with greater potency, whereas shortening the linker by two PEG units **(R1-1C in Figures 1B, 2C)** caused complete loss of PROTAC activity.

To determine the kinetics of R1-5C mediated degradation of SHP2 in MV4-11 cells, we assessed SHP2 protein abundance at a series of time points after compound addition. SHP2 levels are substantially reduced within 6 h after R1-5C treatment, reaching maximal depletion after 16 h **(Figure 3A).** SHP2 remains depleted at 24 h; however, it reaccumulates to pre-treatment levels by 48h **(Figure S5B)**, consistent with cellular half-lives observed for other PROTACs ^23^.

**Figure 3.**
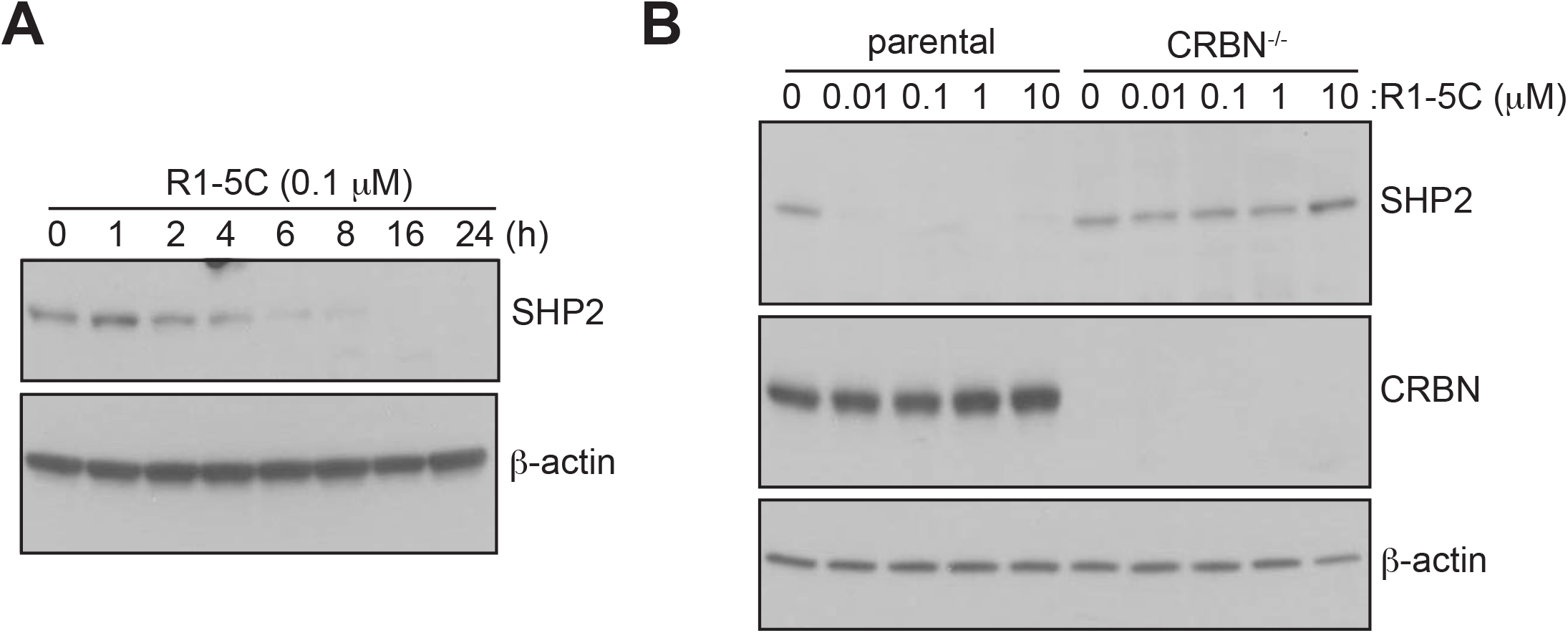
Cell-based evaluation of R1-5C. (A) Time course of SHP2 degradation by R1-5C (100 nM) in MV4;11 cells. Immunoblotting with SHP2 and β-actin antibodies. (B) CRBN^-/-^ and parental MOLT4 cells were treated with increasing doses of R1-5C for 24 h and subjected to Western blotting using SHP2, CRBN and β-actin antibodies

We confirmed the dependence of SHP2 depletion on the CRBN E3 ligase by comparing the degradation activity of our PROTAC compounds in parental MOLT4 and CRBN^-/-^ knockout cells. Whereas PROTAC treatment reduced SHP2 protein abundance in MOLT4 parental cells, PROTAC-dependent degradation of SHP2 was not detectable in CRBN^-/-^ cells, confirming the CRBN requirement for PROTAC activity **(Figures 3B, S5C)**.

We then performed time-resolved quantitative proteomics to further evaluate the selectivity of these compounds for SHP2 and to determine the kinetics of SHP2 degradation in MV4;11 cells. The cells were treated with R1-5C (100 nM for 2, 4, 8, or 16 h) R1-3C (100 nM for 4 or 16 h), R1-1C (100 nM for 4 or 16 h), RMC-4550 (R1-1C (100 nM for for 16 h), pomalidomide (1 µM for 5 h) or a vehicle control (DMSO) and protein abundance was measured quantiatively using 16-plex tandem mass tag (TMT) isobaric labels, as described previously ^15^. R1-5C exhibits striking specificity for degradation of SHP2, which is evident within 4 h **(Figure 4)**. At 16 h many of the secondary effects of SHP2 depletion recapitulate the consequence of allosteric inhibition **(proteins labeled in blue in Figures 4C, 4D, highlighted with yellow dots in Figure S6)**, providing further evidence of the SHP2 specificity of the R1-5C PROTAC. By comparison, R1-3C, which has a shorter linker, resulted in more moderate SHP2 depletion at 16 h **(Figure S7A)**, and R1-1C did not induce SHP2 degradation even at 16 h, as anticipated **(Figure S7B)**. Pomalidomide-induced degradation of classical IMiD targets (IKZF1 and ZFP91) serves as a positive control and validates the authenticity of the proteomics experiment **(Figures 4E and S7C)**. Quantitative proteomics of MOLT4 cells treated with R1-5C compared to DMSO control after 5 h also showed that SHP2 is the only protein significantly reduced in abundance in MOLT4 cells **(Figure S8)**.

**Figure 4.**
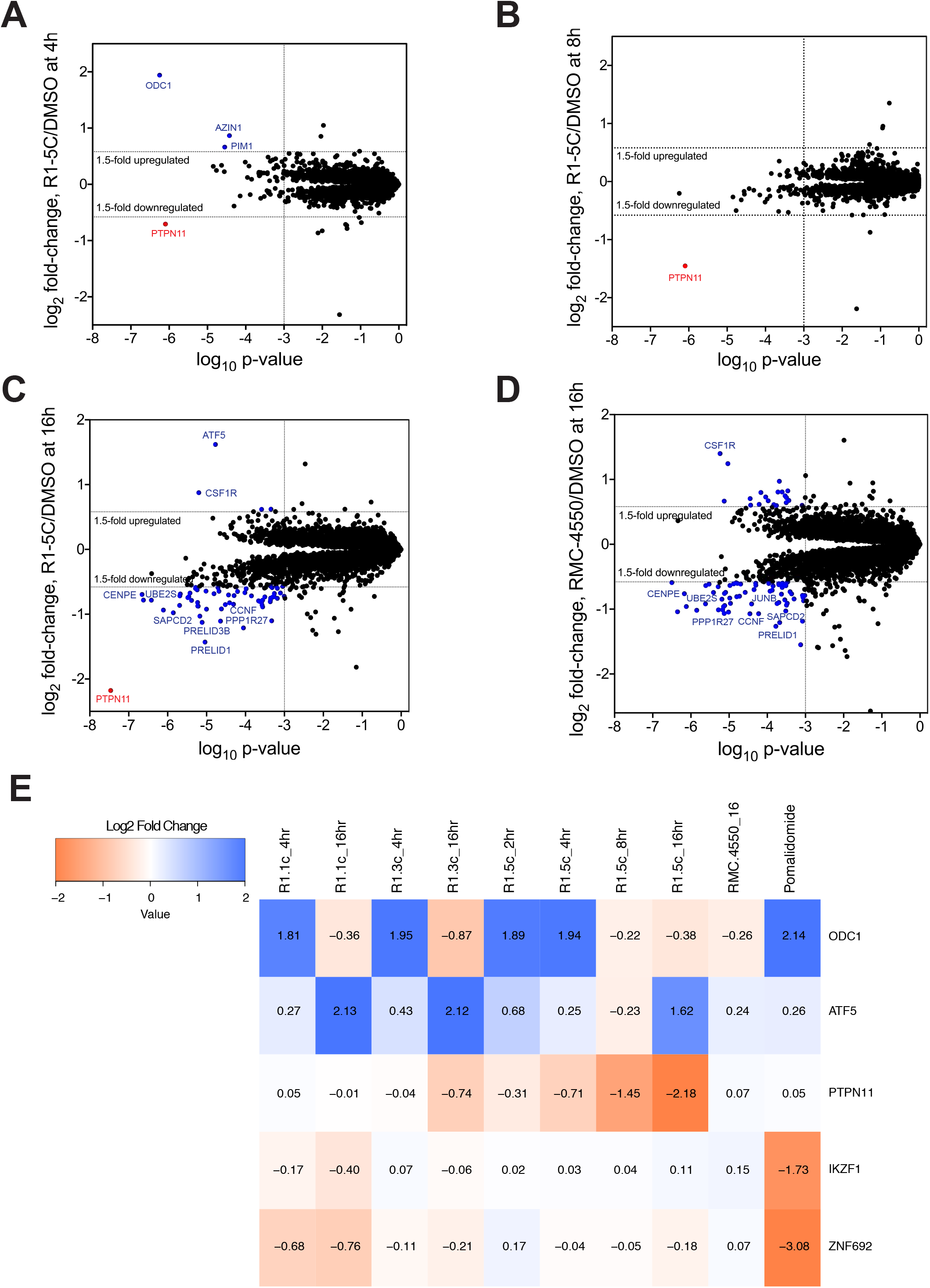
Proteomics analysis showing selective SHP2 degradation by R1-5C. (A-D) Scatterplots displaying relative fold-change in SHP2 abundance following treatment of MV4;11 cells with 100 nM R1-5C for 4 h (A), 8 h (B), 16 h (C) or 100 nM RMC-4550 (D). SHP2/PTPN11 is highlighted in red. Hits highlighted in blue in (C) and (D) indicate changes in abundance of proteins at 16 h time point due to secondary effects (such as transcriptional responses) of SHP2 degradation or inhibition. (E) Heatmap of the protein abundance changes in MV4;11 cells comparing treatment with 100 nM R1-1C (4 h and 16 h), 100 nM R1-3C (4 h and 16 h), 100 nM R1-5C (2 h, 4 h, 8 h and 16 h), 100 nM RMC-4550 (16 h) and 1 μM pomalidomide (5 h). The heatmap colors are scaled with red indicating a decrease in protein abundance (−2 log2 FC) and blue indicating an increase (2 log2 FC) in protein abundance.

To assess the time-dependent recovery of SHP2 protein abundance, we treated MV4-11 cells with 100 nM R1-5C for 24 h, washed out the compound, and lysed cells at various time points after washout for immunoblotting. SHP2 protein abundance recovered to basal (DMSO-treated) levels 24 h after washout **(Figure S9)**.

We also examined whether R1-5C affects MAPK signaling. We treated KYSE-520 cells with R1-5C or DMSO carrier and monitored the levels of DUSP6 transcript, a commonly used pharmacodynamic marker for MAPK pathway activity downstream of SHP2 ^4^. Cells treated with R1-5C showed significant downregulation of DUSP6 mRNA amounts **(Figures 5A and S10)**. Suppression of DUSP6 transcript abundance is observed both at 2 h and 16 h after treatment; the reduction at the late time point reports on the effect of SHP2 degradation on DUSP6 mRNA amounts, whereas the reduction at the 2 h time point indicates that allosteric inhibition of SHP2 by the RMC-4550 warhead is ongoing prior to SHP2 depletion. The inhibition of cancer cells by R1-5C was also assessed in a cell proliferation assay, which showed that R1-5C significantly inhibits KYSE-520 and MV4;11 cell growth, with an inhibitory effect comparable to that of RMC-4550 **(Figures 5B and 5C)**.

**Figure 5.**
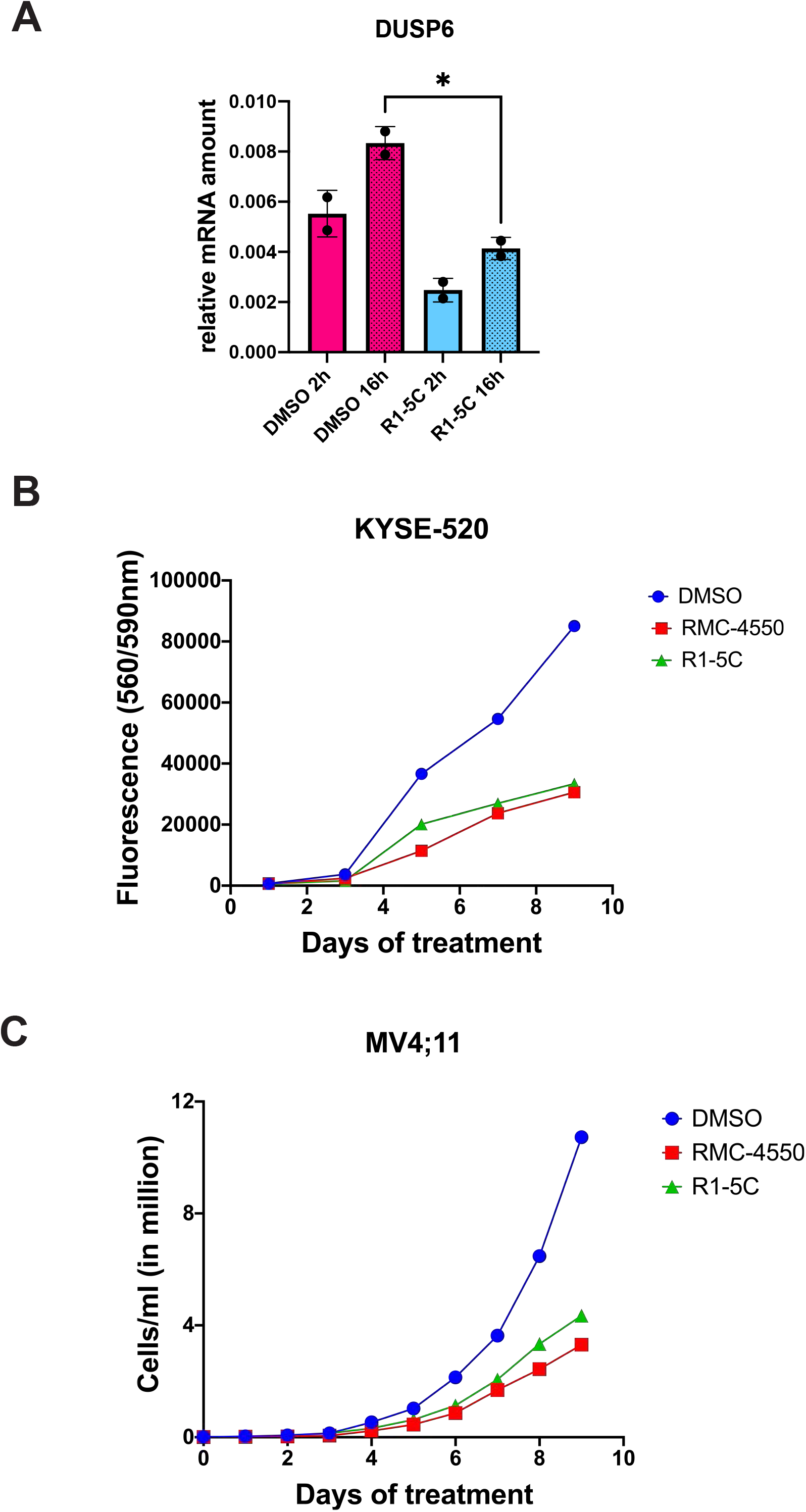
R1-5C inhibits MAPK signaling and suppresses cancer cell growth. (A) Downregulation of DUSP6 transcript levels in KYSE-520 cells after 2 h and 16 h treatment with 200 nM R1-5C or DMSO carrier. DUSP6 mRNA was quantified by RT-qPCR using primer set B. (B and C) Cells were treated with 100 nM R1-5C, 100 nM RMC-4550 or DMSO carrier in triplicate and cell numbers were assessed at the indicated time points by automated cell counting (for MV4;11 cells) or CellTiter-Blue assay (for KYSE-520 cells).

## Discussion

We report here the design and evaluation of R1-5C, a potent and highly selective SHP2 PROTAC featuring an SHP2 allosteric site-binding warhead and the CRBN-targeting IMiD pomalidomide. R1-5C expands the range of PROTACs targeting SHP2, which currently include a VHL-targeting PROTAC ^33^ and the Novartis clinical candidate TNO155 coupled to thalidomide^34^.

Key features required for the degradative activity of our designed compound include the warhead coupling site, the linker chemistry, and the linker length. The high degree of selectivity of R1-5C for SHP2 has been documented in this study using stringent whole proteome analysis in two different cell lines, MV4;11 **(Figure 4)** and MOLT4 **(Figure S8)**. In both of these lines, SHP2 (gene name PTPN11) is the only protein that shows a statistically significant reduction in abundance at time points from 4-8 hours, before secondary effects of SHP2 inhibition can be observed. By comparison, the selectivity of other reported SHP2-targeting PROTACS has not yet been assessed using whole proteomic studies ^33, 34^.

Whereas some IMiD-dependent PROTACs can induce detectable degradation of target proteins within 0.5 h, several hours of treatment with R1-5C are required before significant depletion of SHP2 is observed in either cell line. Although the origin of the slower onset of SHP2 degradation is not clear, the other recently reported, SHP2-targeting PROTACs also exhibit similar degradation kinetics based on Western blot analysis, independent of whether degradation is mediated by VHL ^33^ or CRBN ^34^.

R1-5C and related compounds are poised to serve as useful tools to investigate physiological roles of SHP2. An important question to be addressed in future work is whether R1-5C can be optimized to power degradation of oncogenic, mutant forms of SHP2. Such SHP2 PROTACs would then extend compound efficacy beyond RTK and ERK-dependent cancers that harbor wild-type SHP2 to cancers with mutant SHP2 and to patients with human genetic disorders like Noonan and LEOPARD syndromes.

## Acknowledgments

We thank Jon Aster members of the Blacklow laboratory for helpful discussions.

## Data availability

Diffraction data and refined coordinates have been deposited in the Protein Data Bank under accession code XXX.

---

**Figure S1.**
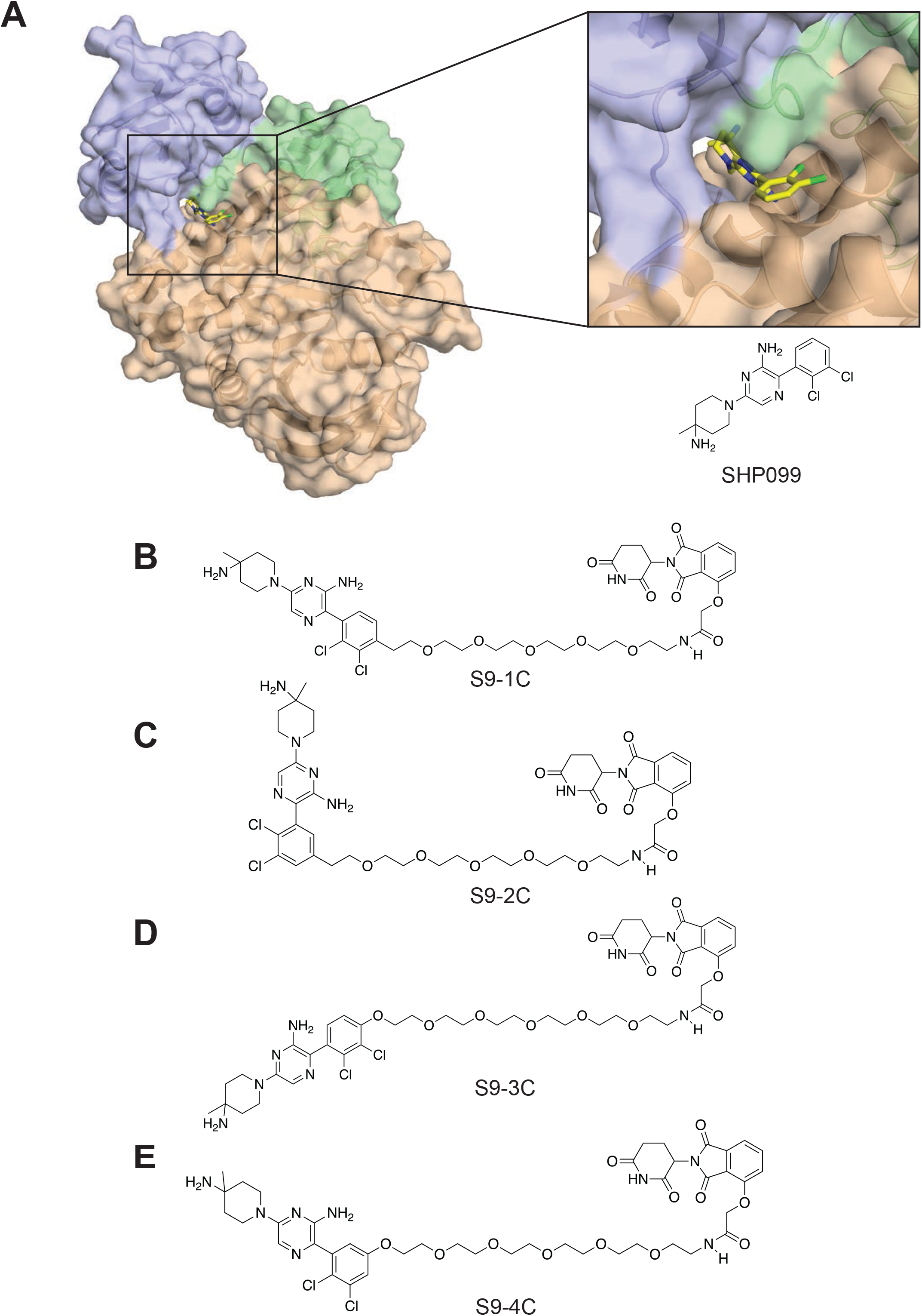
Structure of SHP2 protein in complex with SHP099. (A) Chemical structure of SHP099 and X-ray crystal structure of SHP2 in complex with SHP099 (PDB code 5EHR). Surface representation of SHP2 in complex with SHP099 bound in the central tunnel formed at the interface of N-SH2 (green), C-SH2 (blue) and PTP (wheat) domains. (B-E) Chemical structures of SHP099-IMiD conjugates, S9-1C, S9-2C, S9-3C and S9-4C.

**Figure S2:**
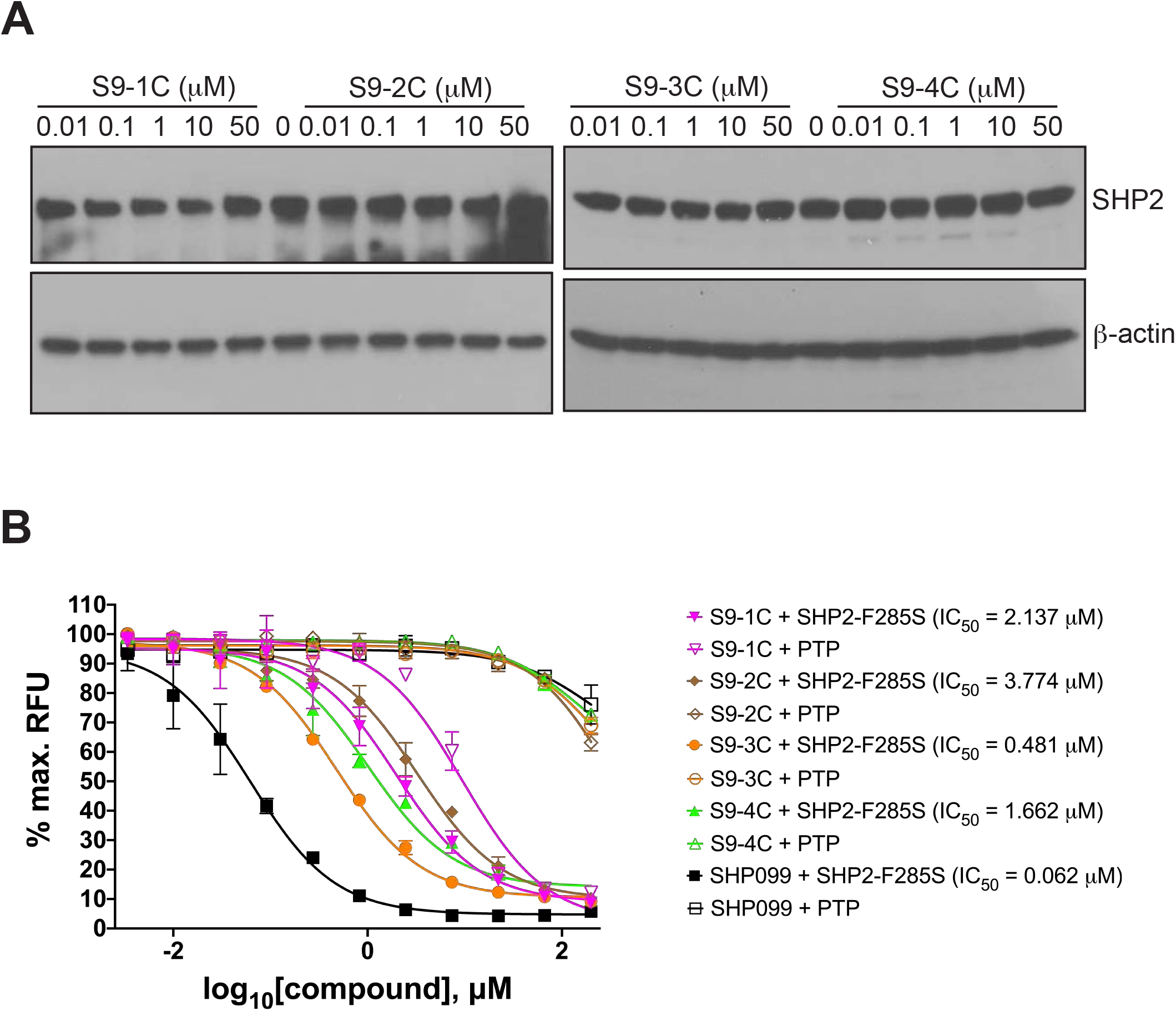
E**v**aluation **of SHP099-IMiD conjugates.** (A) MV4;11 cells were treated with increasing doses of S9-1C, S9-2C, S9-3C and S9-4C for 24 h and subjected to Western blotting using SHP2 and β-actin antibodies. (B) Inhibition of SHP2(F285S)- or PTP-catalyzed DIFMUP dephosphorylation by SHP099 and SHP099-IMiD conjugates.

**Figure S3.**
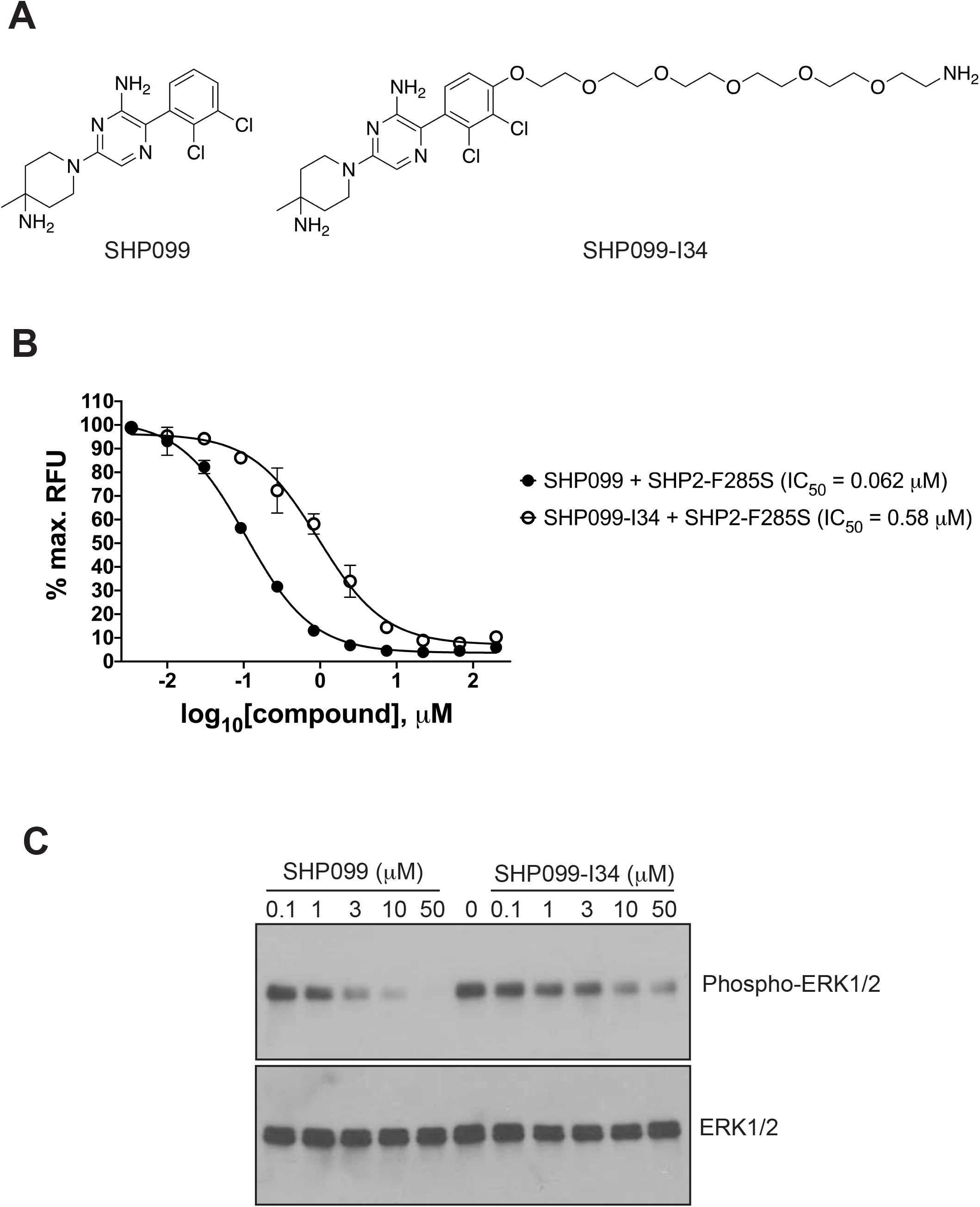
Evaluation of SHP099-PEG6 intermediate compound. (A) Chemical structures of SHP099 and SHP099-PEG6 intermediate compound (SHP099-I34). (B) Inhibition of SHP2-F285S-mediated DIFMUP dephosphorylation by SHP099 and SHP099-I34. (C) MV4;11 cells were treated with increasing doses of SHP099 and SHP099-I34 for 24 h and subjected to Western blotting using phospho-ERK1/2 and ERK1/2 antibodies.

**Figure S4.**
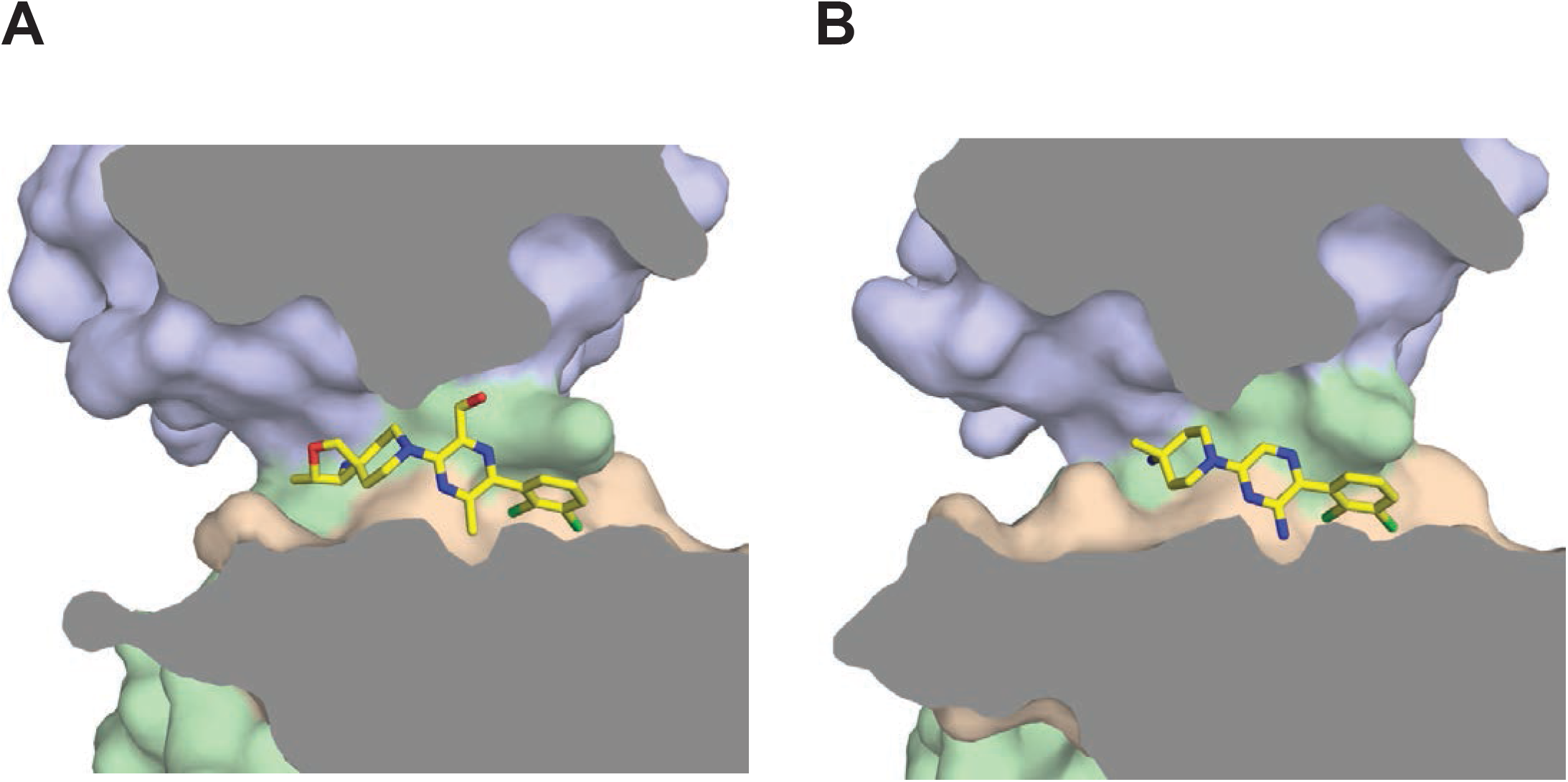
Allosteric compound SHP2 binding pocket. A cross-section view of the SHP2 binding pocket with (A) RMC-4550 and (B) SHP099 reveals the extent of solvent exposure to the dichlorophenyl ring for both compounds. Surface representation of SHP2 is colored as in figure 1 and interior colored in grey. The bound RMC-4550 and SHP099 compounds are shown in yellow sticks.

**Figure S5.**
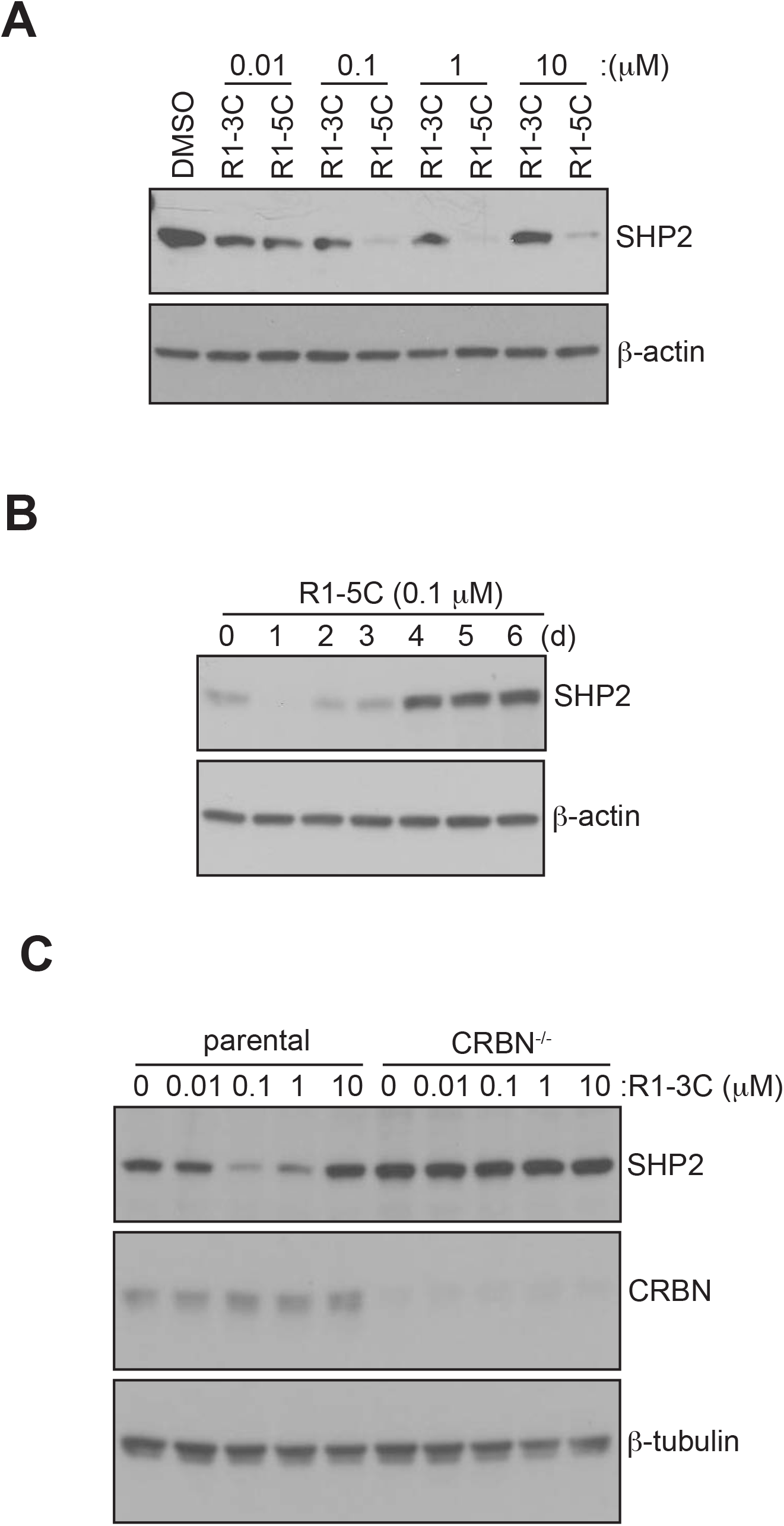
Cellular evaluation of RMC-4550-based PROTAC candidates. (A) MV4;11 cells were treated with DMSO carrier or increasing doses of R1-3C and R1-5C for 24 h and subjected to Western blotting using SHP2 and β-actin antibodies. (B) MV4;11 cells were treated with 100 nM R1-5C and cells were harvested after 1, 2, 3, 4, 5 or 6 days and subjected to immunoblotting using SHP2 and β-actin antibodies. (C) CRBN^-/-^ and parental MOLT4 cells were treated with increasing doses of R1-3C for 24 h and subjected to Western blotting using SHP2, CRBN and β-tubulin antibodies.

**Figure S6.**
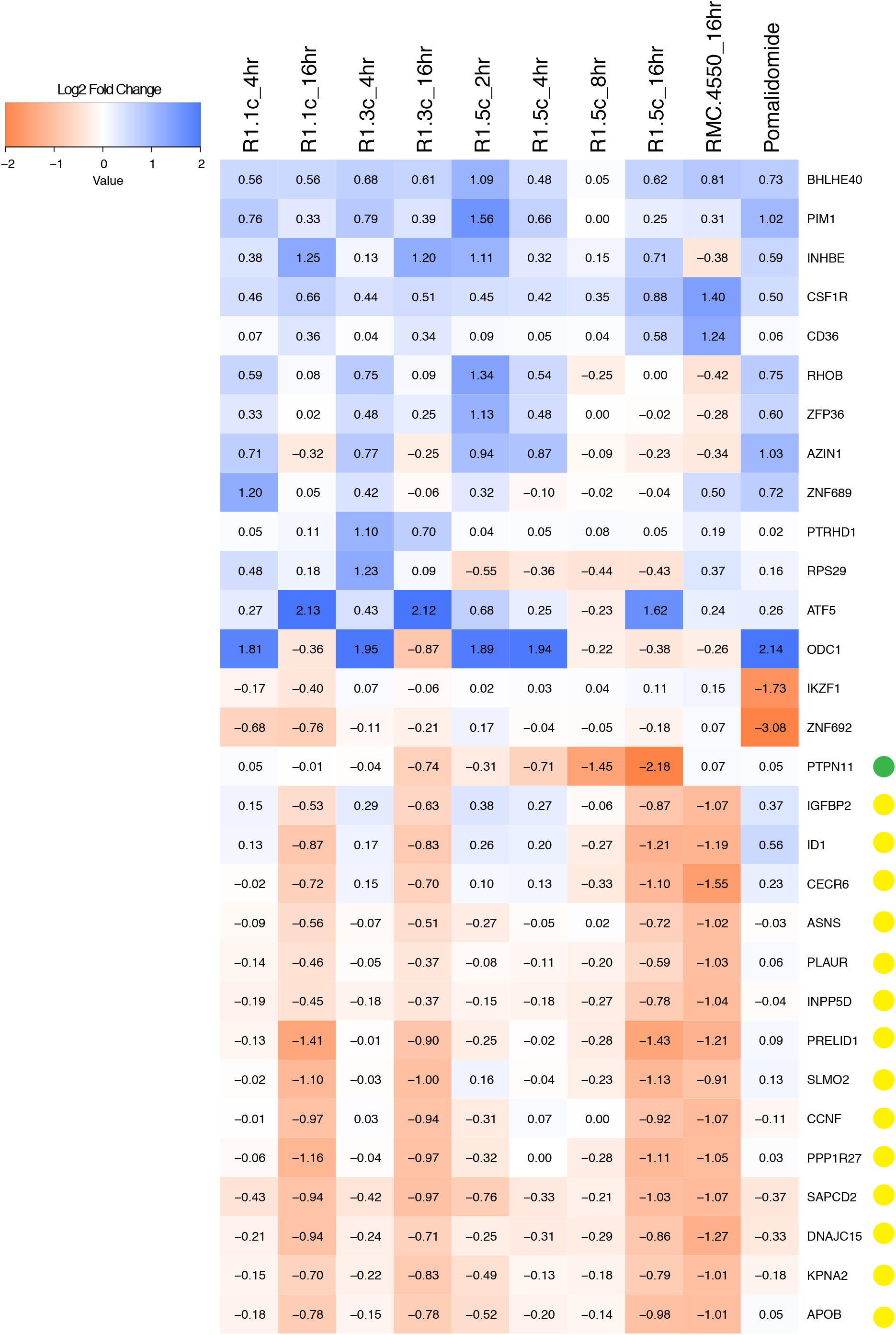
Proteome-wide changes caused by SHP2 PROTAC compounds. Heatmap of the protein abundance changes in MV4;11 cells comparing treatment with R1-1C, R1-3C, R1-5C, RMC-4550 and pomalidomide. The heatmap colors are scaled with red indicating a decrease in protein abundance (−2 log2 FC) and blue indicating an increase (2 log2 FC) in protein abundance. The green dot marks fold-changes in SHP2 protein. The yellow dots represent changes in other proteins due to secondary effects (such as transcriptional responses) of SHP2 degradation or SHP2 inhibition at 16 h time points.

**Figure S7.**
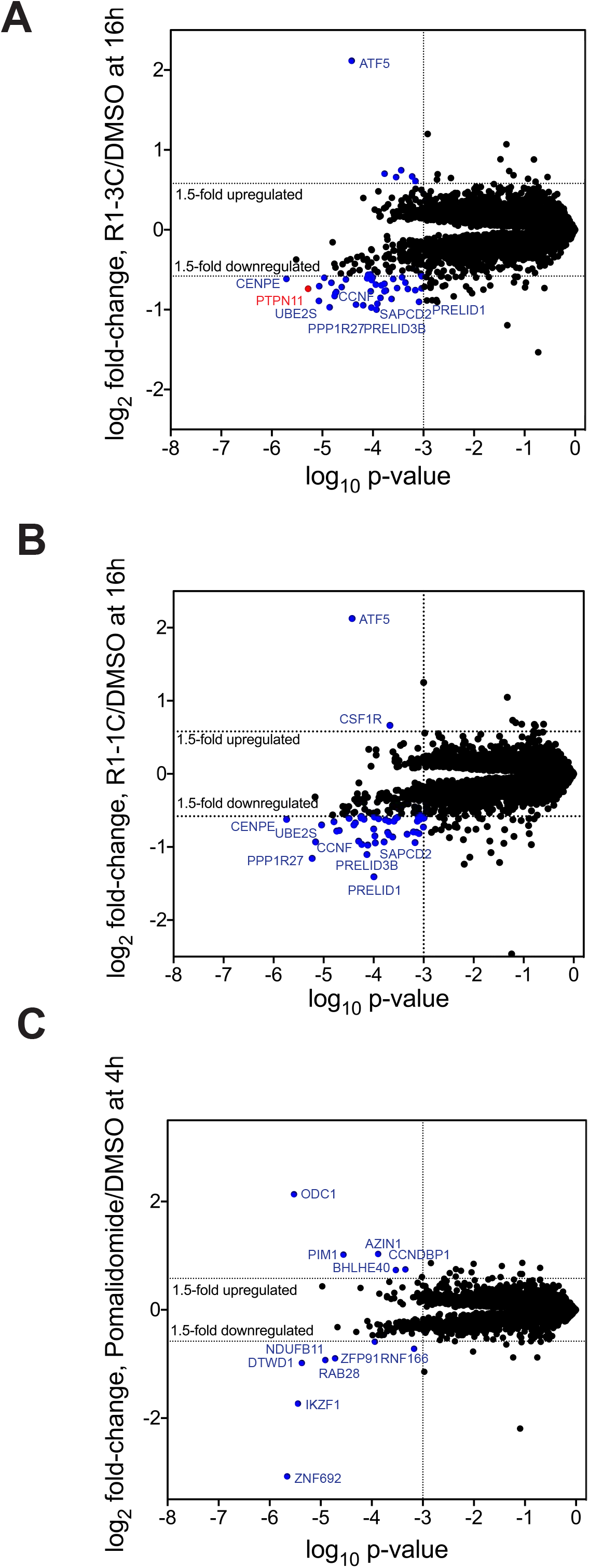
Proteomic profiling of R1-3C, R1-1C and pomalidomide. Scatterplots depicting relative fold-changes in protein abundances following treatment of MV4;11 cells with 100 nM R1-3C for 16 h (A), 100 nM R1-1C for 16 h (B) or 1 μM pomalidomide for 5 h (C). SHP2/PTPN11 is highlighted in red in (A). Hits highlighted in blue in (A) and (B) indicate changes in abundance of proteins at 16 h time point due to secondary effects (such as transcriptional responses) of SHP2 degradation or inhibition.

**Figure S8.**
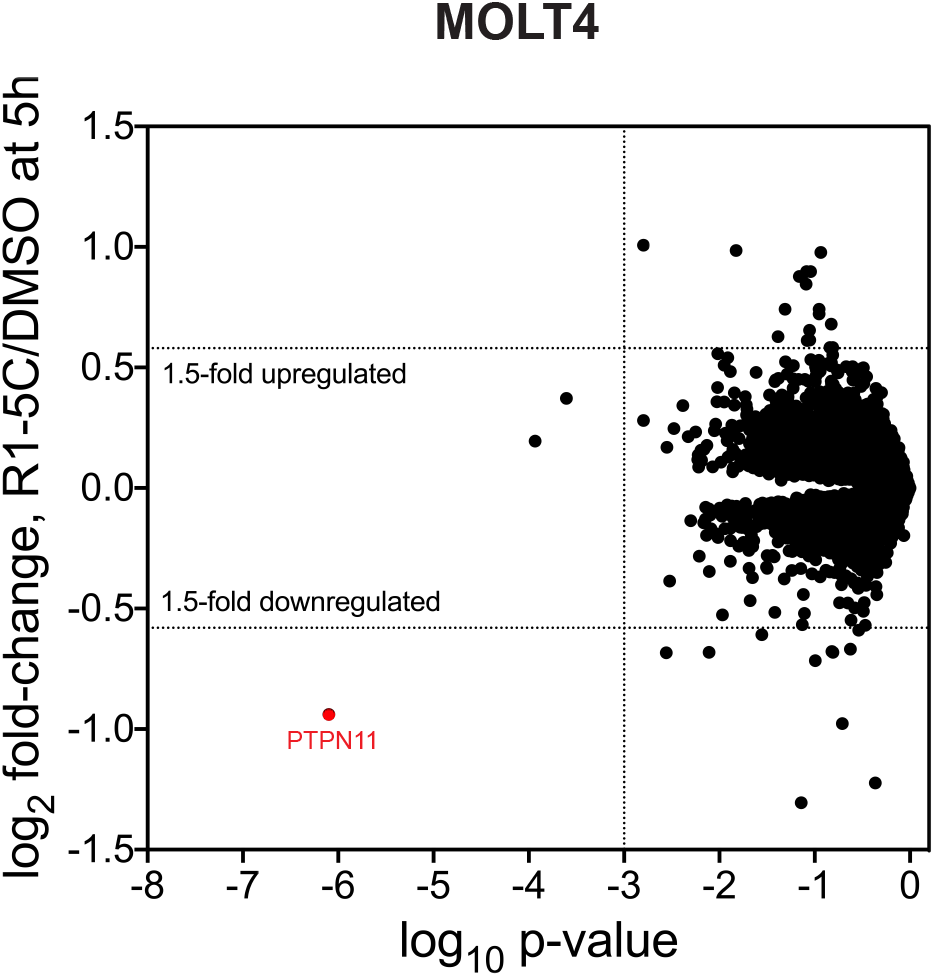
Selective SHP2 degradation by R1-5C in MOLT4 cells. MOLT4 cells were treated with 1 μM R1-5C for 5 h and subjected to TMT-based proteomics. Scatterplot depicting relative fold-change in SHP2 abundance upon R1-5C treatment. SHP2/PTPN11 is highlighted in red.

**Figure S9.**
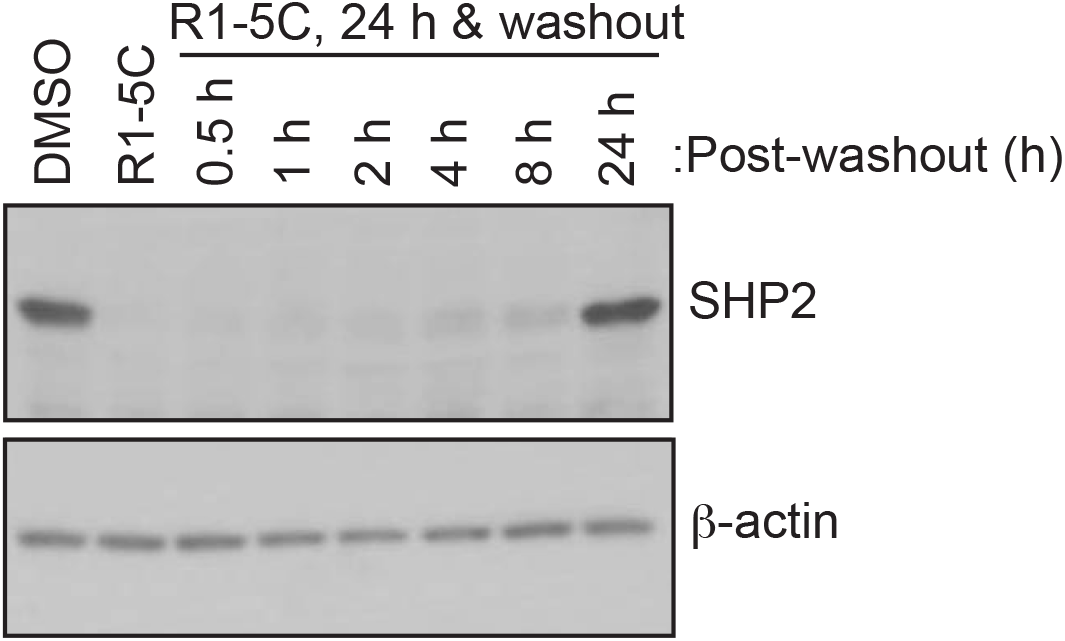
SHP2 recovery after R1-5C washout. MV4;11 cells were treated with 100 nM R1-5C or DMSO carrier for 24h, followed by compound washout and incubation in media. Cells were harvested after 0.5, 1, 2, 4, 8 or 24 h post-washout and SHP2 protein levels were assessed by immunoblotting.

**Figure S10.**
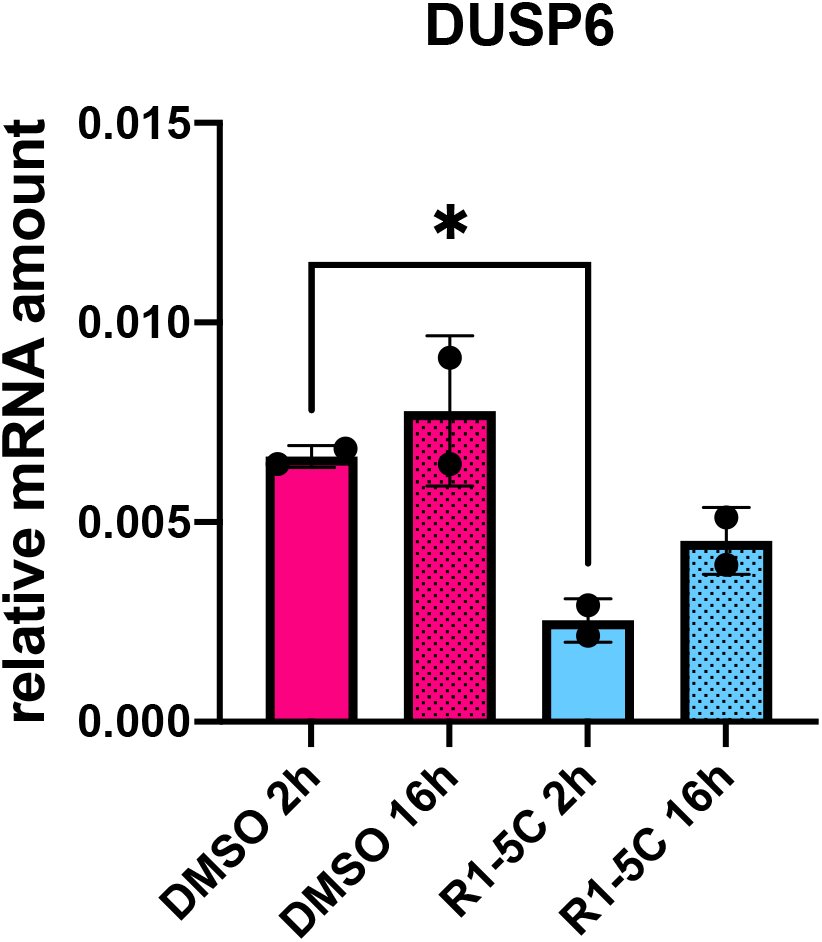
R1-5C inhibits MAPK signaling. Downregulation of DUSP6 transcript levels in KYSE-520 cells after 2 h and 16 h treatment with 200 nM R1-5C or DMSO carrier. DUSP6 mRNA was quantified by RT-qPCR using primer set A.

**Table S1:**
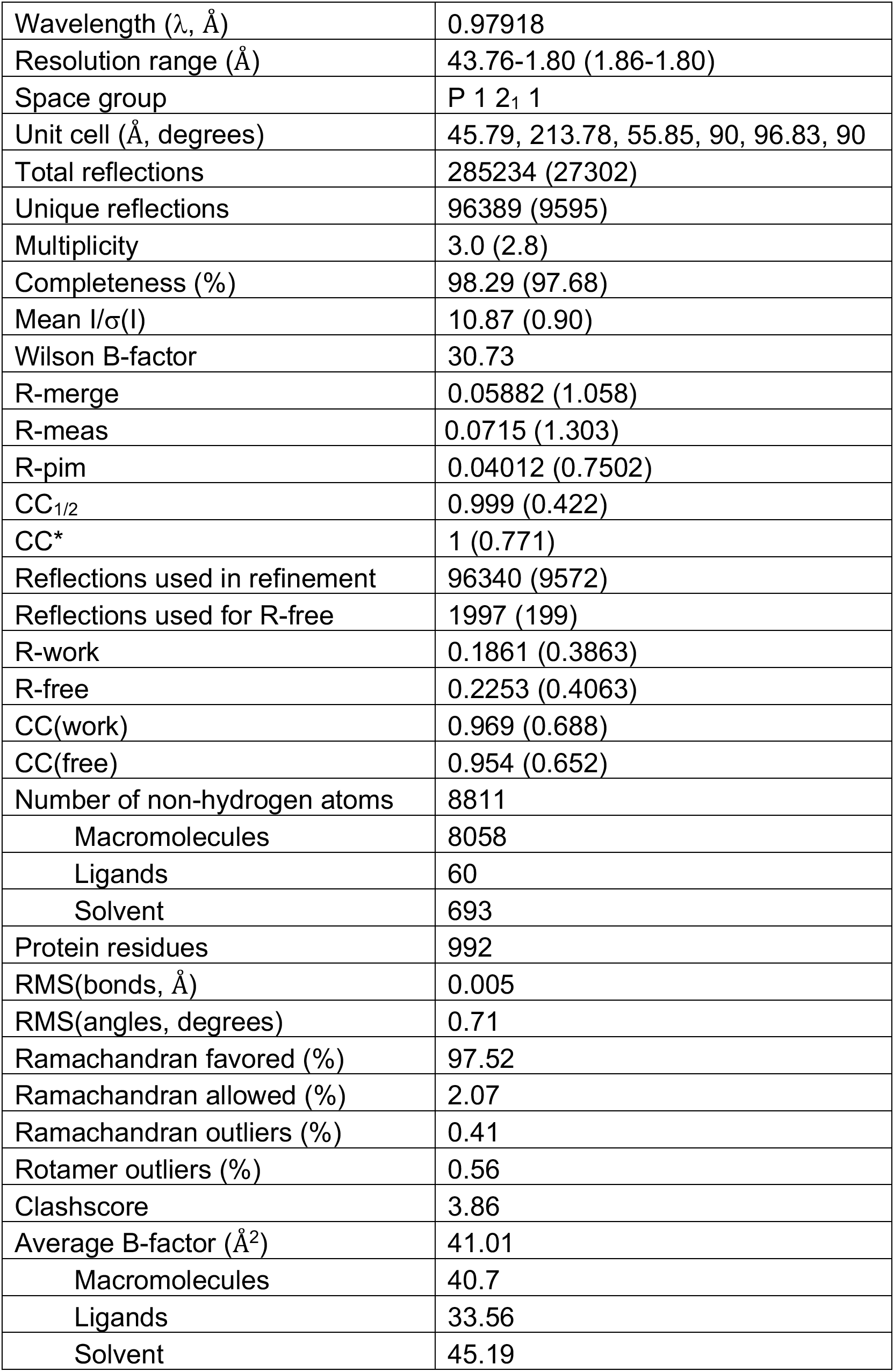
X-ray crystallography data collection and refinement statistics.

**Figure S11.**
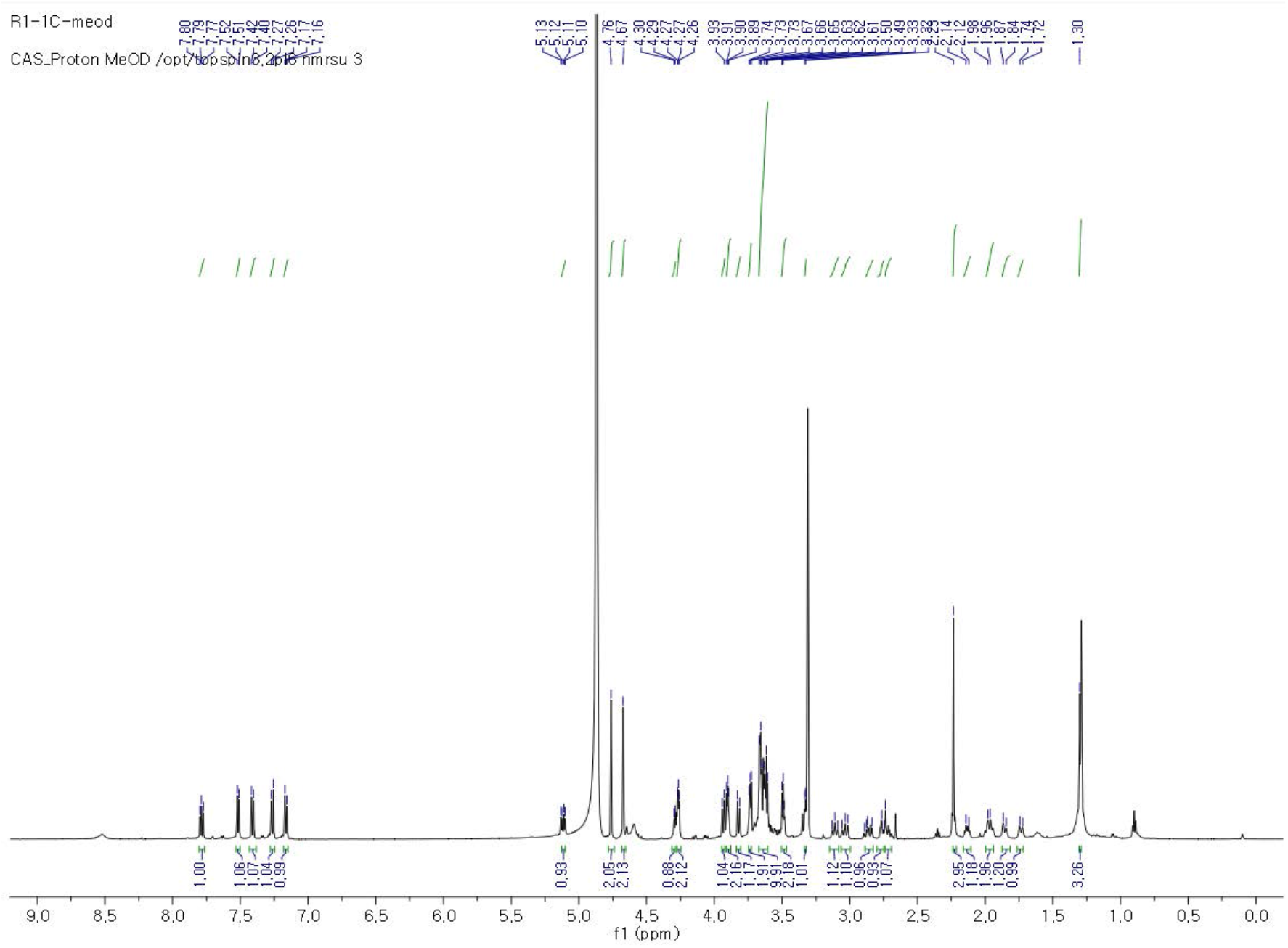
^1^H NMR spectrum of R1-1C in MeOH-*d*4.

**Figure S12.**
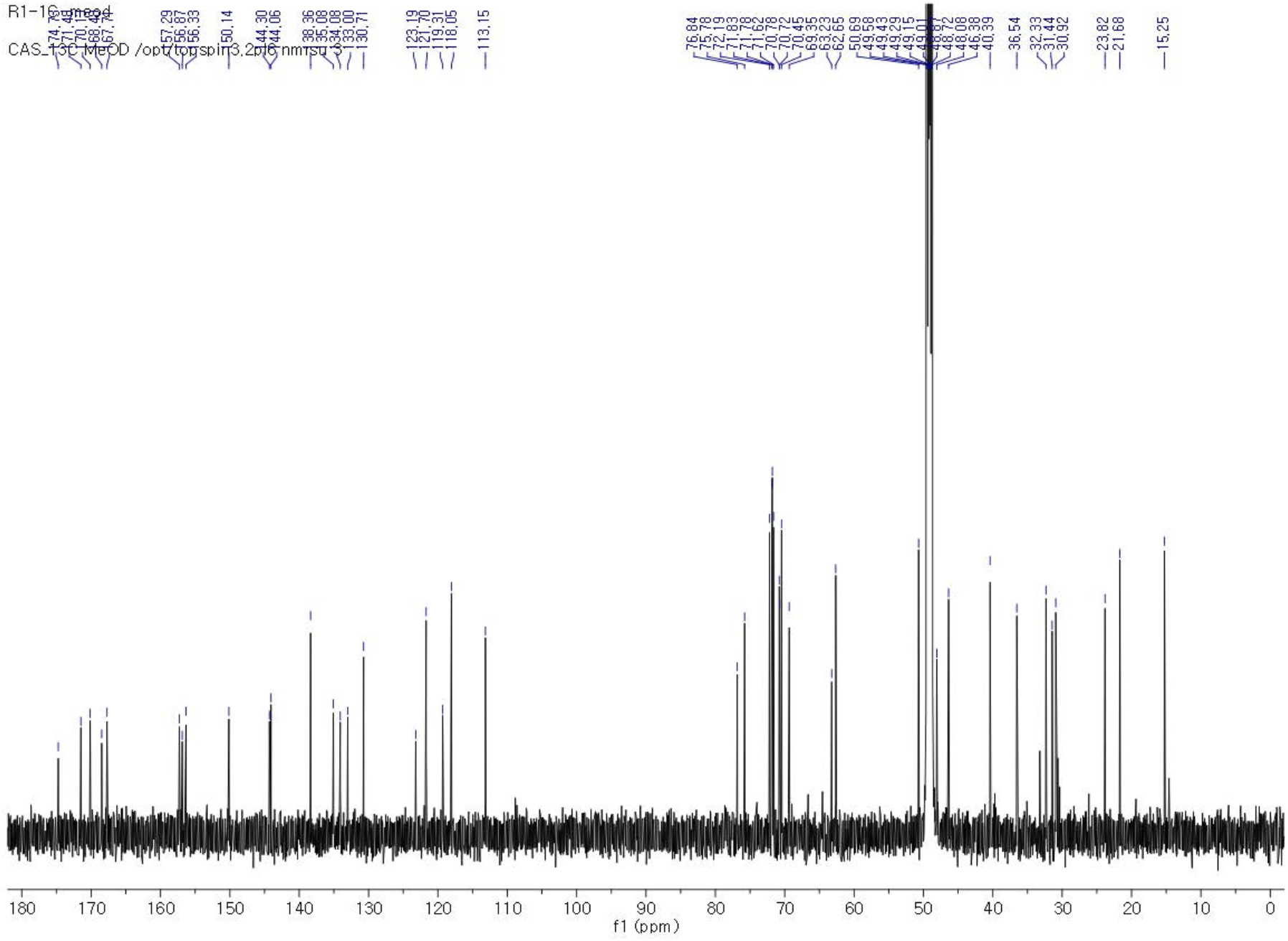
^13^C NMR spectrum of R1-1C in MeOH-*d*4.

**Figure S13.**
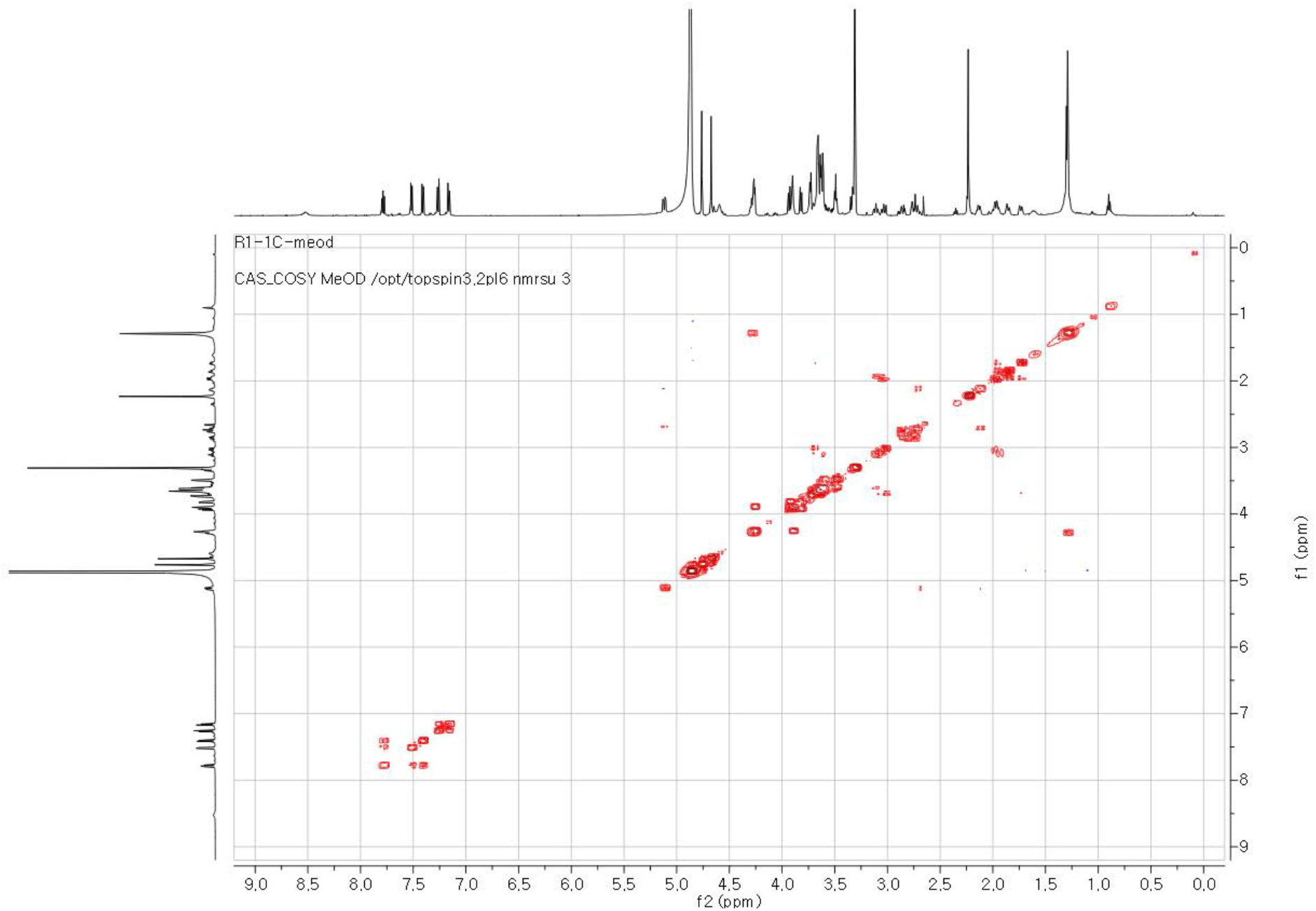
COSY NMR spectrum of R1-1C in MeOH-*d*4.

**Figure S14.**
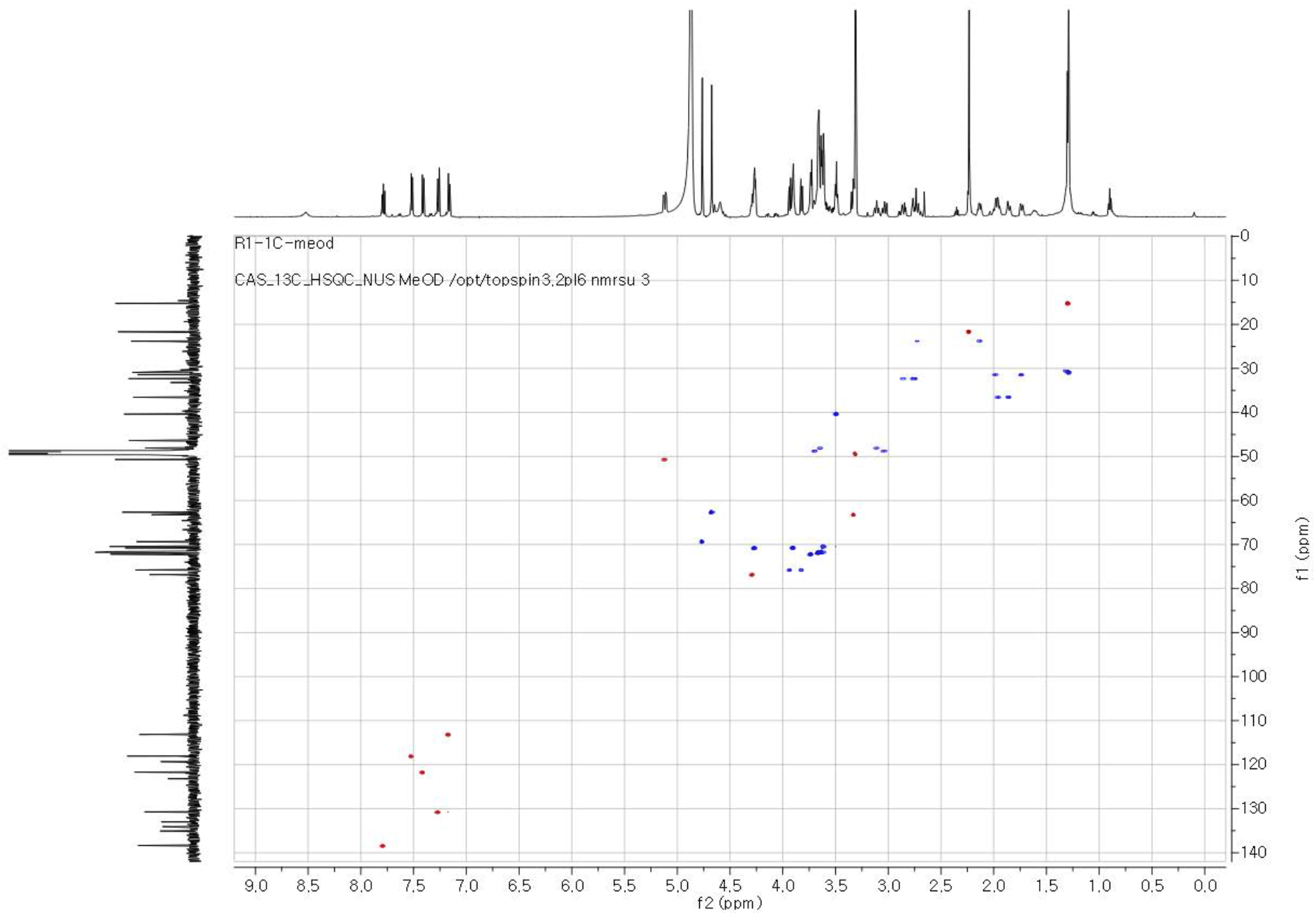
HSQC NMR spectrum of R1-1C in MeOH-*d*4.

**Figure S15.**
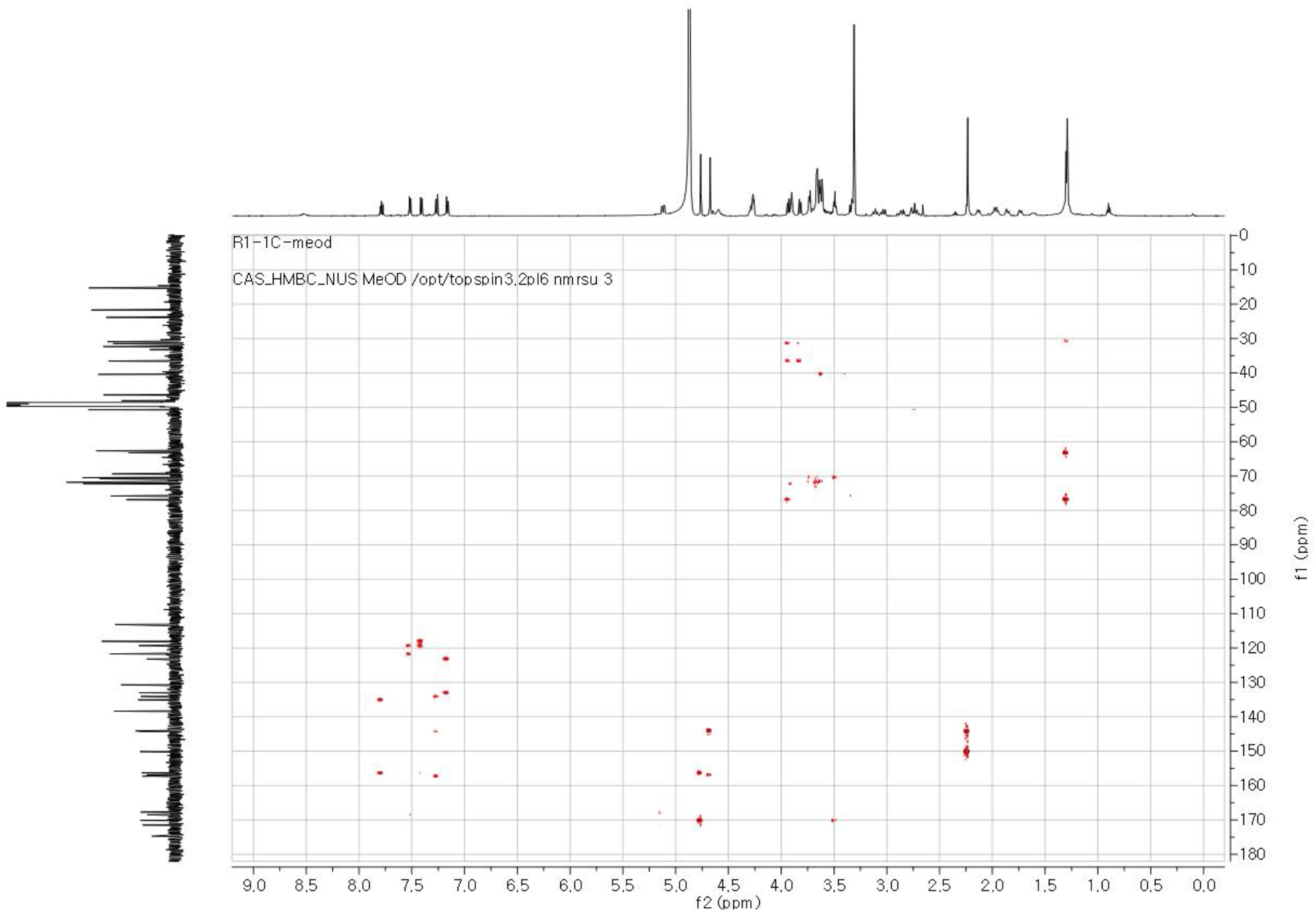
HMBC NMR spectrum of R1-1C in MeOH-*d*4.

**Table S2.** ^1^H and ^13^C NMR data of R1-1C in MeOH-*d*4.

| no. | $\delta_{\text{H}}$, mult (*J* in Hz)<sup>a</sup> | $\delta_{\text{C}}$<sup>b</sup> | |
| --- | --- | --- | --- |
| 1 |  | 174.7 | C |
| 2a | 2.86, m | 32.3 | CH<sub>2</sub> |
| 2b | 2.74, m |  |  |
| 3a | 2.72, m | 23.8 | CH<sub>2</sub> |
| 3b | 2.14, m |  |  |
| 4 | 5.11, m | 50.7 | CH |
| 5 |  | 171.5 | C |
| 6 |  | 167.7 | C |
| 7 |  | 135.1 | C |
| 8 | 7.52, d (8.0) | 118.1 | CH |
| 9 | 7.78, t (8.0) | 138.4 | CH |
| 10 | 7.41, d (8.0) | 121.7 | CH |
| 11 |  | 156.4 | C |
| 12 |  | 119.3 | C |
| 13 |  | 168.5 | C |
| 14 | 4.77, s | 69.4 | CH<sub>2</sub> |
| 15 |  | 170.1 | C |
| 16 | 3.50, m | 40.4 | CH<sub>2</sub> |
| 17-21 | 3.62-3.67 | 72.1-69.3 | CH<sub>2</sub> |
| 22 | 3.90, m | 70.7 | CH<sub>2</sub> |
| 23 | 4.27, m | 70.8 | CH<sub>2</sub> |
| 24 |  | 157.3 | C |
| 25 |  | 123.2 | C |
| 26 |  | 134.1 | C |
| 27 |  | 133.0 | C |
| 28 | 7.27, d (8.0) | 130.7 | CH |
| 29 | 7.16, d (8.0) | 113.1 | CH |
| 30 |  | 144.3 | C |
| 31 |  | 150.1 | C |
| 32 | 2.24, s | 21.7 | CH<sub>3</sub> |
| 33 |  | 156.9 | C |
| 34 |  | 144.1 | C |
| 35 | 4.68, s | 62.7 | CH<sub>2</sub> |
| 36a | 3.64, m | 48.1 | CH<sub>2</sub> |
| 36b | 3.10, m |  |  |
| 37a | 1.98, m | 30.9 | CH<sub>2</sub> |
| 37b | 1.73, m |  |  |
| 38 |  | 46.4 | C |
| 39a | 1.96, m | 36.5 | CH<sub>2</sub> |
| 39b | 1.86, m |  |  |
| 40a | 3.69, m | 48.8 | CH<sub>2</sub> |
| 40b | 3.04, m |  |  |
| 41 | 3.32, t, (5.0) | 63.2 | CH |
| 42 | 4.28, m | 76.8 | CH |
| 43 | 1.30, d (6.5) | 15.3 | CH<sub>3</sub> |
| 44a | 3.93, d (8.5) | 75.8 | CH<sub>2</sub> |
| 44b | 3.82, d (8.5) |  |  |

<sup>a</sup>. <sup>1</sup>H: 600 MHz,<sup>b</sup> <sup>13</sup>C: 150 MHz

**Figure S16.**
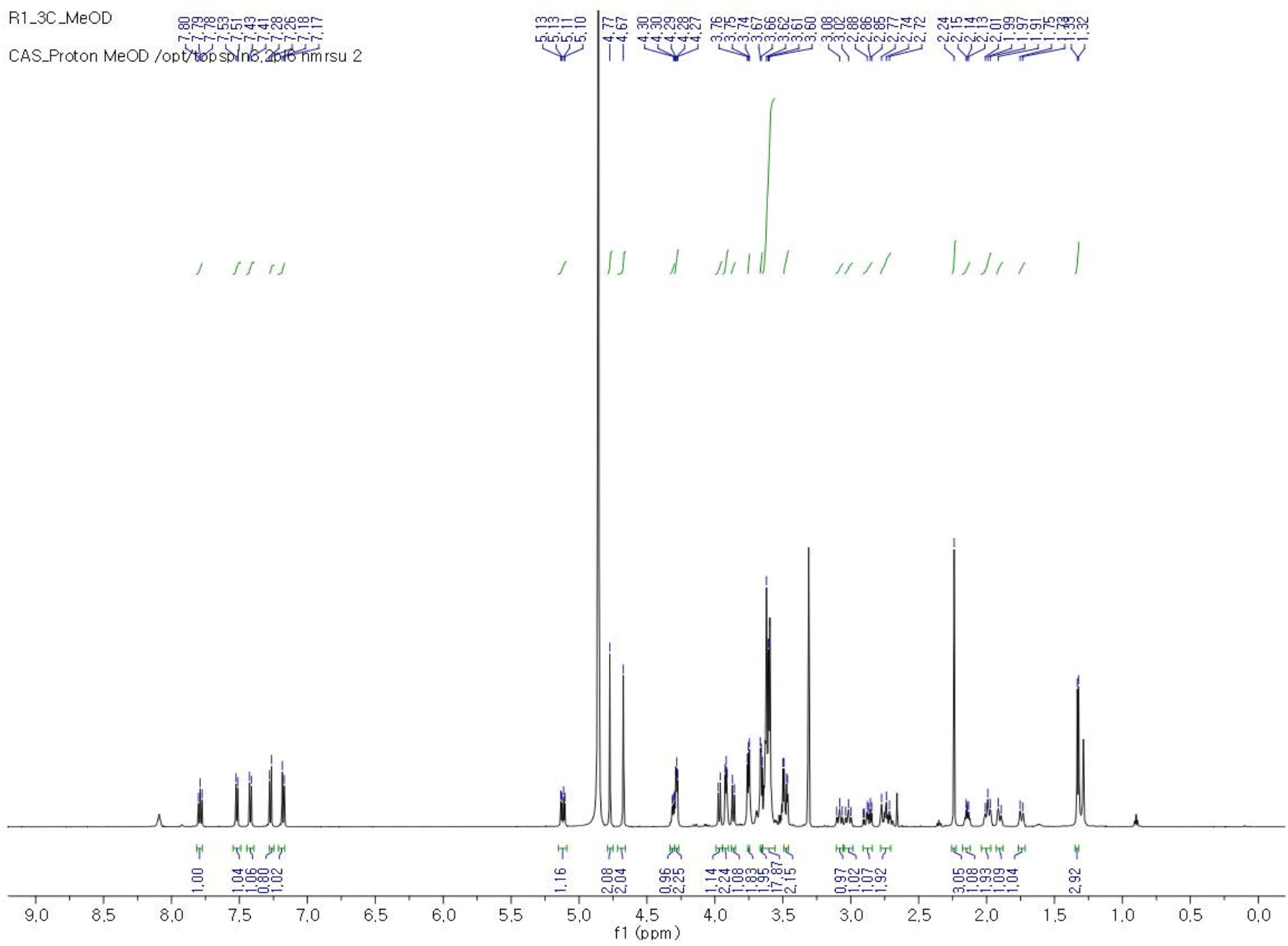
^1^H NMR spectrum of R1-3C in MeOH-*d*4.

**Figure S17.**
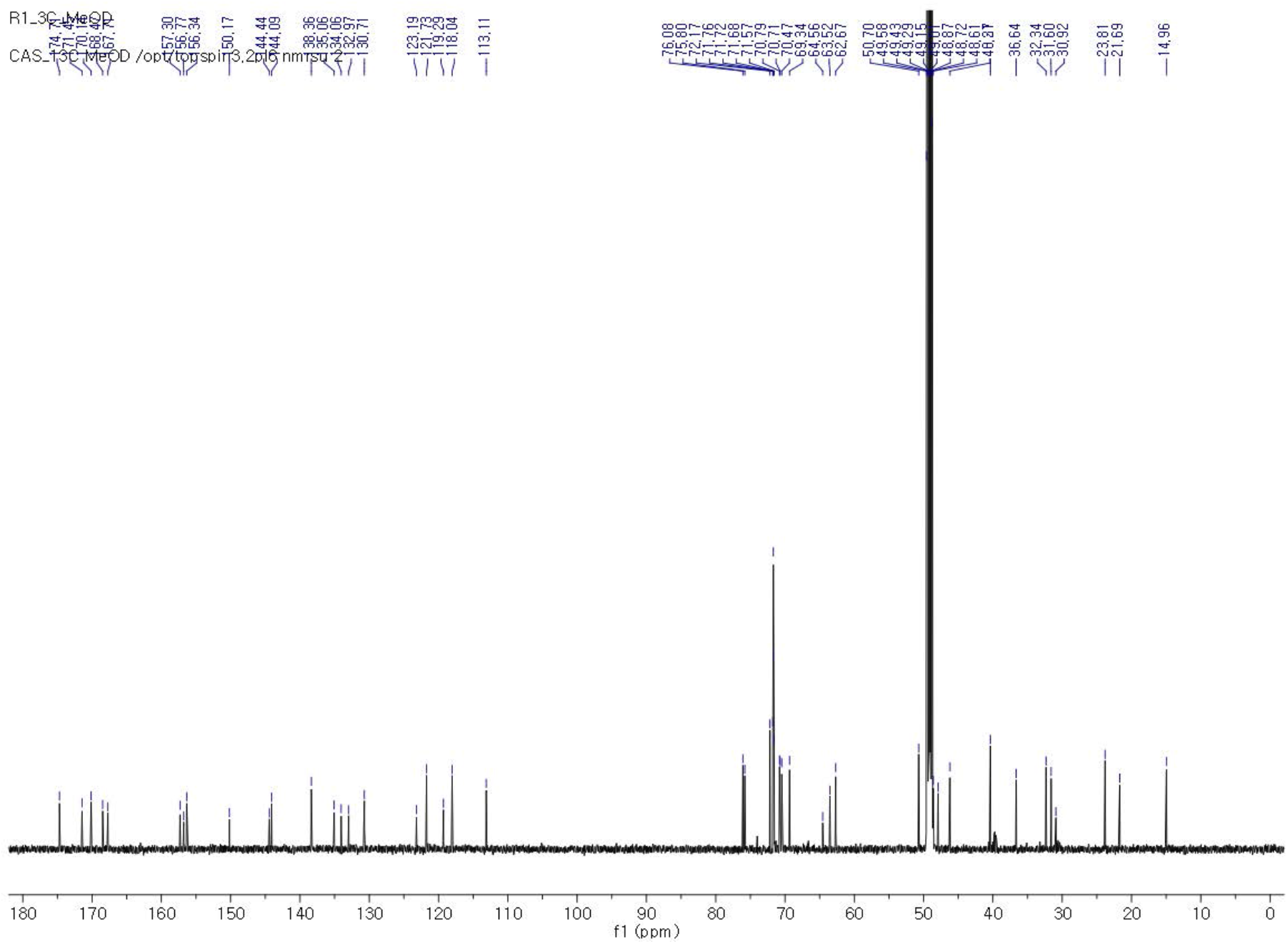
^13^C NMR spectrum of R1-3C in MeOH-*d*4.

**Figure S18.**
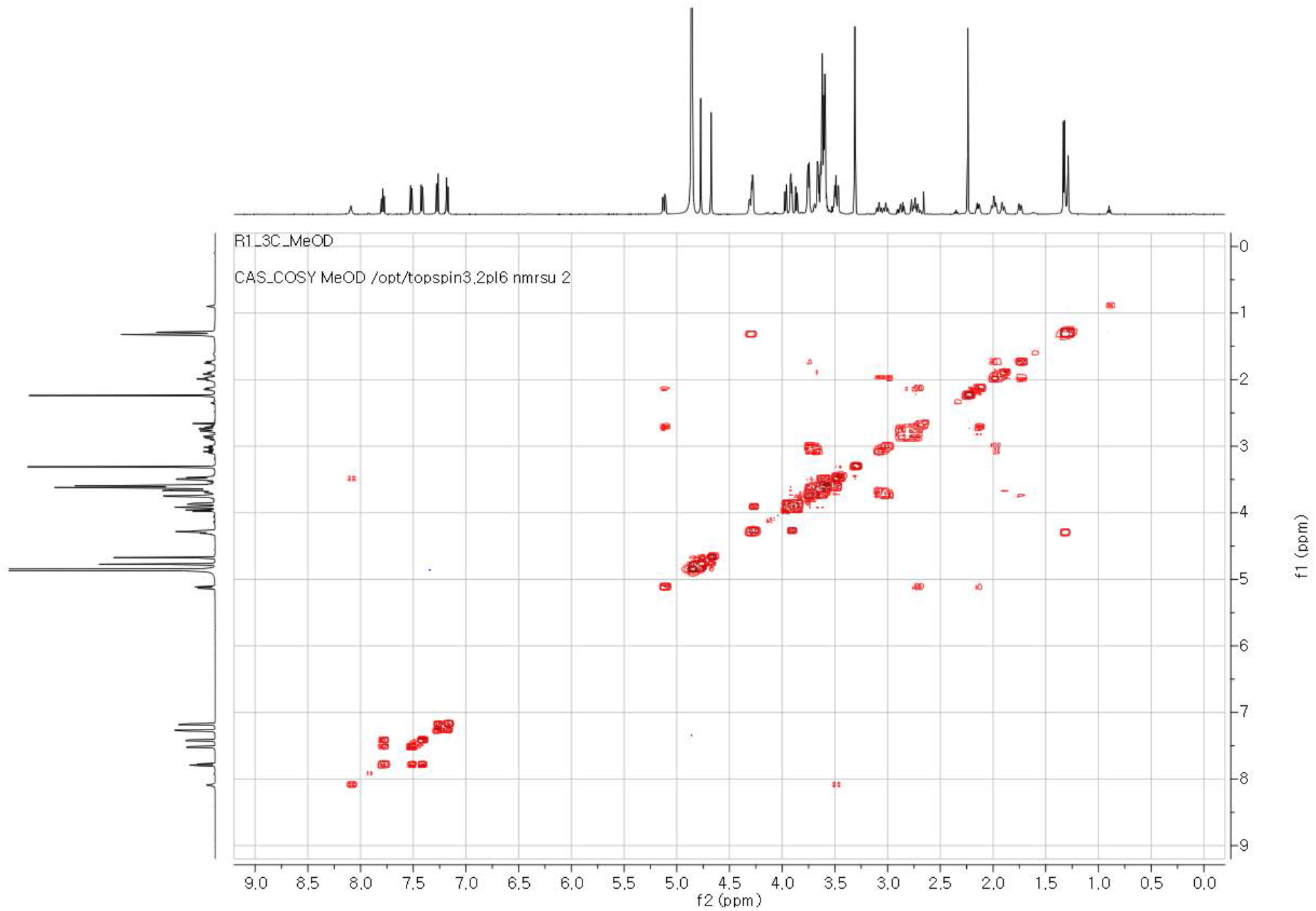
COSY NMR spectrum of R1-3C in MeOH-*d*4.

**Figure S19.**
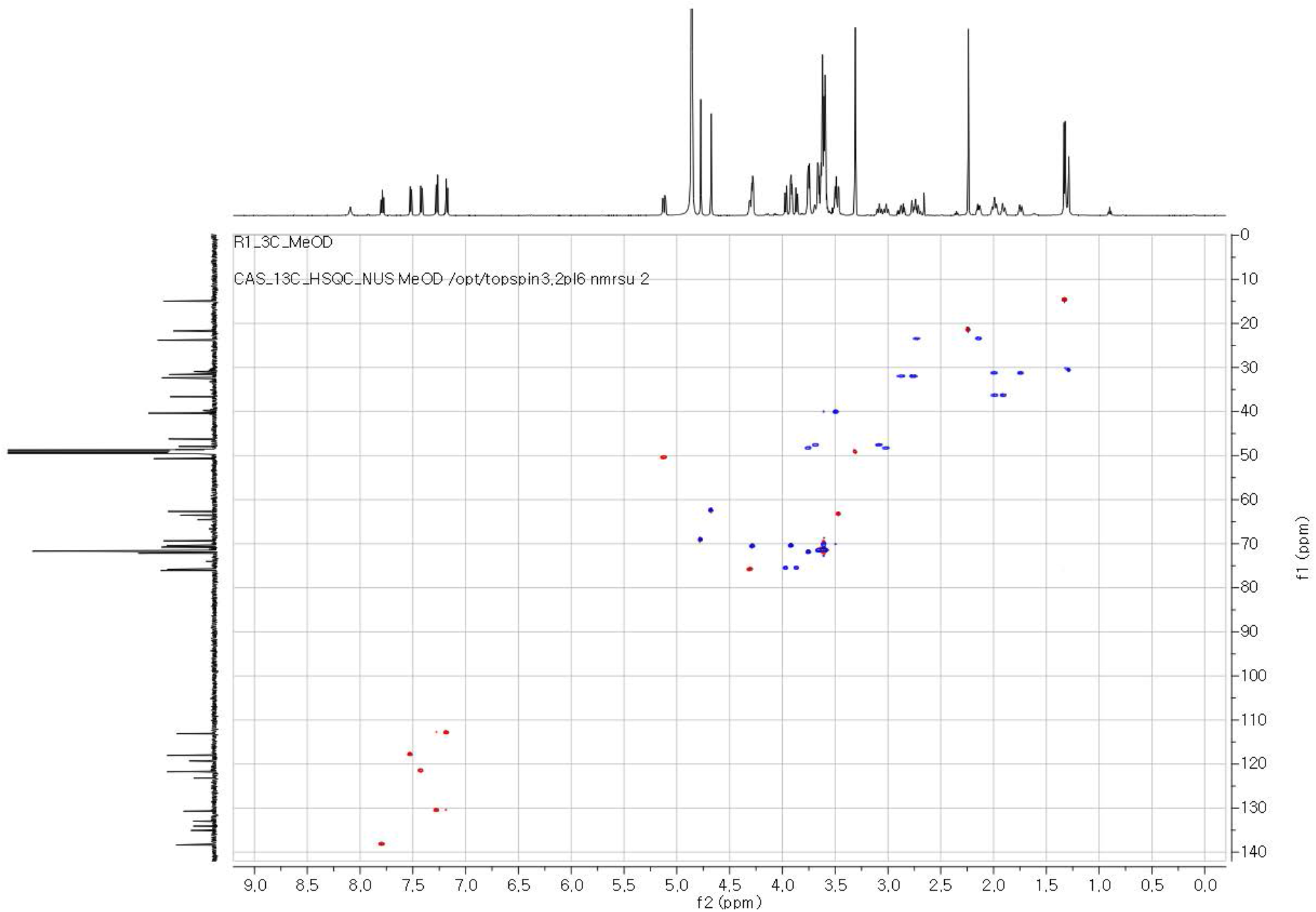
HSQC NMR spectrum of R1-3C in MeOH-*d*4.

**Figure S20.**
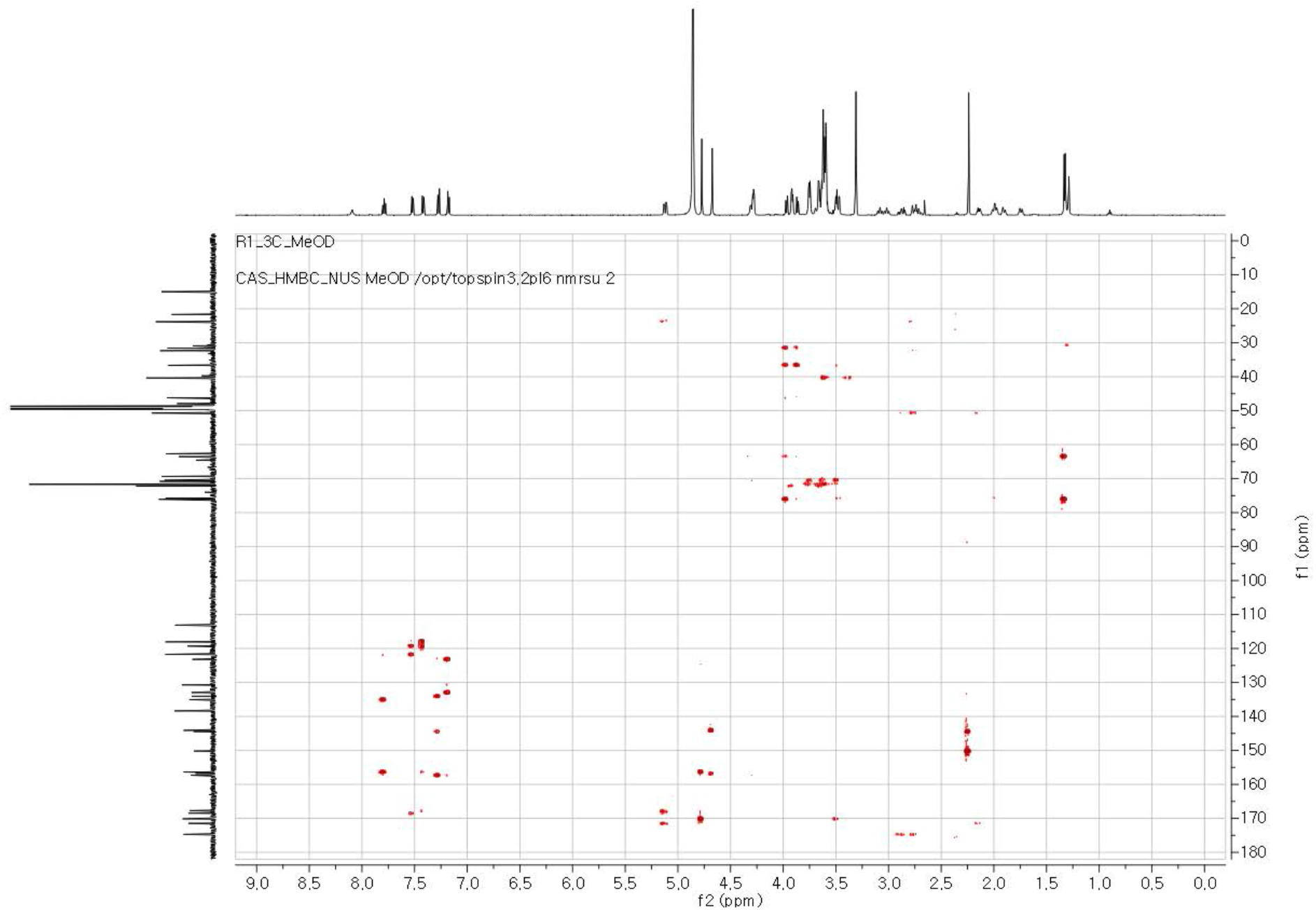
HMBC NMR spectrum of R1-3C in MeOH-*d*4.

**Table S3.** ^1^H and ^13^C NMR data of R1-3C in MeOH-*d*4.

| no. | $\delta_{\text{H}}$, mult ($J$ in Hz)<sup>a</sup> | $\delta_{\text{C}}$<sup>b</sup> | |
| --- | --- | --- | --- |
| 1 |  | 174.7 | C |
| 2a | 2.86, m | 32.3 | $\text{CH}_2$ |
| 2b | 2.74, m |  |  |
| 3a | 2.72, m | 23.8 | $\text{CH}_2$ |
| 3b | 2.14, m |  |  |
| 4 | 5.11, m | 50.7 | CH |
| 5 |  | 171.5 | C |
| 6 |  | 167.7 | C |
| 7 |  | 135.1 | C |
| 8 | 7.53, d (8.0) | 118.1 | CH |
| 9 | 7.79, t (8.0) | 138.4 | CH |
| 10 | 7.42, d (8.0) | 121.7 | CH |
| 11 |  | 156.3 | C |
| 12 |  | 119.3 | C |
| 13 |  | 168.5 | C |
| 14 | 4.77, s | 69.3 | $\text{CH}_2$ |
| 15 |  | 170.1 | C |
| 16 | 3.50, m | 40.4 | $\text{CH}_2$ |
| 17-25 | 3.59-3.62 | 72.2-69.3 | $\text{CH}_2$ |
| 26 | 3.91, m | 70.7 | $\text{CH}_2$ |
| 27 | 4.28, m | 70.8 | $\text{CH}_2$ |
| 28 |  | 157.3 | C |
| 29 |  | 123.2 | C |
| 30 |  | 134.1 | C |
| 31 |  | 133.0 | C |
| 32 | 7.27, d (8.0) | 130.7 | CH |
| 33 | 7.18, d (8.0) | 113.1 | CH |
| 34 |  | 144.4 | C |
| 35 |  | 150.2 | C |
| 36 | 2.24, s | 21.7 | $\text{CH}_3$ |
| 37 |  | 156.8 | C |
| 38 |  | 144.1 | C |
| 39 | 4.67, s | 62.7 | $\text{CH}_2$ |
| 40a | 3.66, m | 47.9 | $\text{CH}_2$ |
| 40b | 3.08, m |  |  |
| 41a | 1.99, m | 30.9 | $\text{CH}_2$ |
| 41b | 1.74, m |  |  |
| 42 |  | 46.2 | C |
| 43a | 1.98, m | 36.5 | $\text{CH}_2$ |
| 43b | 1.90, m |  |  |
| 44a | 3.75, m | 48.6 | $\text{CH}_2$ |
| 44b | 3.02, m |  |  |
| 45 | 3.32, t, (5.0) | 63.5 | CH |
| 46 | 4.28, m | 76.1 | CH |
| 47 | 1.30, d (6.5) | 15.0 | $\text{CH}_3$ |
| 48a | 3.97, d (8.5) | 75.8 | $\text{CH}_2$ |
| 48b | 3.87, d (8.5) |  |  |

<sup>a</sup>. $^1\text{H}$ : 600 MHz,<sup>b</sup> $^{13}\text{C}$ : 150 MHz

**Figure S21.**
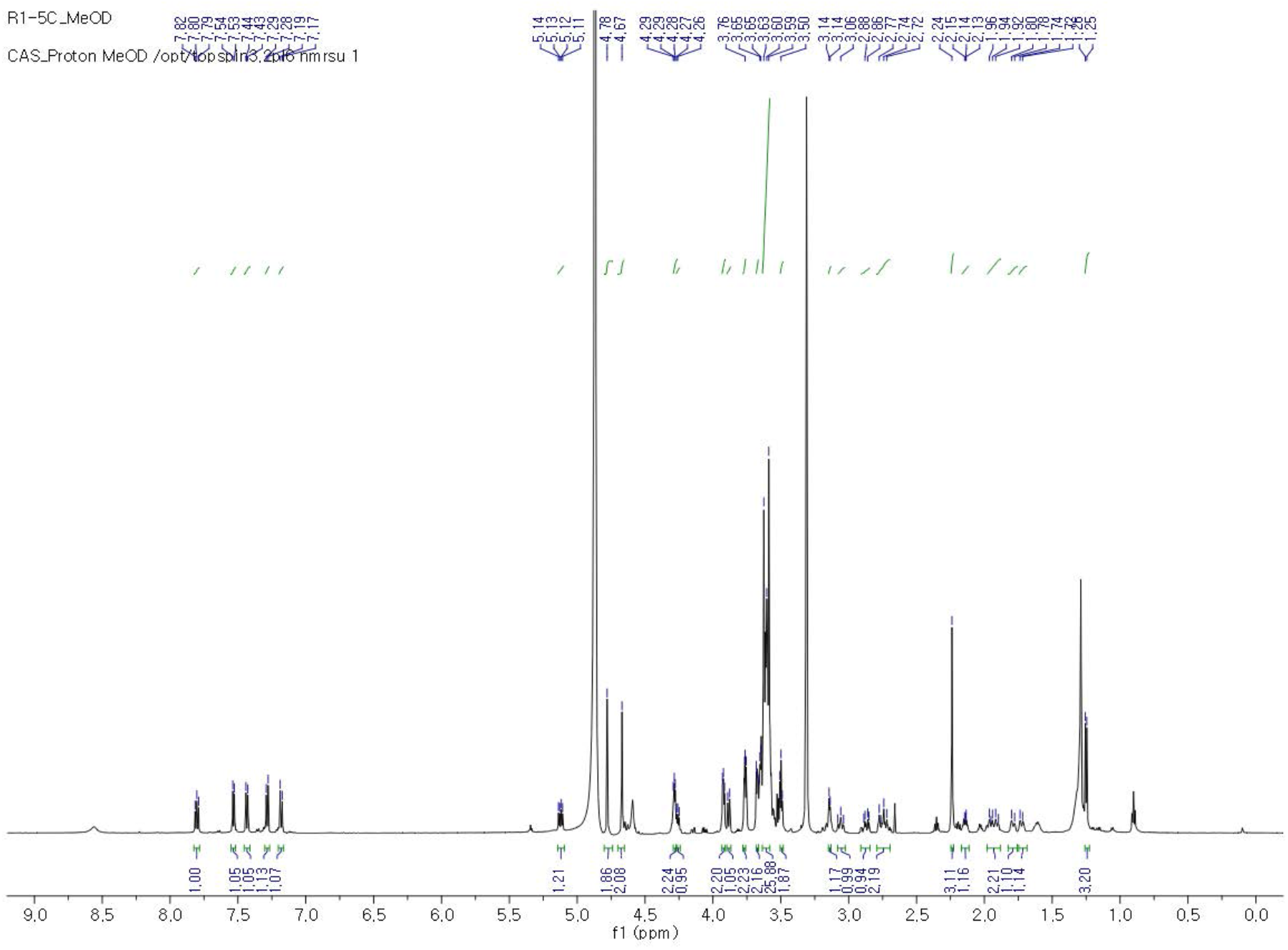
^1^H NMR spectrum of R1-5C in MeOH-*d*4.

**Figure S22.**
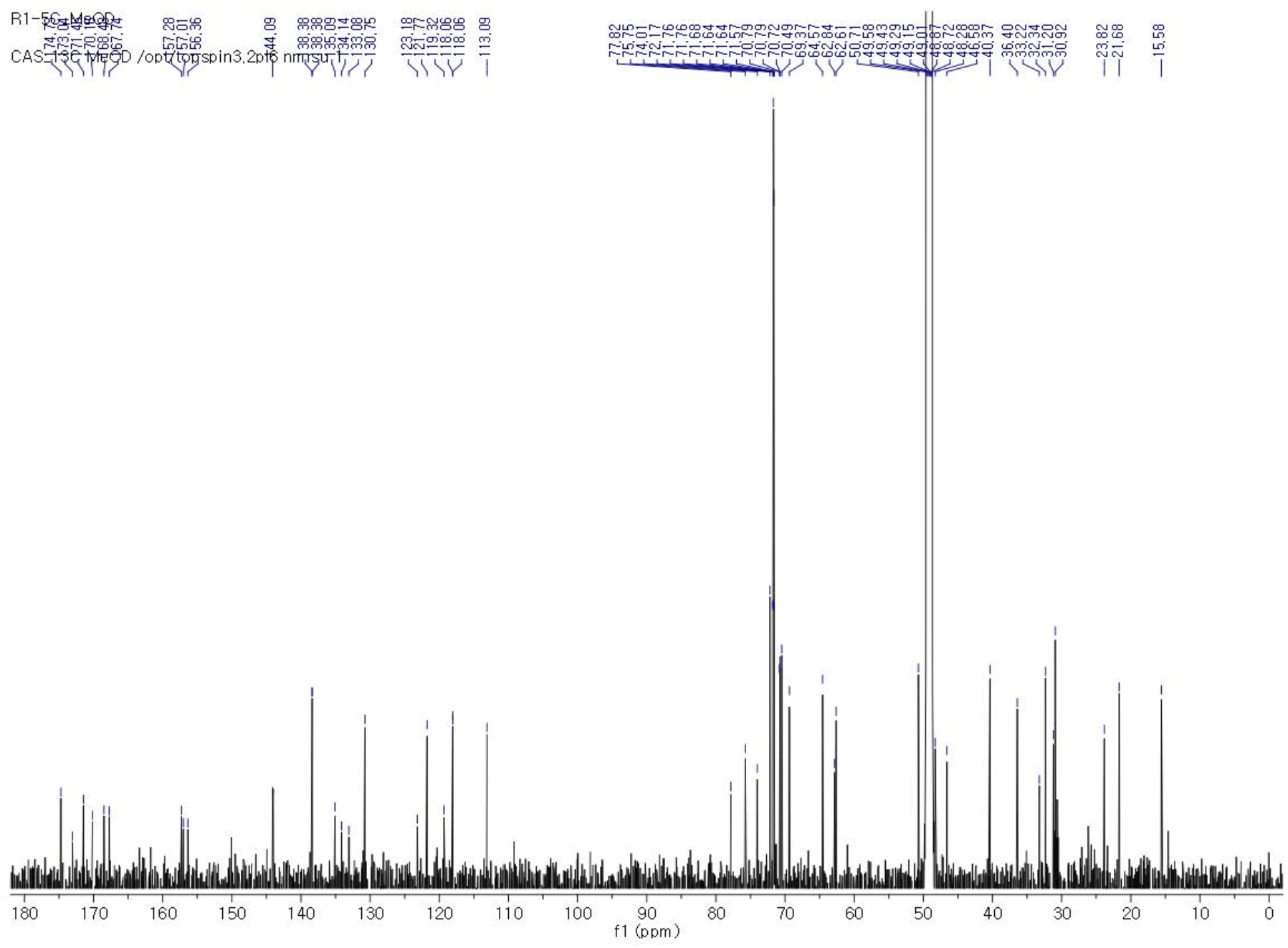
^13^C NMR spectrum of R1-5C in MeOH-*d*4.

**Figure S23.**
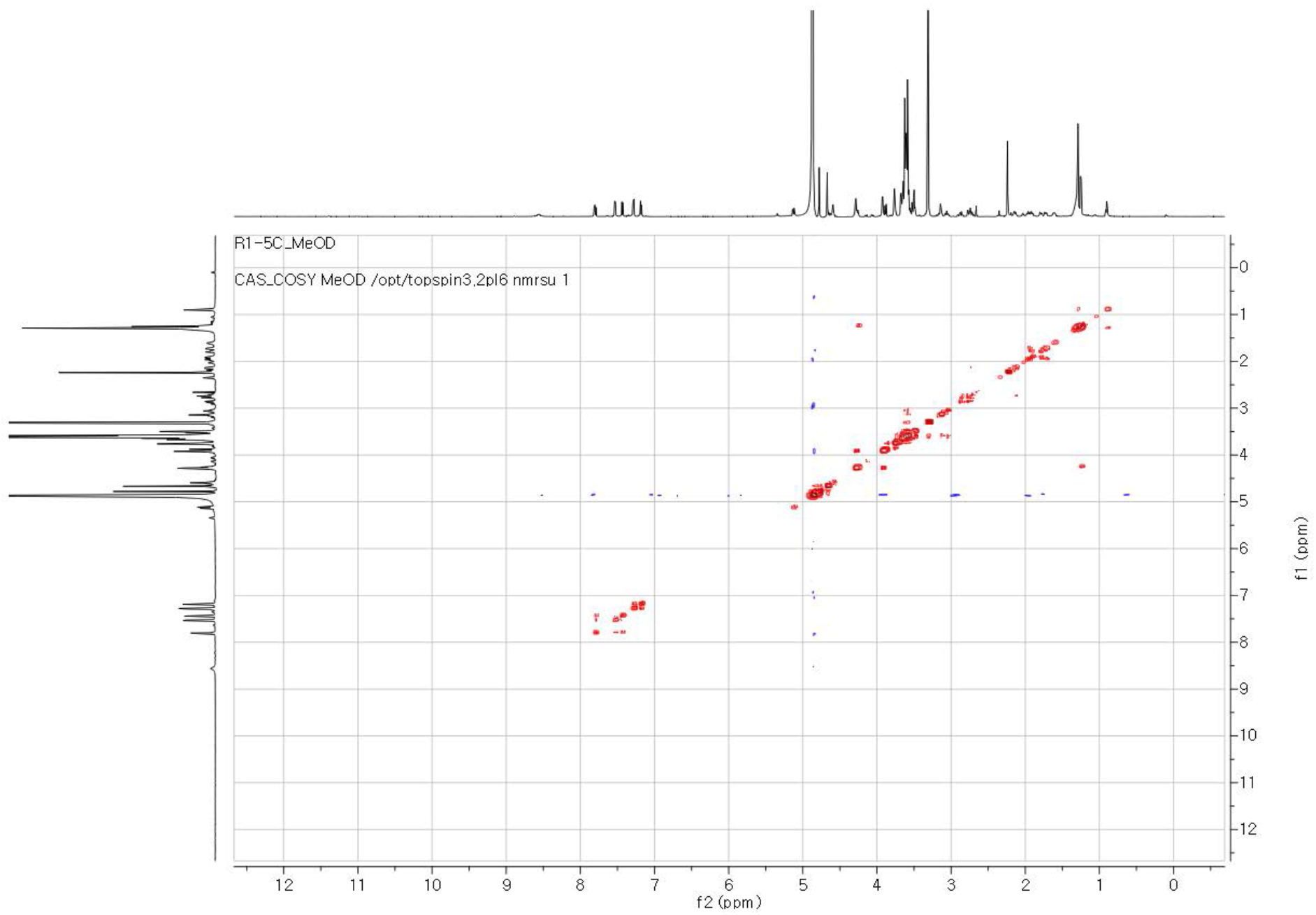
COSY NMR spectrum of R1-5C in MeOH-*d*4.

**Figure S24.**
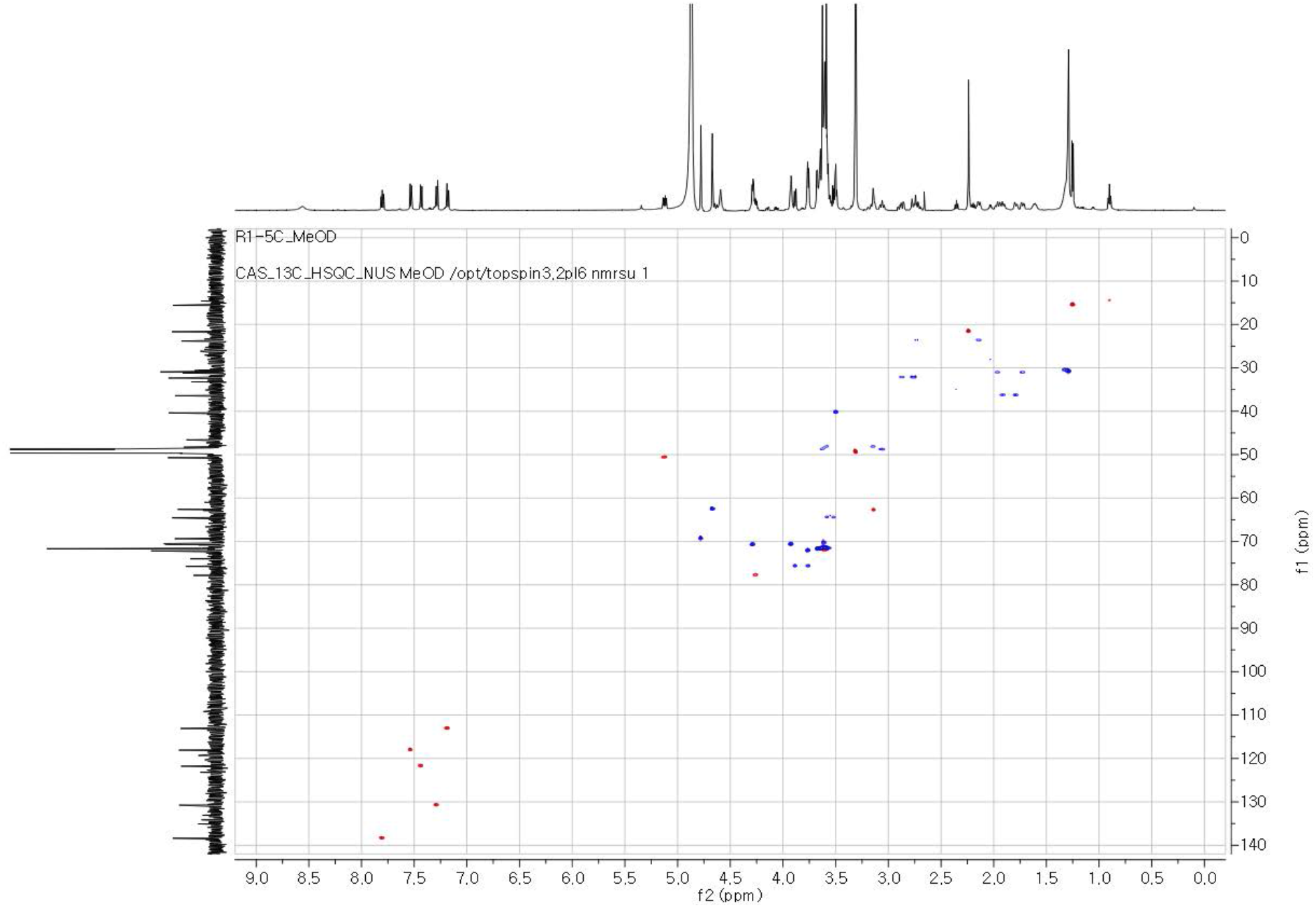
HSQC NMR spectrum of R1-5C in MeOH-*d*4.

**Figure S25.**
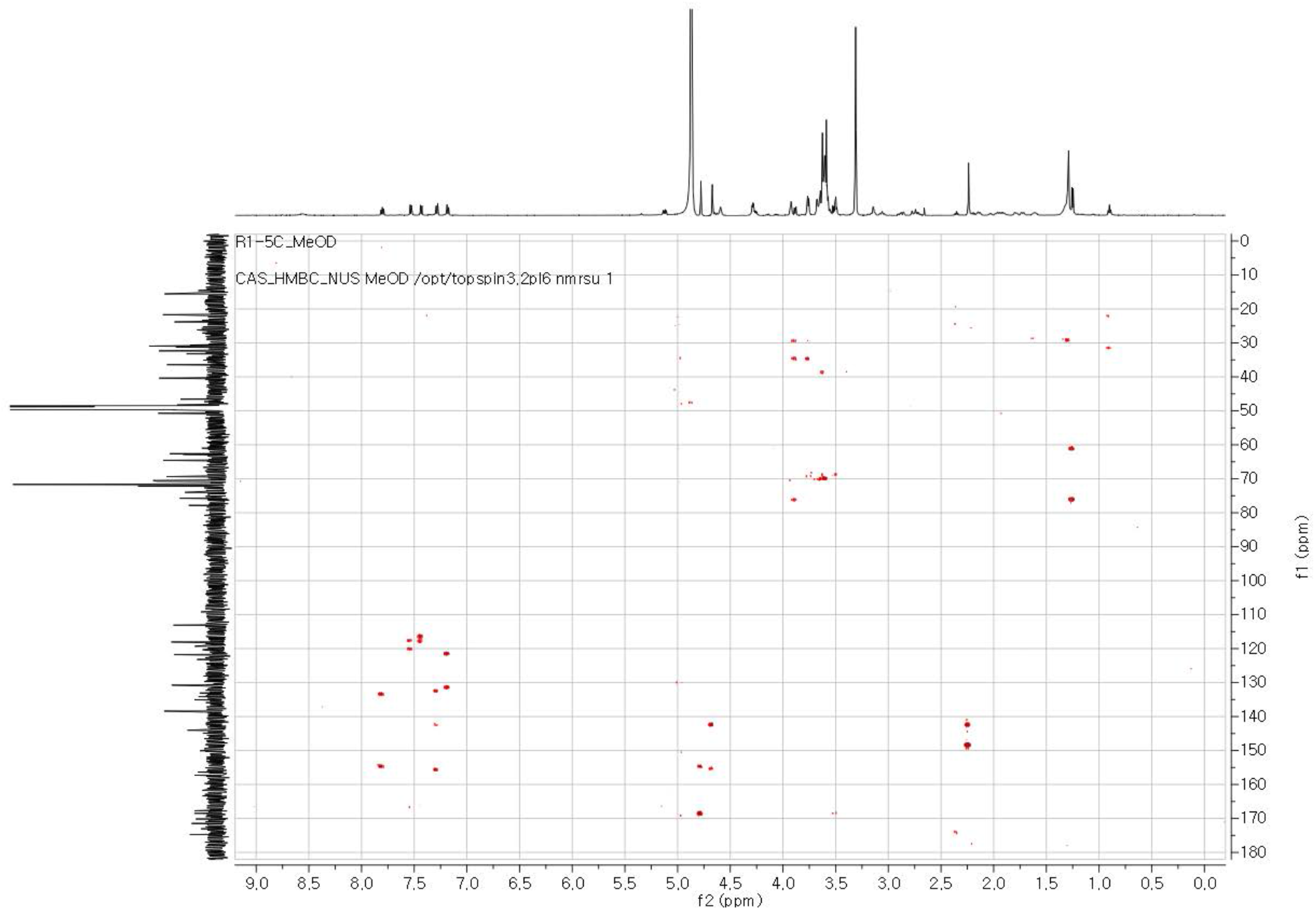
HMBC NMR spectrum of R1-5C in MeOH-*d*4.

**Table S4.** ^1^H and ^13^C NMR data of R1-5C in MeOH-*d*4.

| no. | $\delta_{\text{H}}$, mult ($J$ in Hz)<sup>a</sup> | $\delta_{\text{C}}$<sup>b</sup> | |
| --- | --- | --- | --- |
| 1 |  | 174.7 | C |
| 2a | 2.86, m | 32.3 | CH<sub>2</sub> |
| 2b | 2.77, m |  |  |
| 3a | 2.72, m | 23.8 | CH<sub>2</sub> |
| 3b | 2.14, m |  |  |
| 4 | 5.12, m | 50.7 | CH |
| 5 |  | 171.5 | C |
| 6 |  | 167.7 | C |
| 7 |  | 135.1 | C |
| 8 | 7.53, d (8.0) | 118.1 | CH |
| 9 | 7.80, t (8.0) | 138.4 | CH |
| 10 | 7.43, d (8.0) | 121.8 | CH |
| 11 |  | 156.4 | C |
| 12 |  | 119.3 | C |
| 13 |  | 168.5 | C |
| 14 | 4.78, s | 69.3 | CH<sub>2</sub> |
| 15 |  | 170.2 | C |
| 16 | 3.50, m | 40.4 | CH<sub>2</sub> |
| 17-29 | 3.59-3.62 | 72.2-69.4 | CH<sub>2</sub> |
| 30 | 3.92, m | 70.7 | CH<sub>2</sub> |
| 31 | 4.28, m | 70.8 | CH<sub>2</sub> |
| 32 |  | 157.3 | C |
| 33 |  | 123.2 | C |
| 34 |  | 134.1 | C |
| 35 |  | 133.1 | C |
| 36 | 7.27, d (8.0) | 130.8 | CH |
| 37 | 7.18, d (8.0) | 113.1 | CH |
| 38 |  | 144.4 | C |
| 39 |  | 150.2 | C |
| 40 | 2.24, s | 21.6 | CH<sub>3</sub> |
| 41 |  | 144.1 | C |
| 42 |  | 157.0 | C |
| 43 | 4.67, s | 62.6 | CH<sub>2</sub> |
| 44a | 3.67, m | 48.3 | CH<sub>2</sub> |
| 44b | 3.15, m |  |  |
| 45a | 1.96, m | 30.9 | CH<sub>2</sub> |
| 45b | 1.73, m |  |  |
| 46 |  | 46.6 | C |
| 47a | 1.92, m | 36.4 | CH<sub>2</sub> |
| 47b | 1.79, m |  |  |
| 48a | 3.76, m | 48.9 | CH<sub>2</sub> |
| 48b | 3.05, m |  |  |
| 49 | 3.14, m | 62.8 | CH |
| 50 | 4.28, m | 77.8 | CH |
| 51 | 1.30, d (6.5) | 15.6 | CH<sub>3</sub> |
| 52a | 3.92, d (8.5) | 75.8 | CH<sub>2</sub> |
| 52b | 3.89, d (8.5) |  |  |

<sup>a</sup>. $^1\text{H}$ : 600 MHz,<sup>b</sup> $^{13}\text{C}$ : 150 MHz
R1-5C

**Figure S26.**
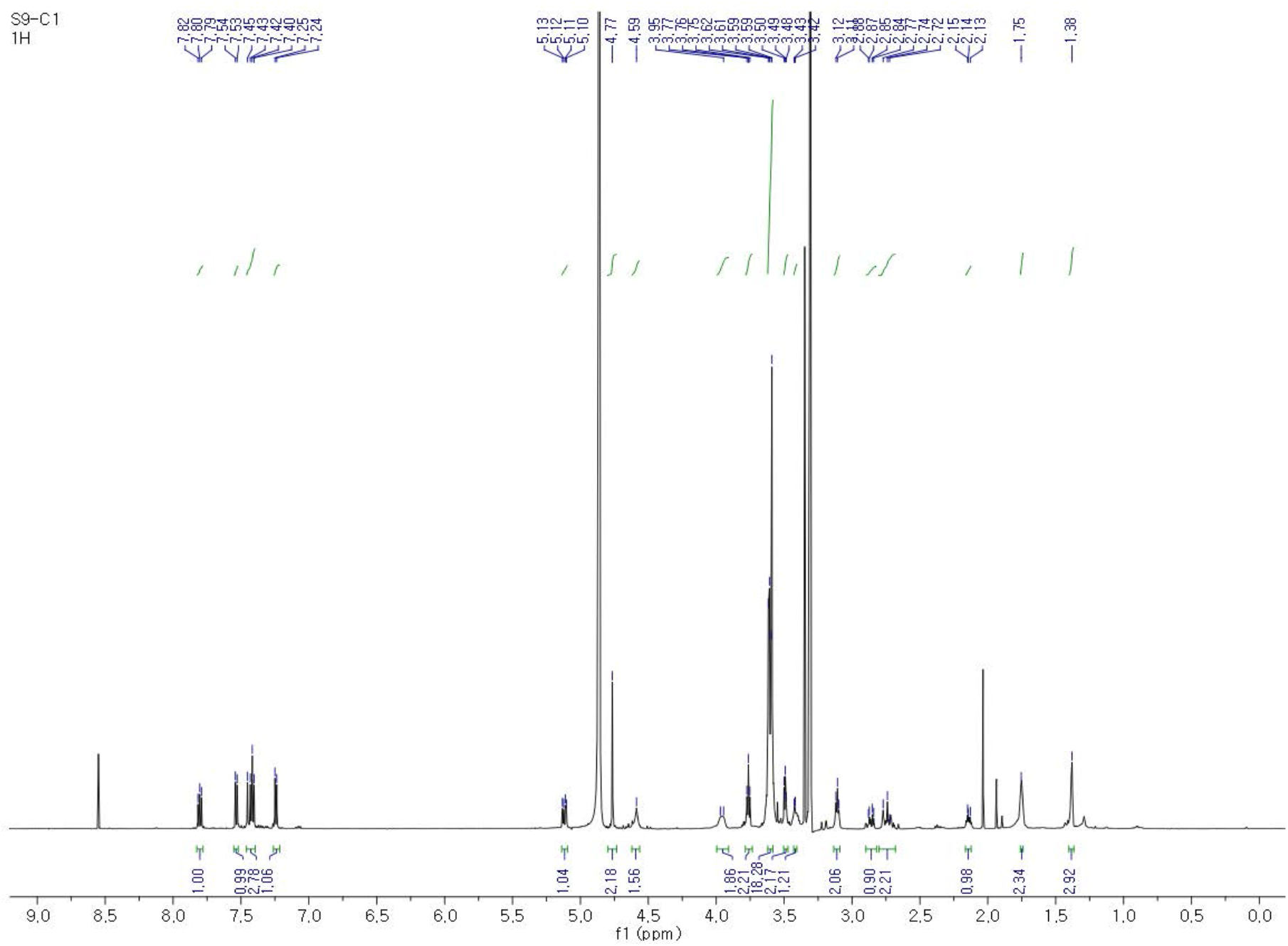
^1^H NMR spectrum of S9-1C in MeOH-*d*4.

**Figure S27.**
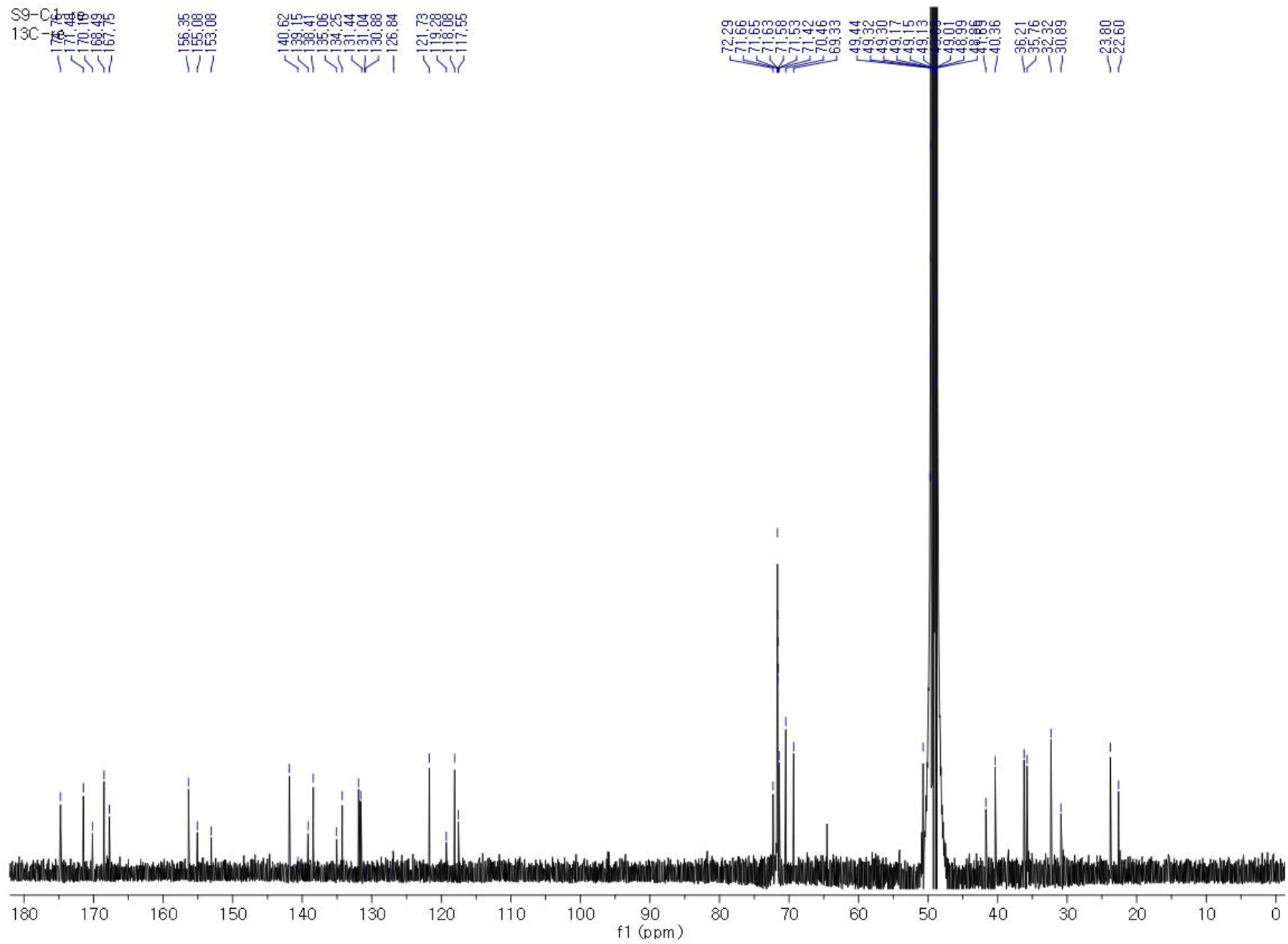
^13^C NMR spectrum of S9-1C in MeOH-*d*4.

**Figure S28.**
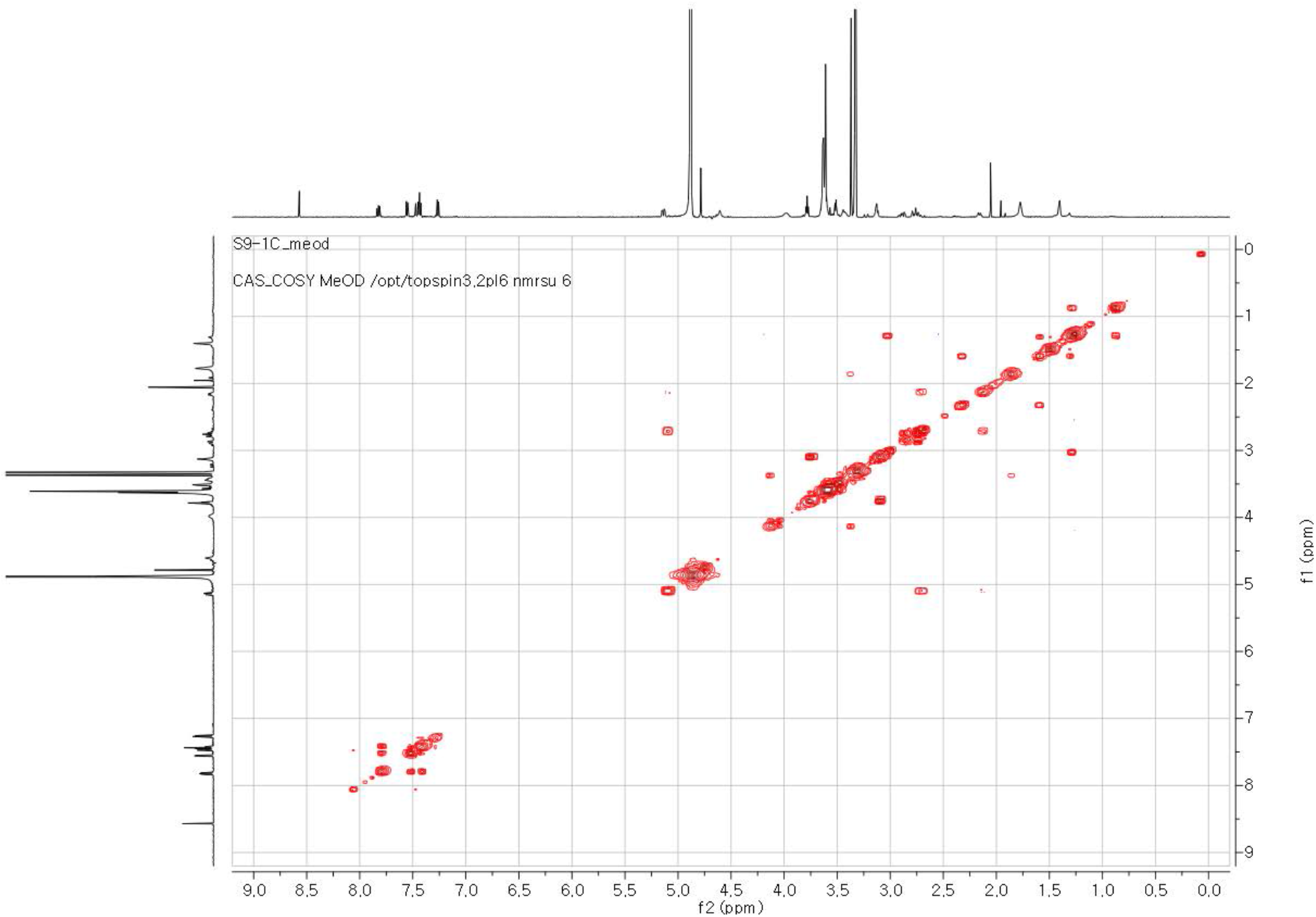
COSY NMR spectrum of S9-1C in MeOH-*d*4.

**Figure S29.**
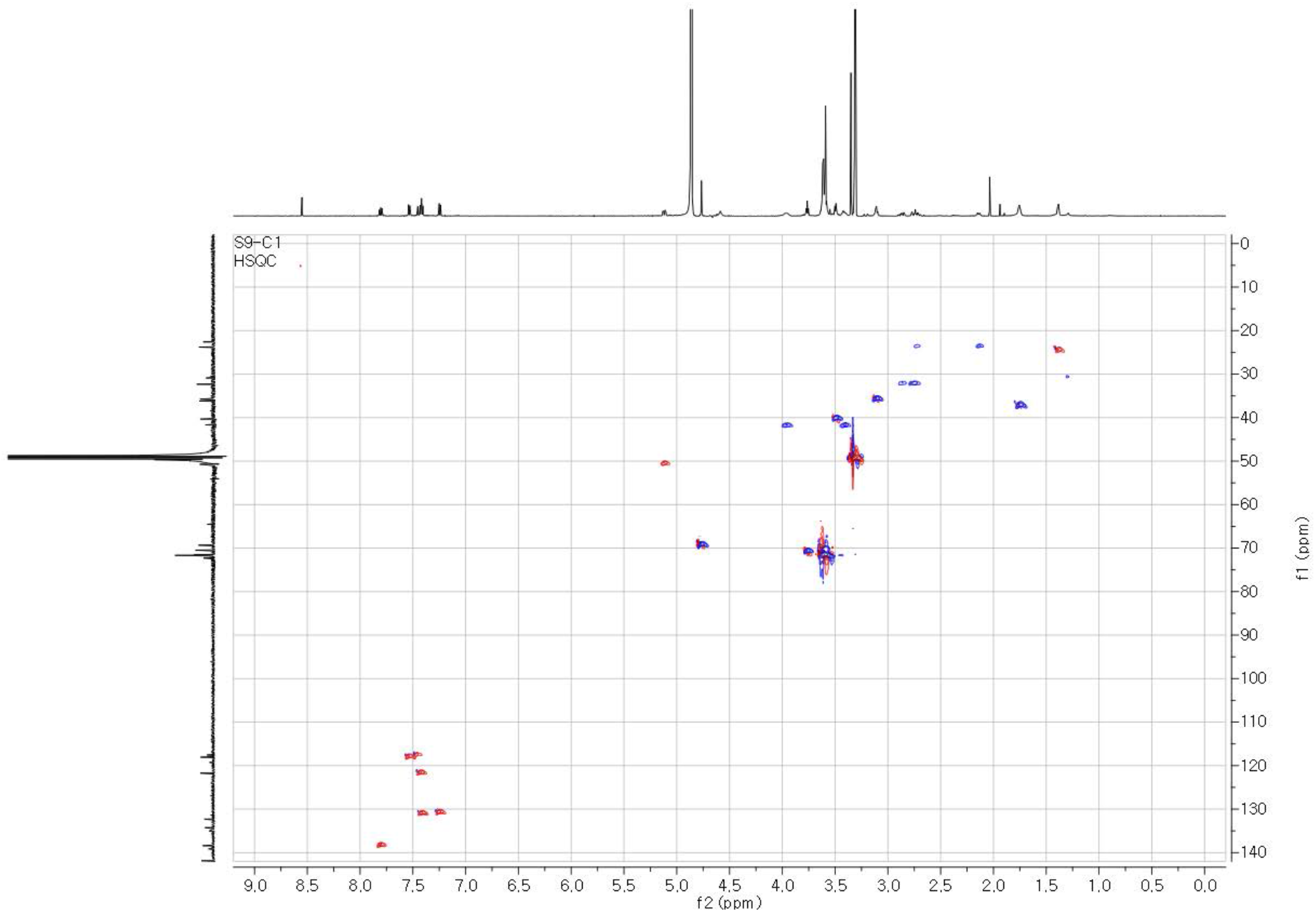
HSQC NMR spectrum of S9-1C in MeOH-*d*4.

**Figure S30.**
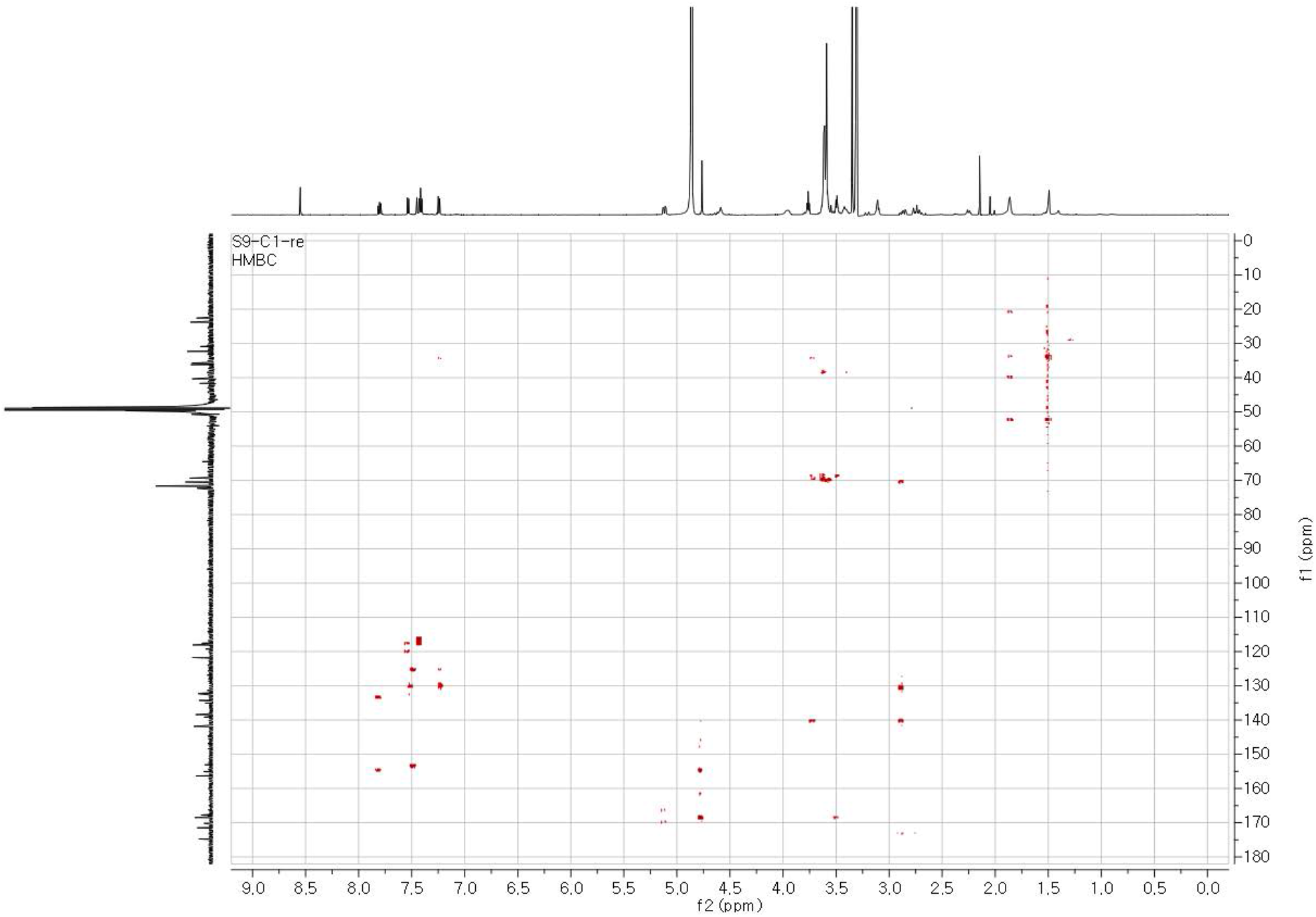
HMBC NMR spectrum of S9-1C in MeOH-*d*4.

**Table S5.**
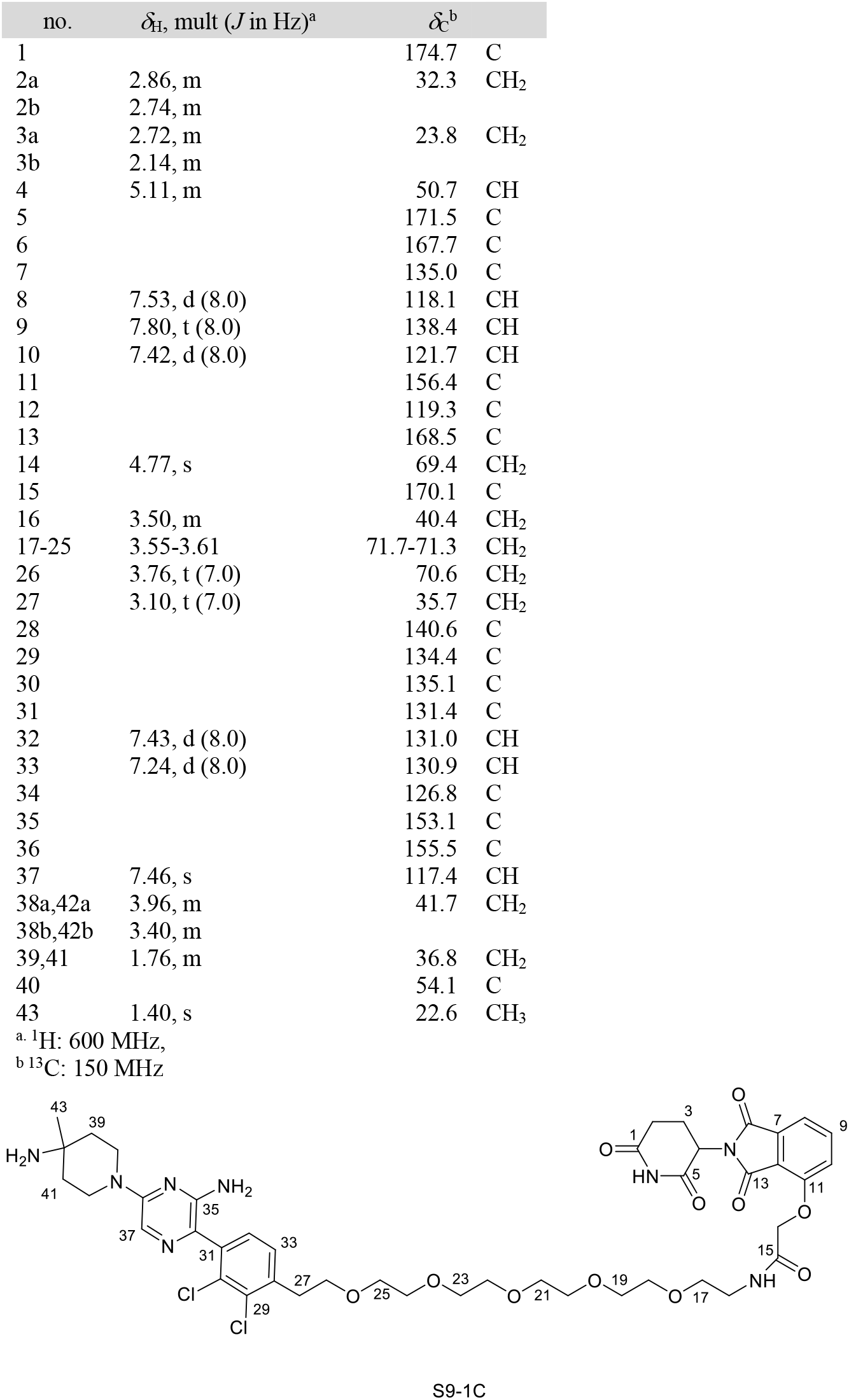
^1^H and ^13^C NMR data of S9-1C in MeOH-*d*4.

**Figure S31.**
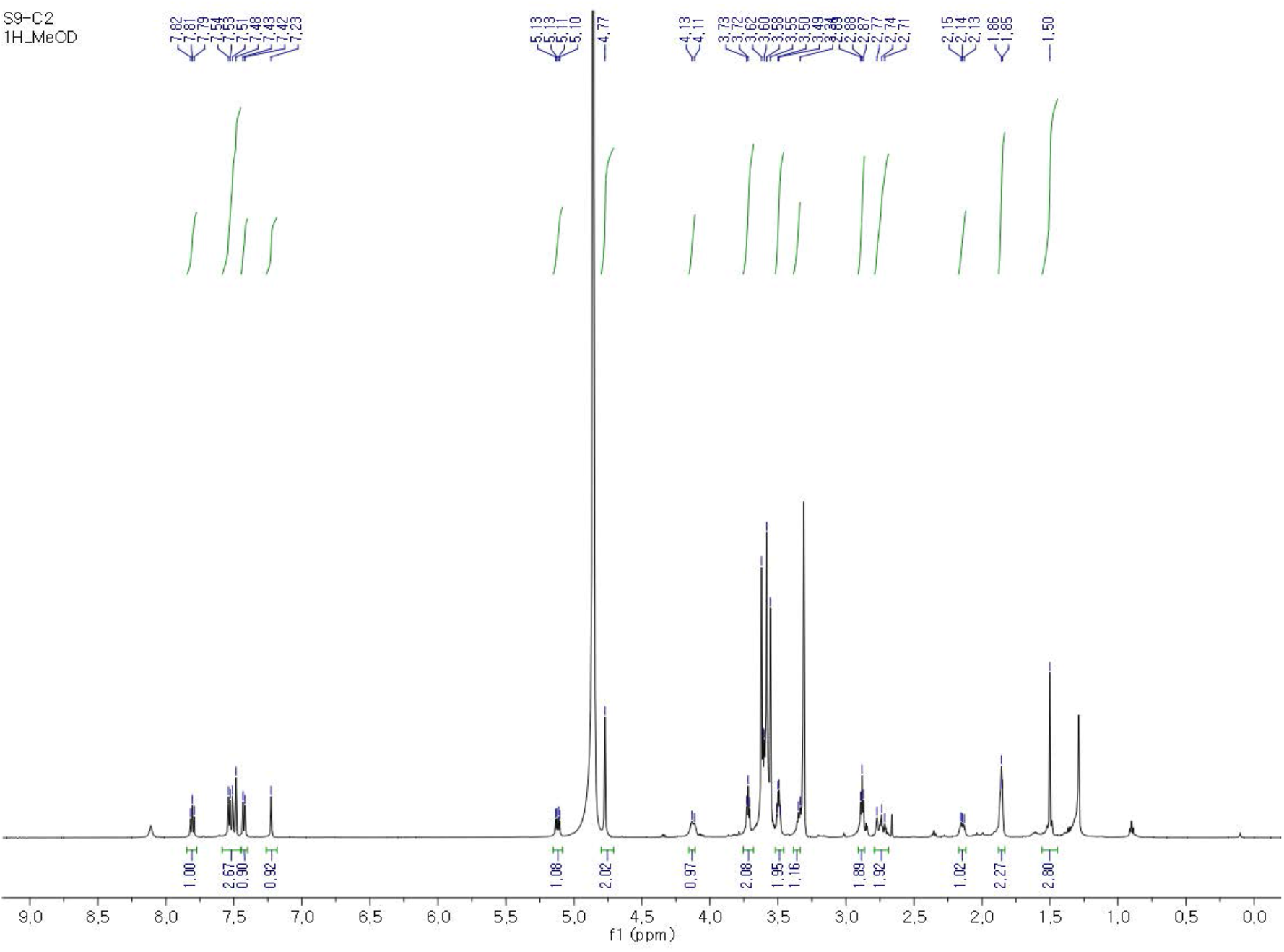
^1^H NMR spectrum of S9-2C in MeOH-*d*4.

**Figure S32.**
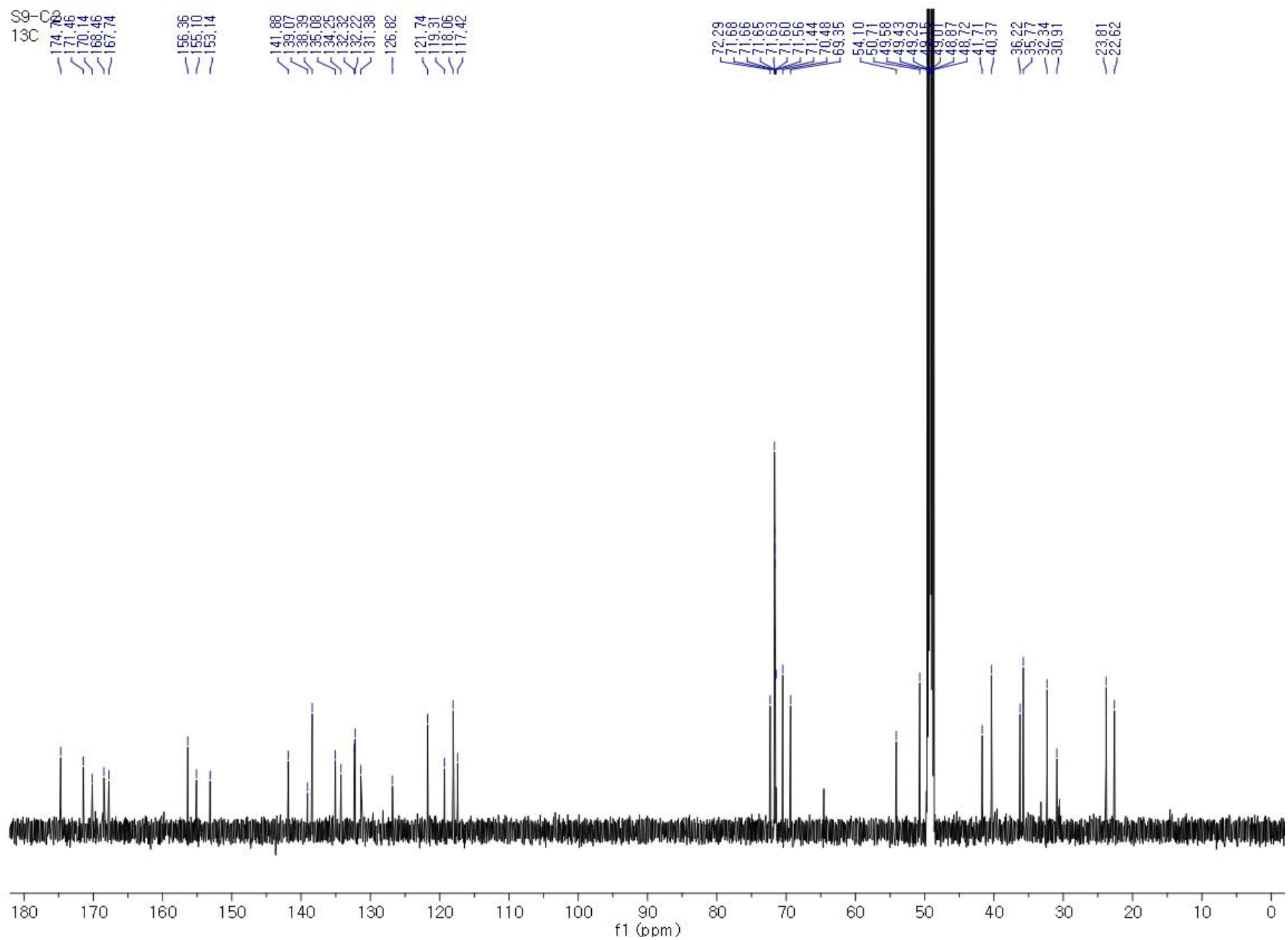
^13^C NMR spectrum of S9-2C in MeOH-*d*4.

**Figure S33.**
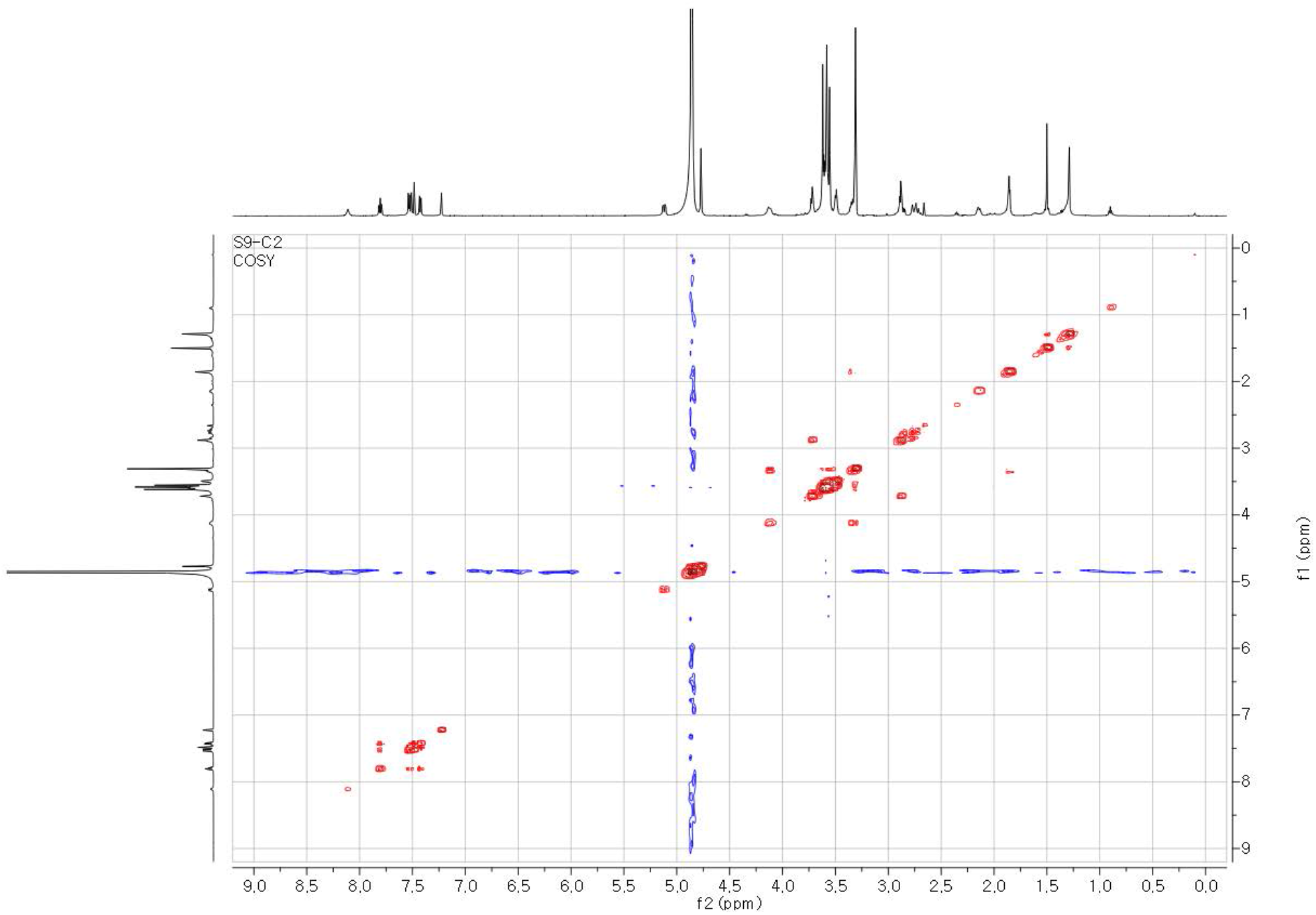
COSY NMR spectrum of S9-2C in MeOH-*d*4.

**Figure S34.**
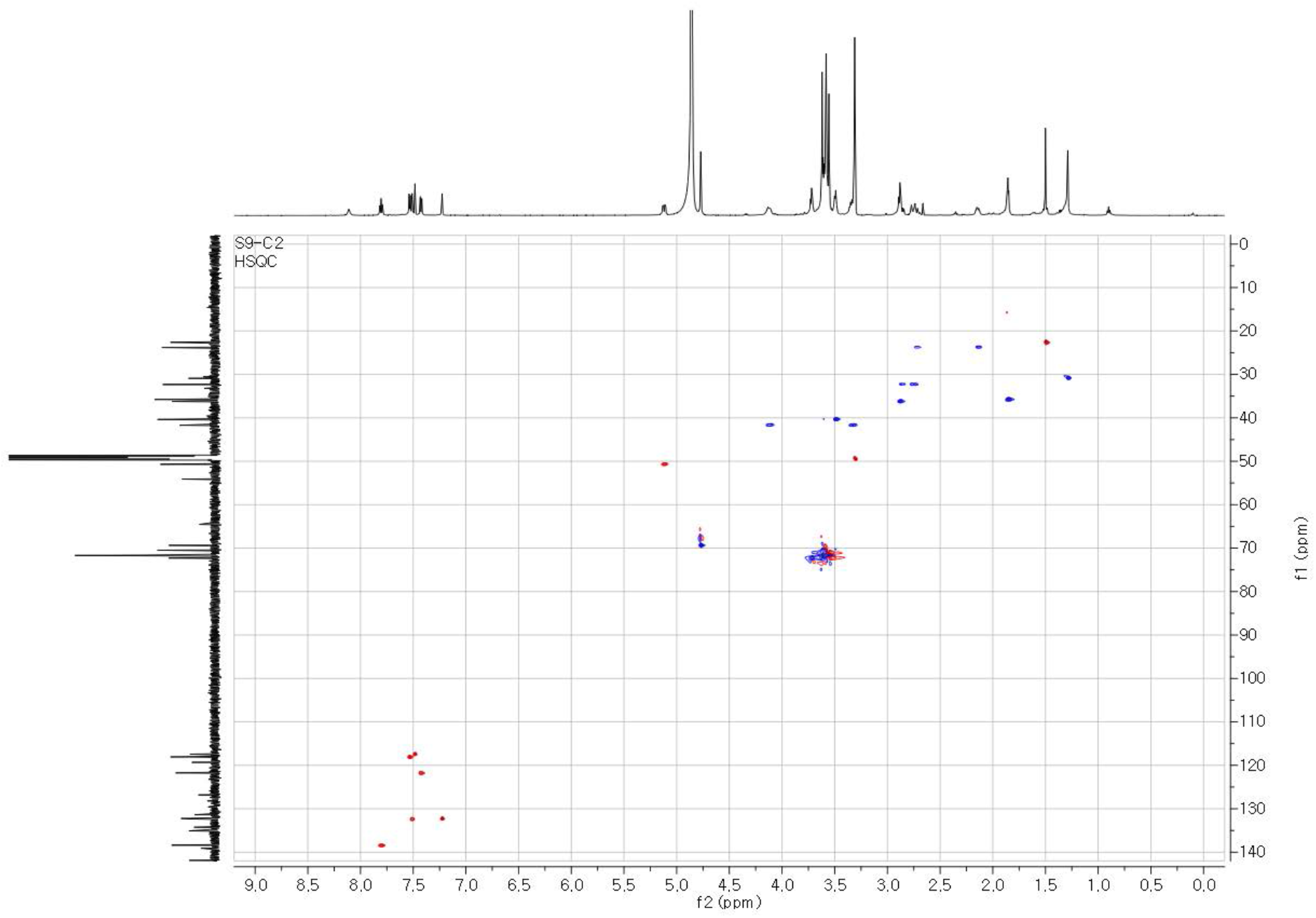
HSQC NMR spectrum of S9-2C in MeOH-*d*4.

**Figure S35.**
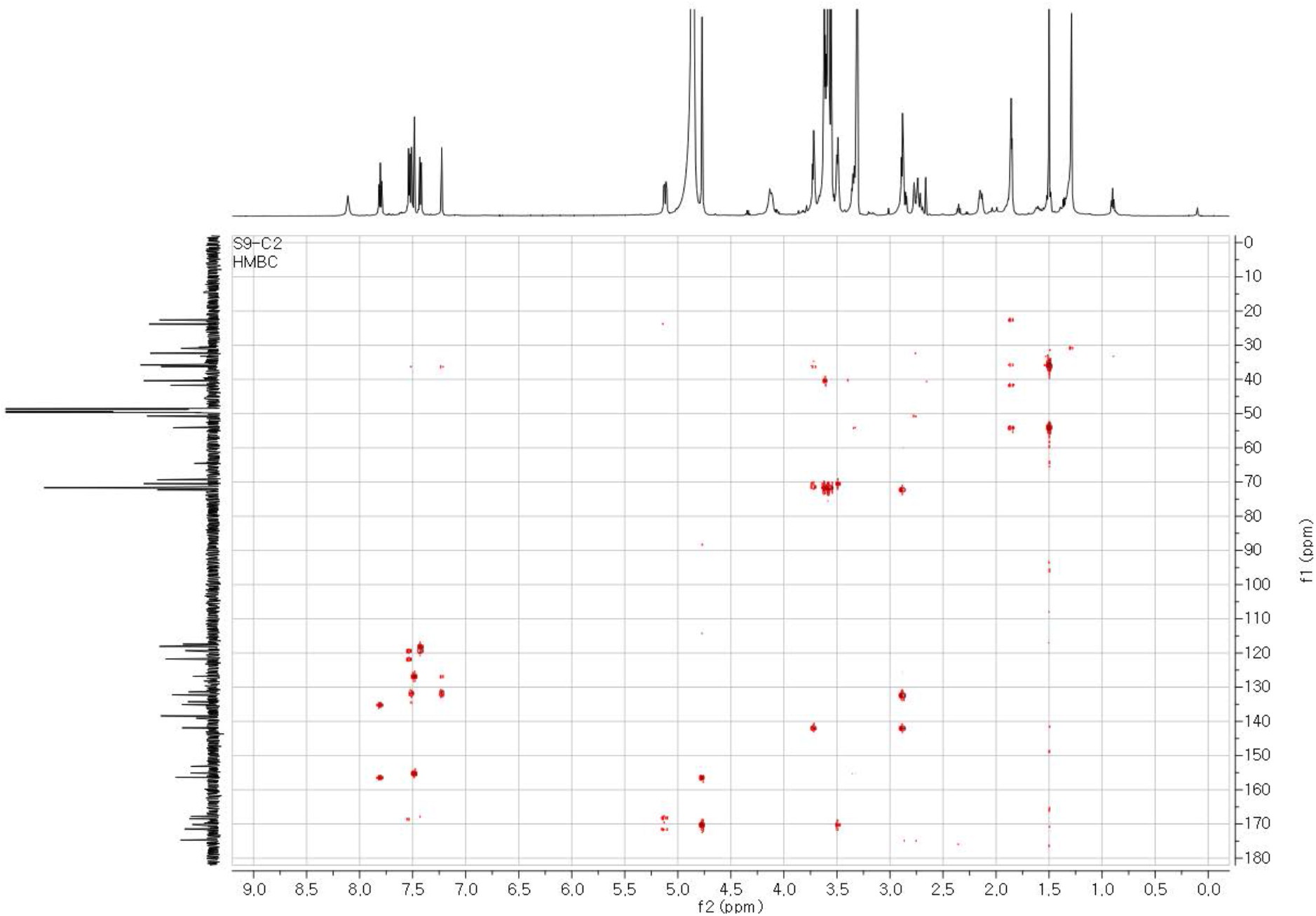
HMBC NMR spectrum of S9-2C in MeOH-*d*4.

**Table S6.**
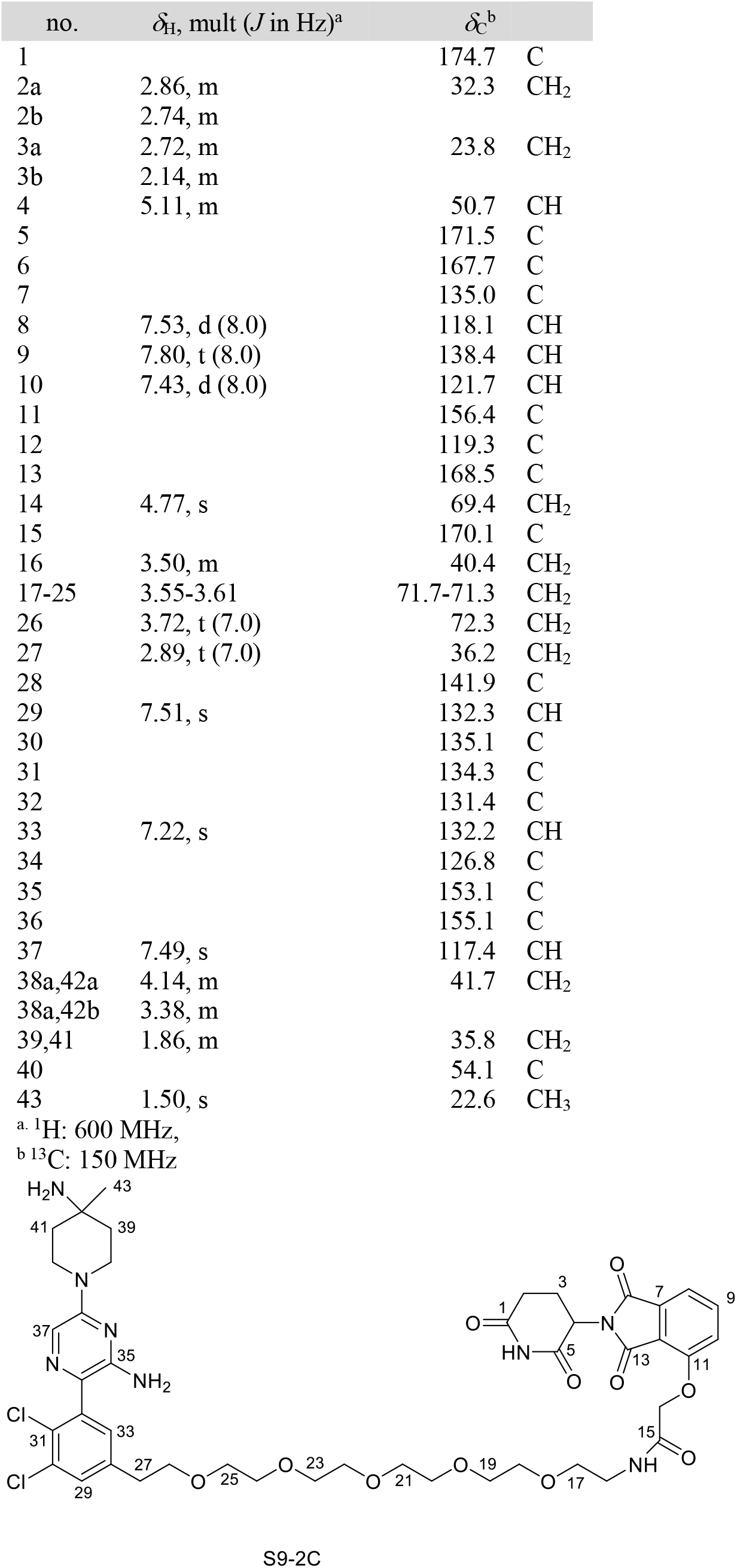
^1^H and ^13^C NMR data of S9-2C in MeOH-*d*4.

**Figure S36.**
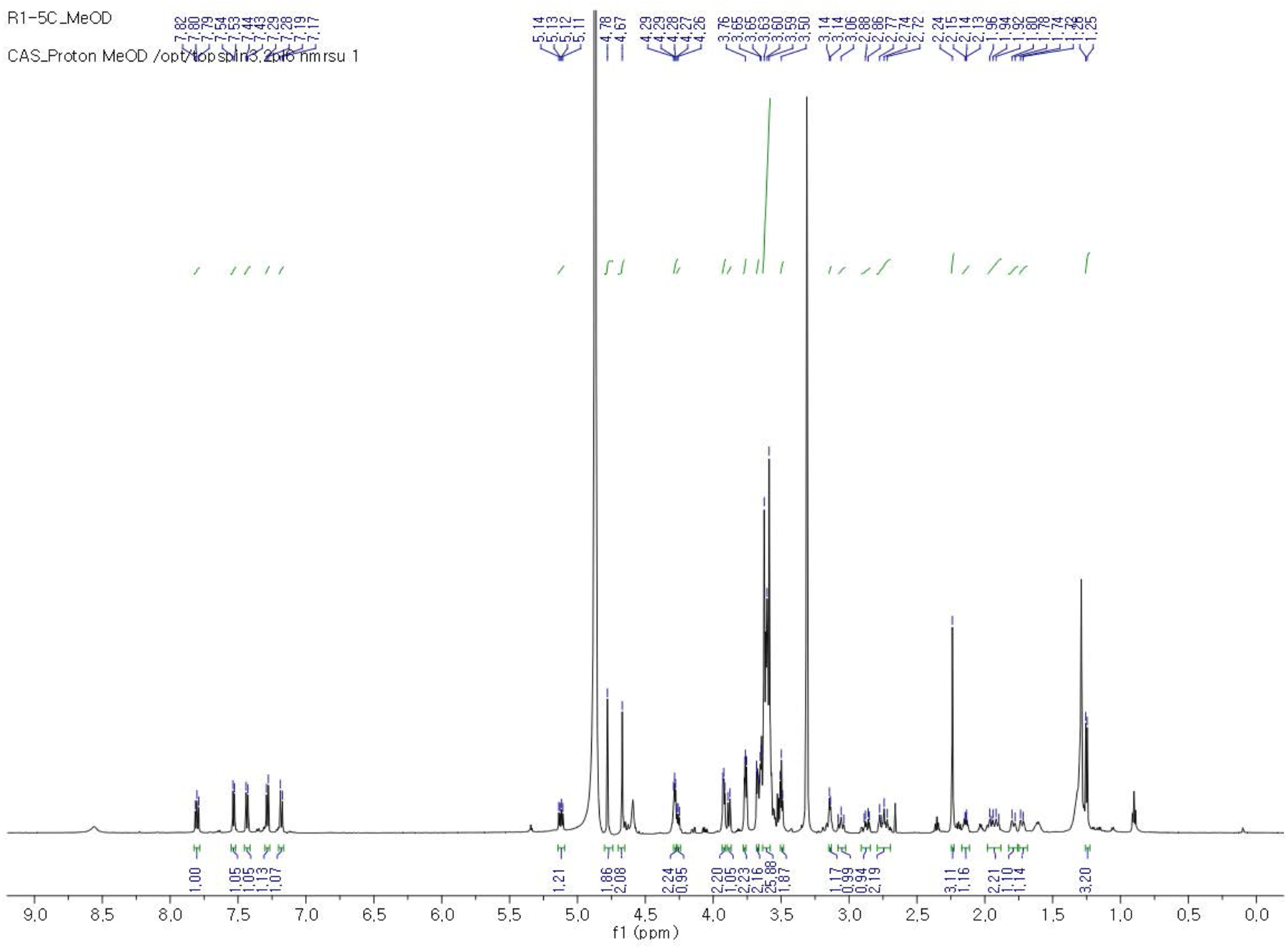
^1^H NMR spectrum of S9-3C in MeOH-*d*4.

**Figure S37.**
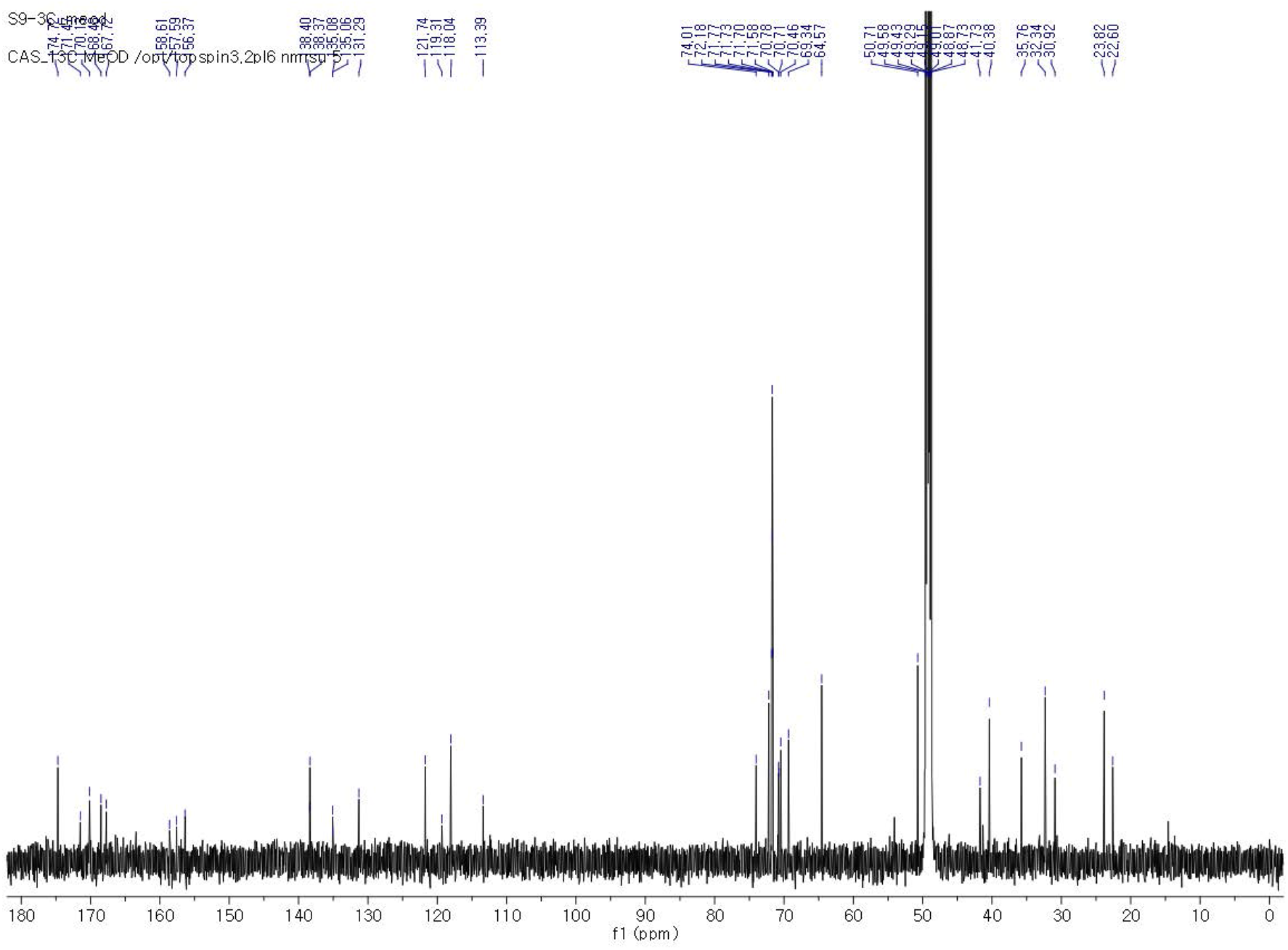
^13^C NMR spectrum of S9-3C in MeOH-*d*4.

**Figure S38.**
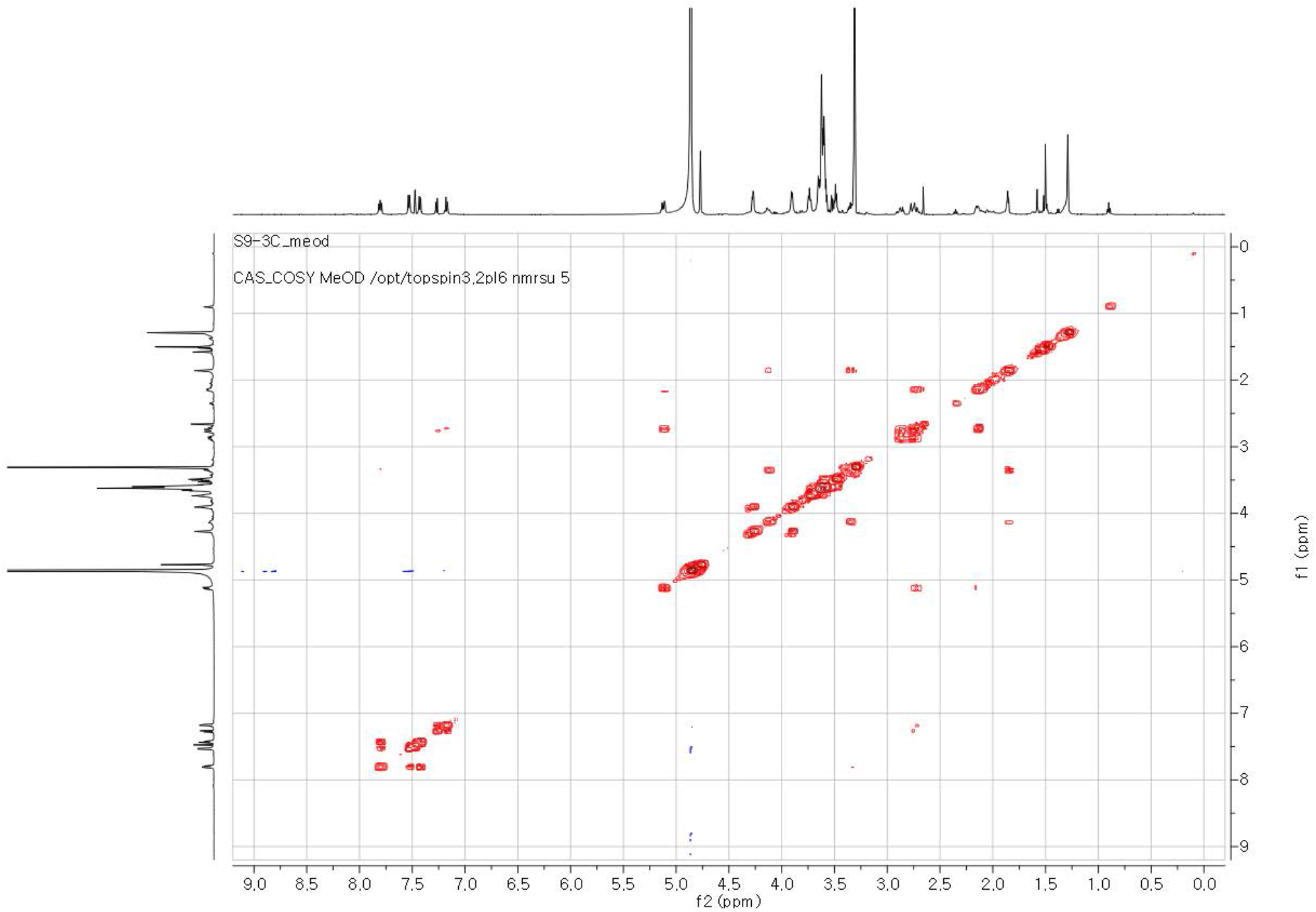
COSY NMR spectrum of S9-3C in MeOH-*d*4.

**Figure S39.**
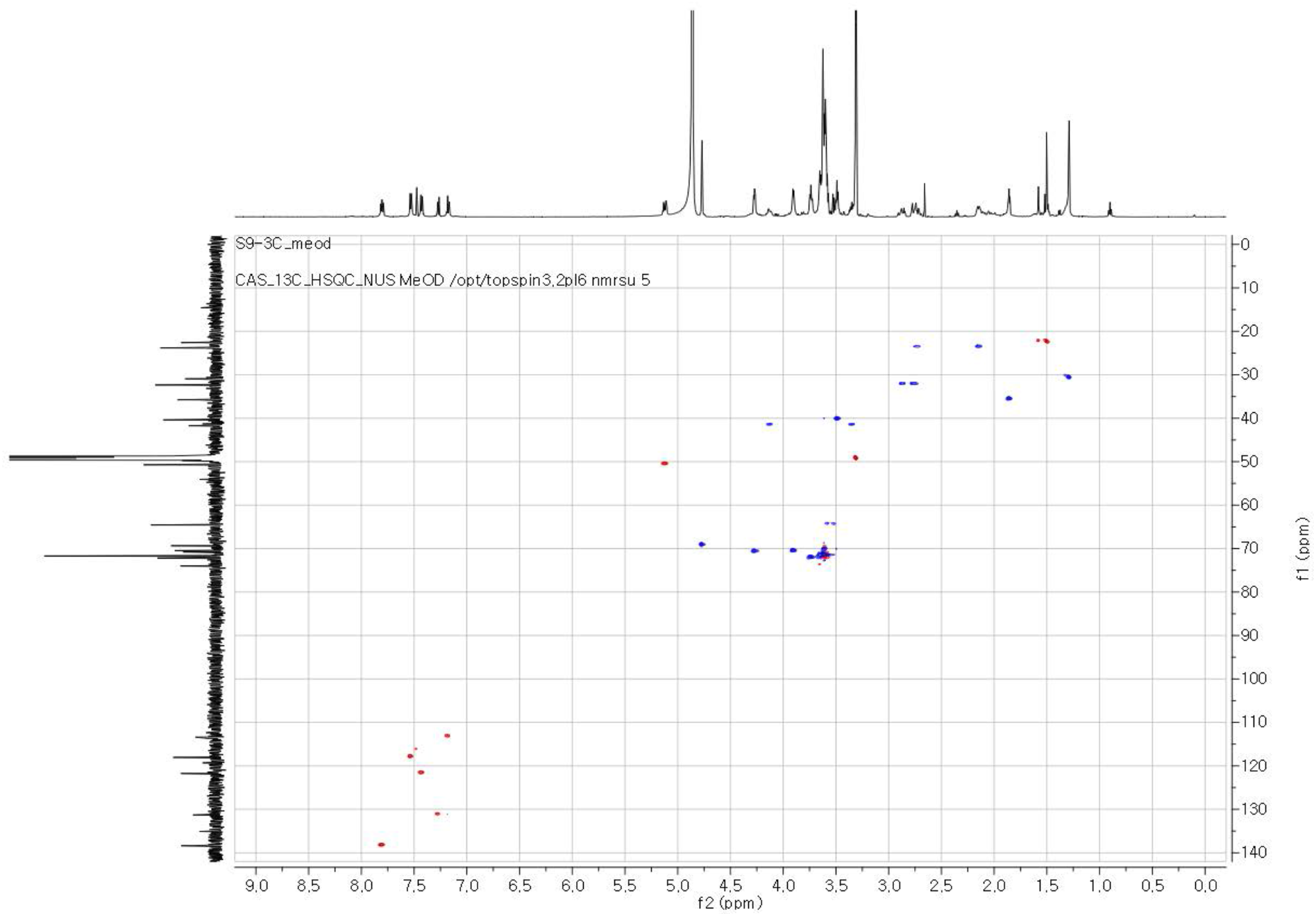
HSQC NMR spectrum of S9-3C in MeOH-*d*4.

**Figure S40.**
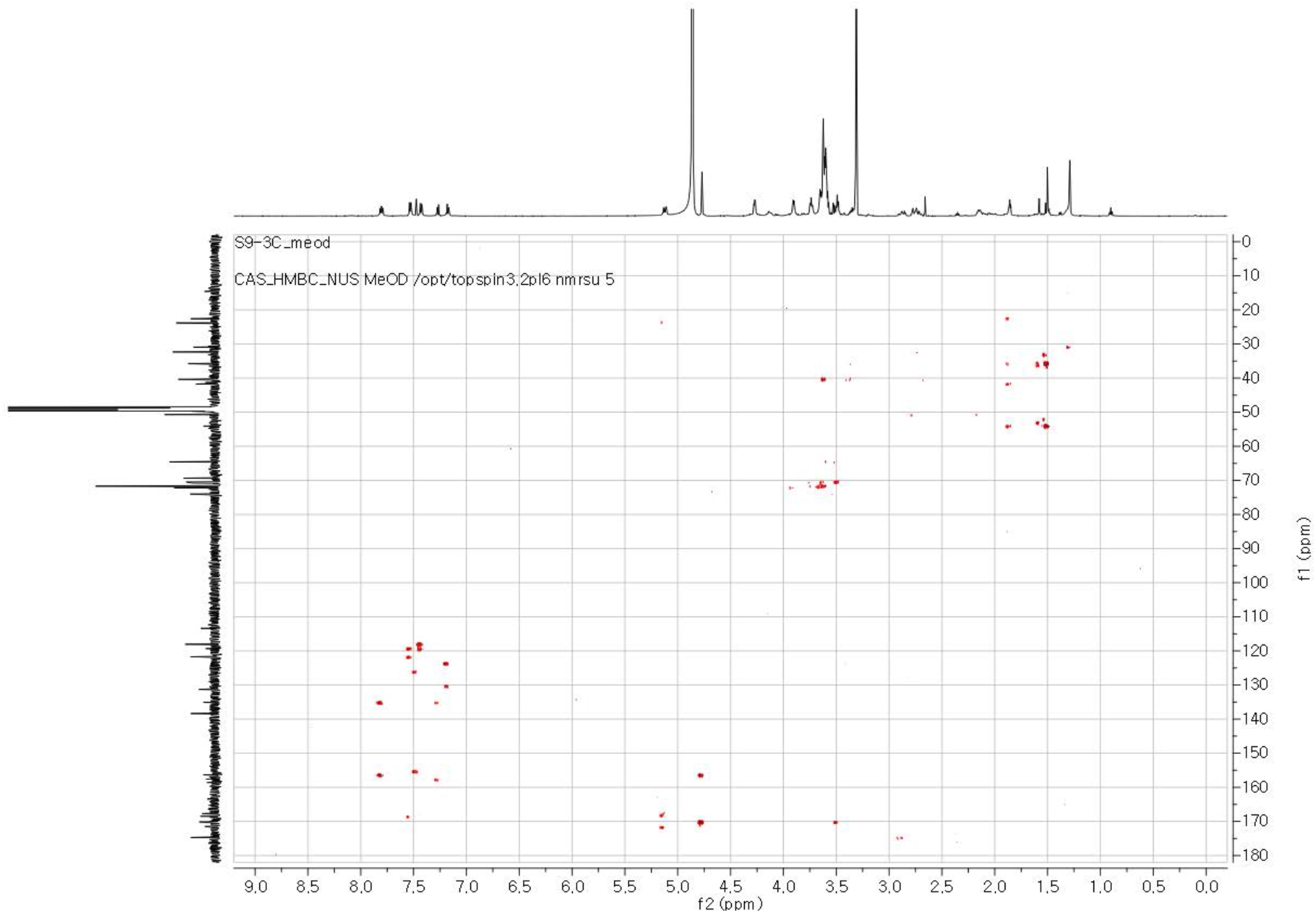
HMBC NMR spectrum of S9-3C in MeOH-*d*4.

**Table S7.** ^1^H and ^13^C NMR data of S9-3C in MeOH-*d*4.

| no. | $\delta_{\text{H}}$, mult ($J$ in Hz)<sup>a</sup> | $\delta_{\text{C}}$<sup>b</sup> | |
| --- | --- | --- | --- |
| 1 |  | 174.7 | C |
| 2a | 2.86, m | 32.3 | $\text{CH}_2$ |
| 2b | 2.74, m |  |  |
| 3a | 2.72, m | 23.8 | $\text{CH}_2$ |
| 3b | 2.14, m |  |  |
| 4 | 5.11, m | 50.7 | CH |
| 5 |  | 171.5 | C |
| 6 |  | 167.7 | C |
| 7 |  | 168.5 | C |
| 8 |  | 135.0 | C |
| 9 | 7.53, d (8.0) | 118.1 | CH |
| 10 | 7.80, t (8.0) | 138.4 | CH |
| 11 | 7.43, d (8.0) | 121.7 | CH |
| 12 |  | 156.4 | C |
| 13 |  | 119.3 | C |
| 14 | 4.77, s | 69.4 | $\text{CH}_2$ |
| 15 |  | 170.1 | C |
| 16 | 3.50, m | 40.4 | $\text{CH}_2$ |
| 17-25 | 3.57-3.62 | 72.2-71.6 | $\text{CH}_2$ |
| 26 | 3.91, m | 70.7 | $\text{CH}_2$ |
| 27 | 4.28, m | 70.8 | $\text{CH}_2$ |
| 28 |  | 157.6 | C |
| 29 |  | 123.9 | C |
| 30 |  | 135.1 | C |
| 31 |  | 130.3 | C |
| 32 | 7.27, d (8.0) | 131.3 | CH |
| 33 | 7.17, d (8.0) | 113.4 | CH |
| 34 |  | 126.4 | C |
| 35 |  | 156.4 | C |
| 36 |  | 158.6 | C |
| 37 | 7.48, s | 116.1 | CH |
| 38a,42a | 4.13, m | 41.4 | $\text{CH}_2$ |
| 38b,42b | 3.35, m |  |  |
| 39,41 | 1.86, m | 35.8 | $\text{CH}_2$ |
| 40 |  | 54.1 | C |
| 43 | 1.50, s | 22.6 | $\text{CH}_3$ |

<sup>a</sup> $^1\text{H}$ : 600 MHz,<sup>b</sup> $^{13}\text{C}$ : 150 MHz

**Figure S41.**
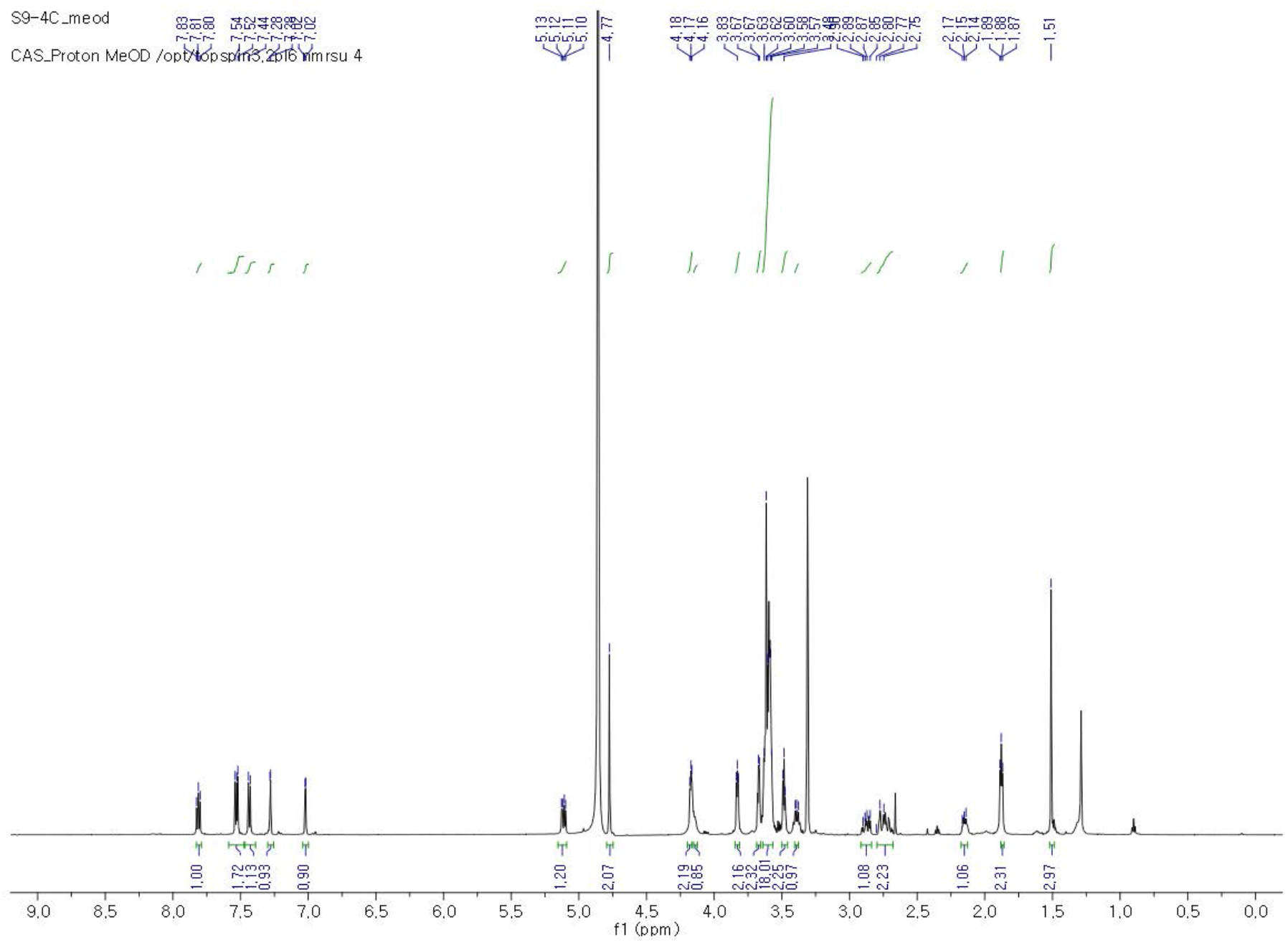
^1^H NMR spectrum of S9-4C in MeOH-*d*4.

**Figure S42.**
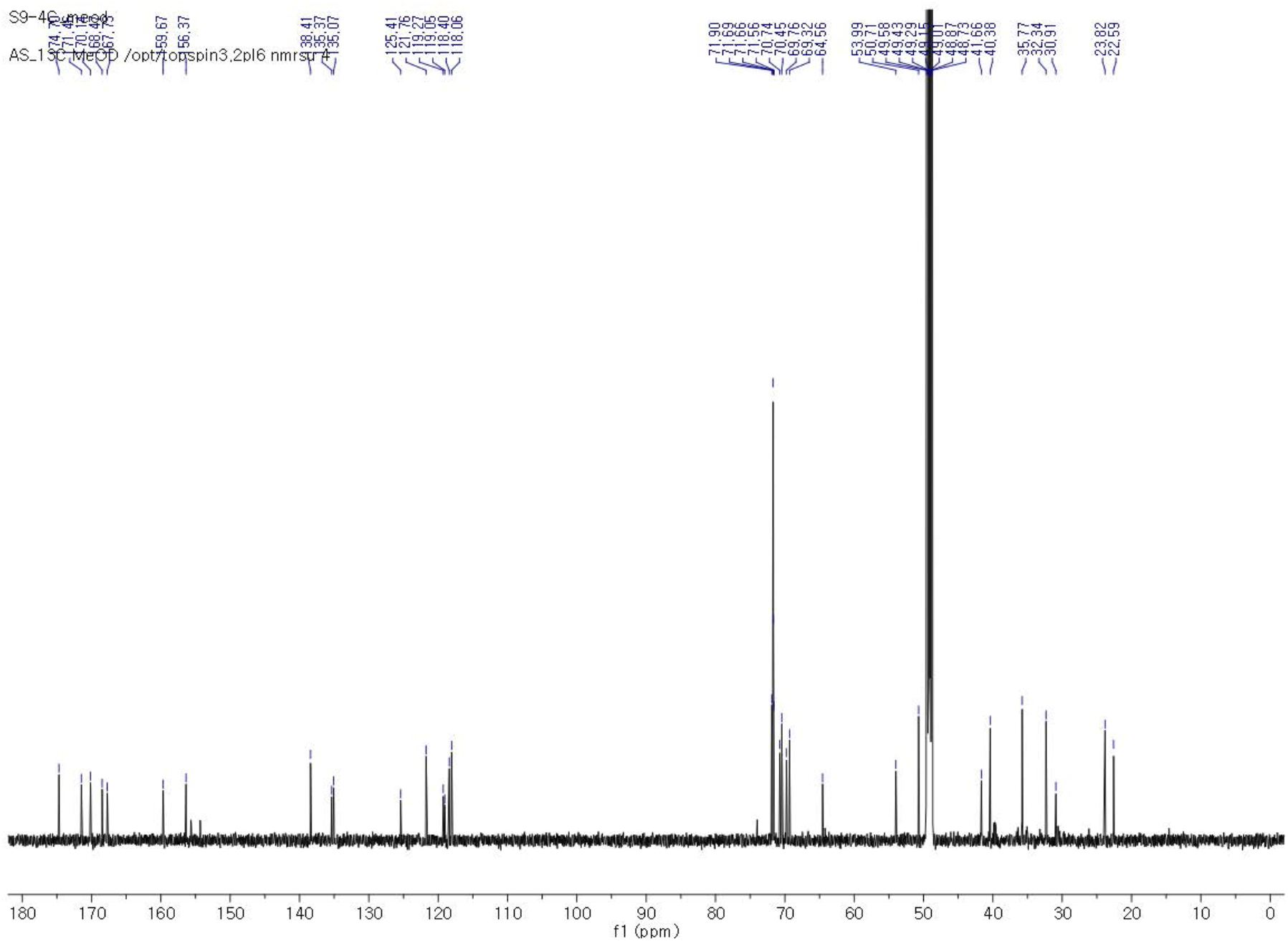
^13^C NMR spectrum of S9-4C in MeOH-*d*4.

**Figure S43.**
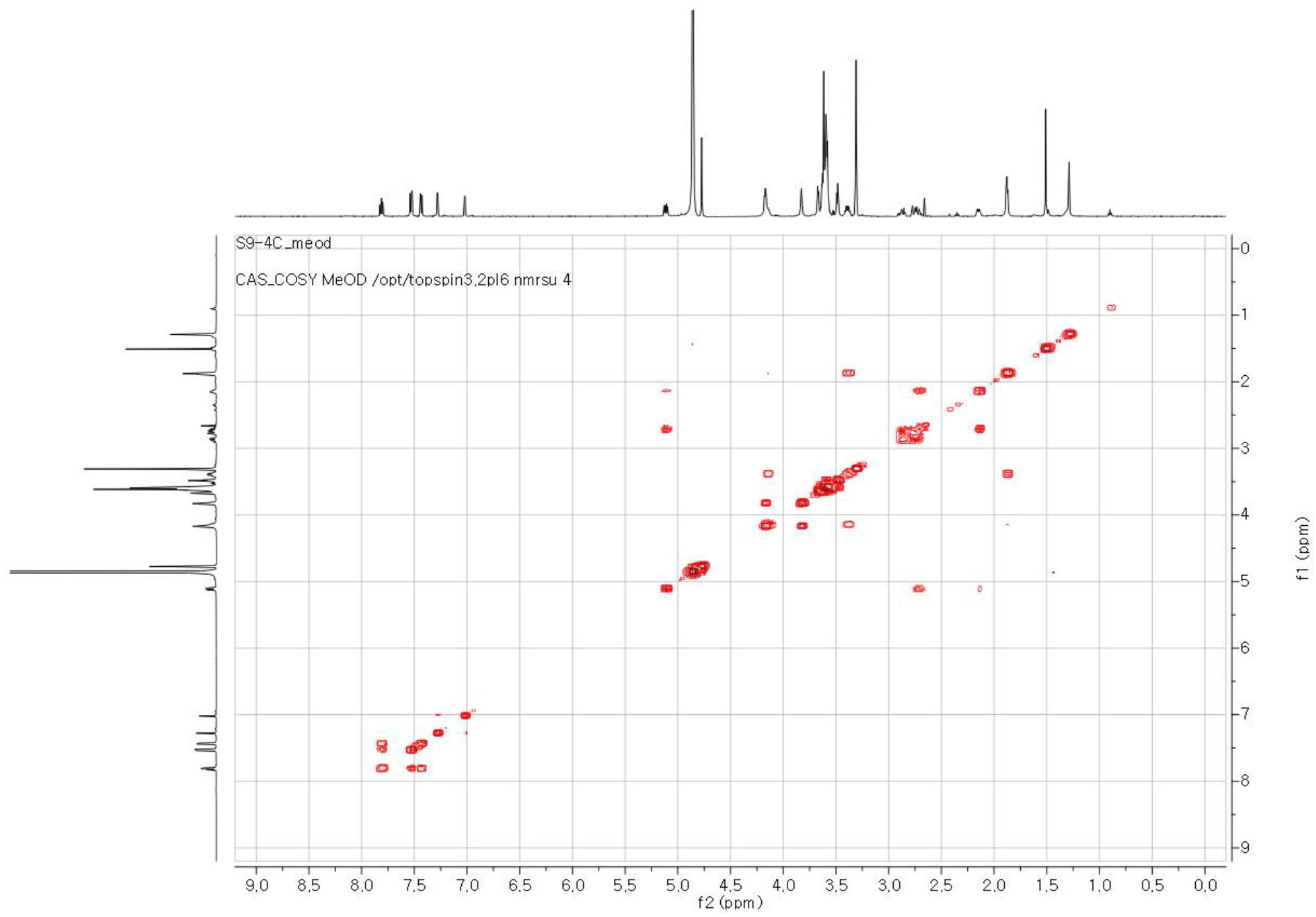
COSY NMR spectrum of S9-4C in MeOH-*d*4.

**Figure S44.**
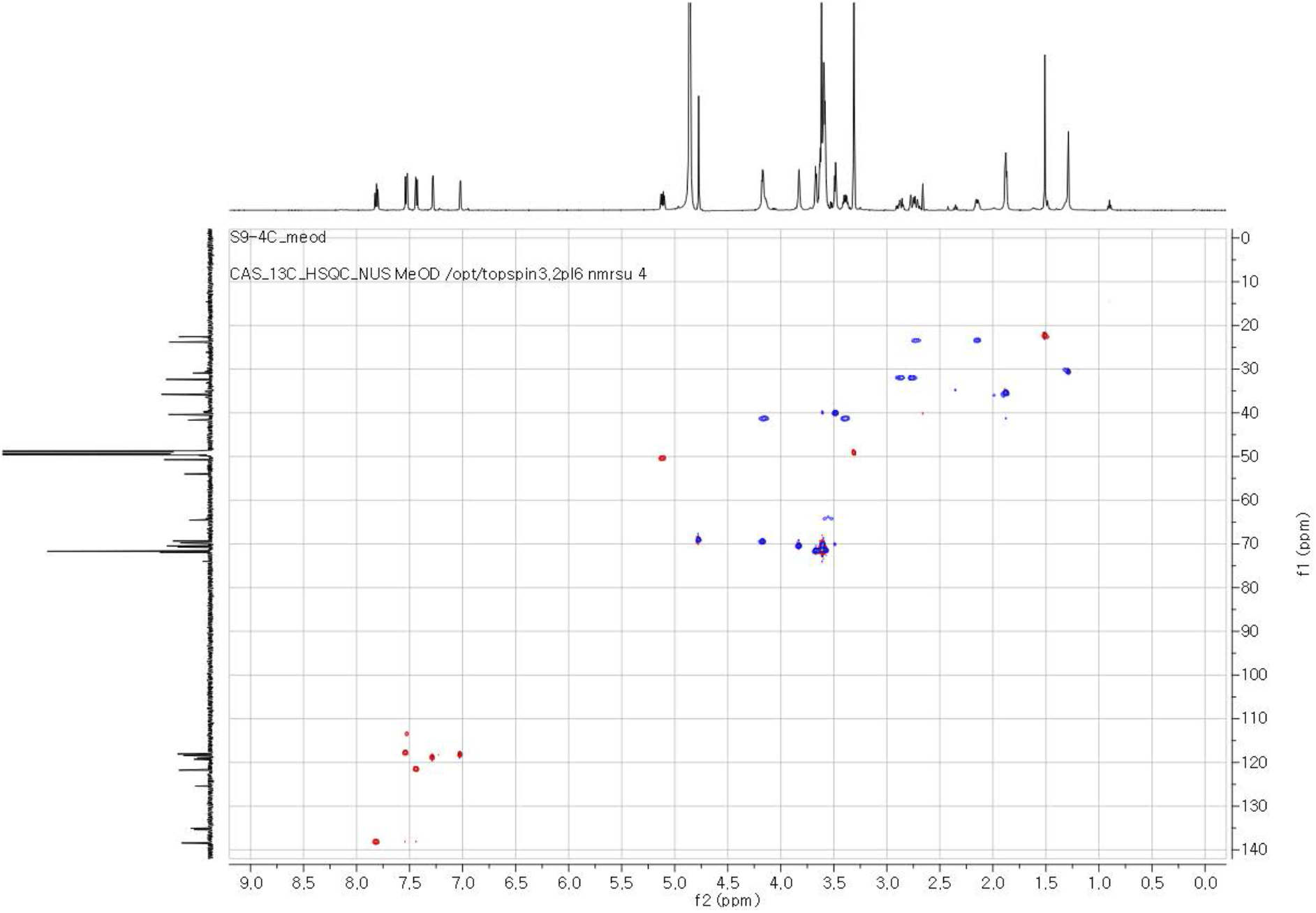
HSQC NMR spectrum of S9-4C in MeOH-*d*4.

**Figure S45.**
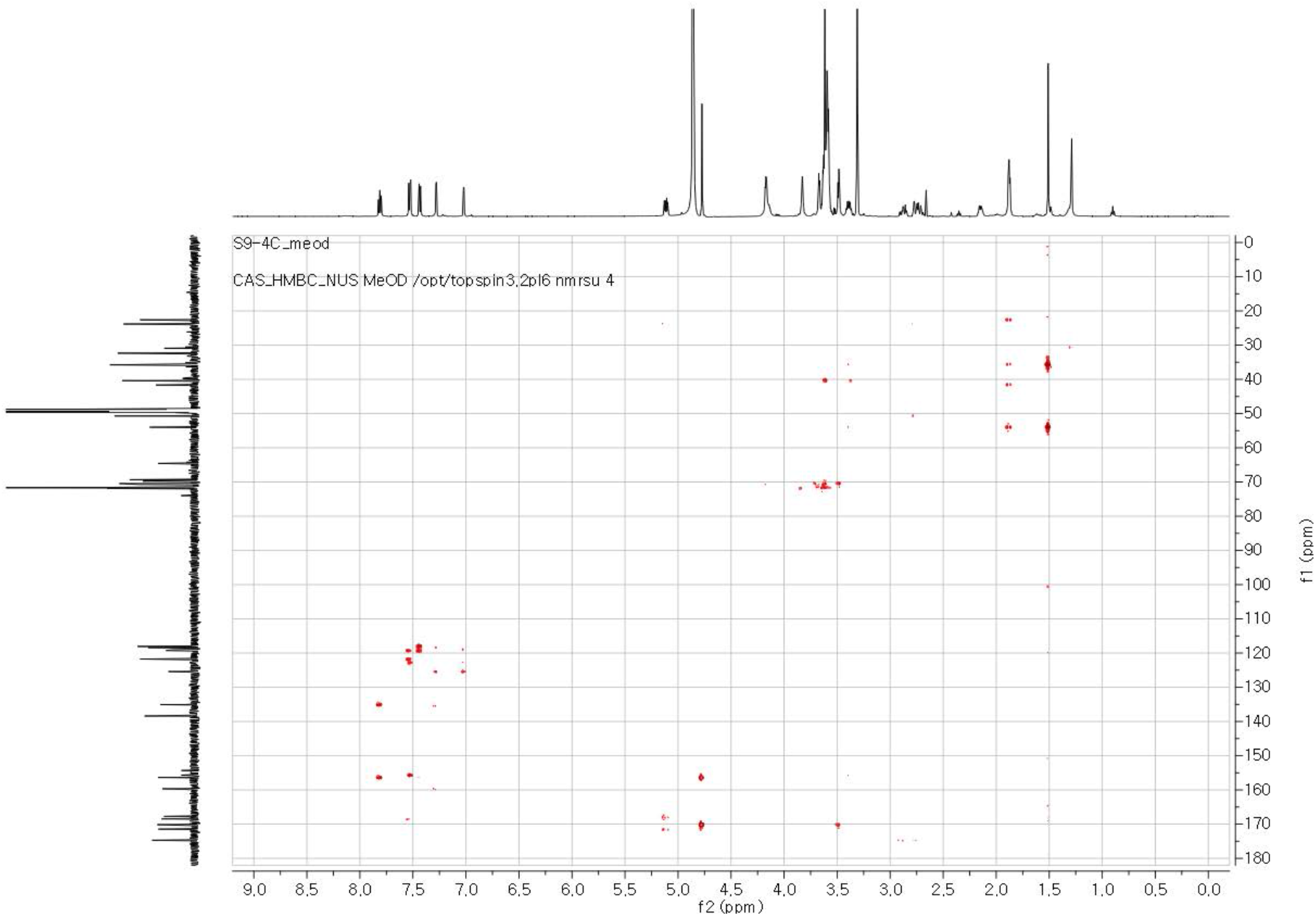
HMBC NMR spectrum of S9-4C in MeOH-*d*4.

**Table S8.**
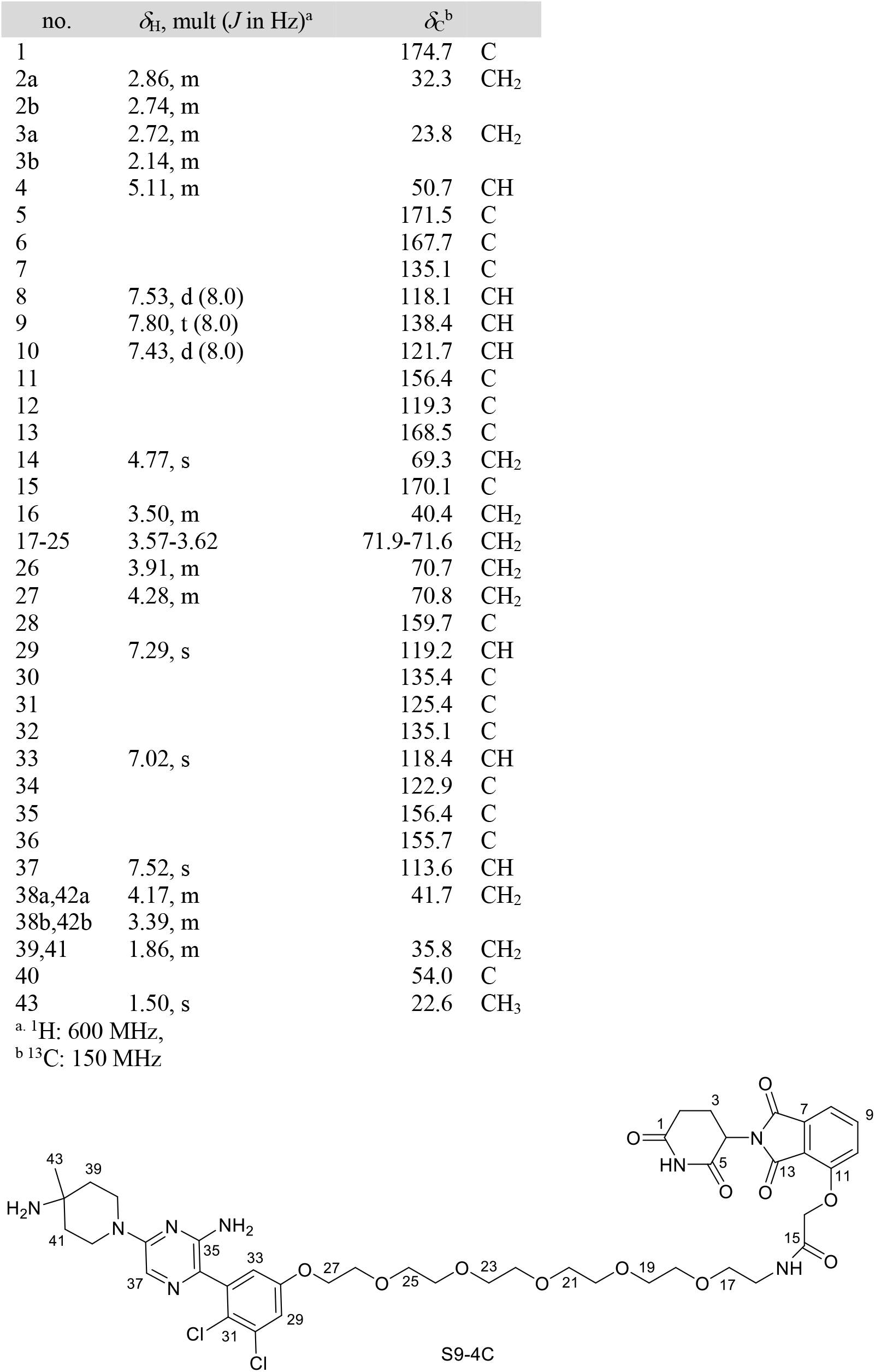
^1^H and ^13^C NMR data of S9-4C in MeOH-*d*4.

**Figure S46.**
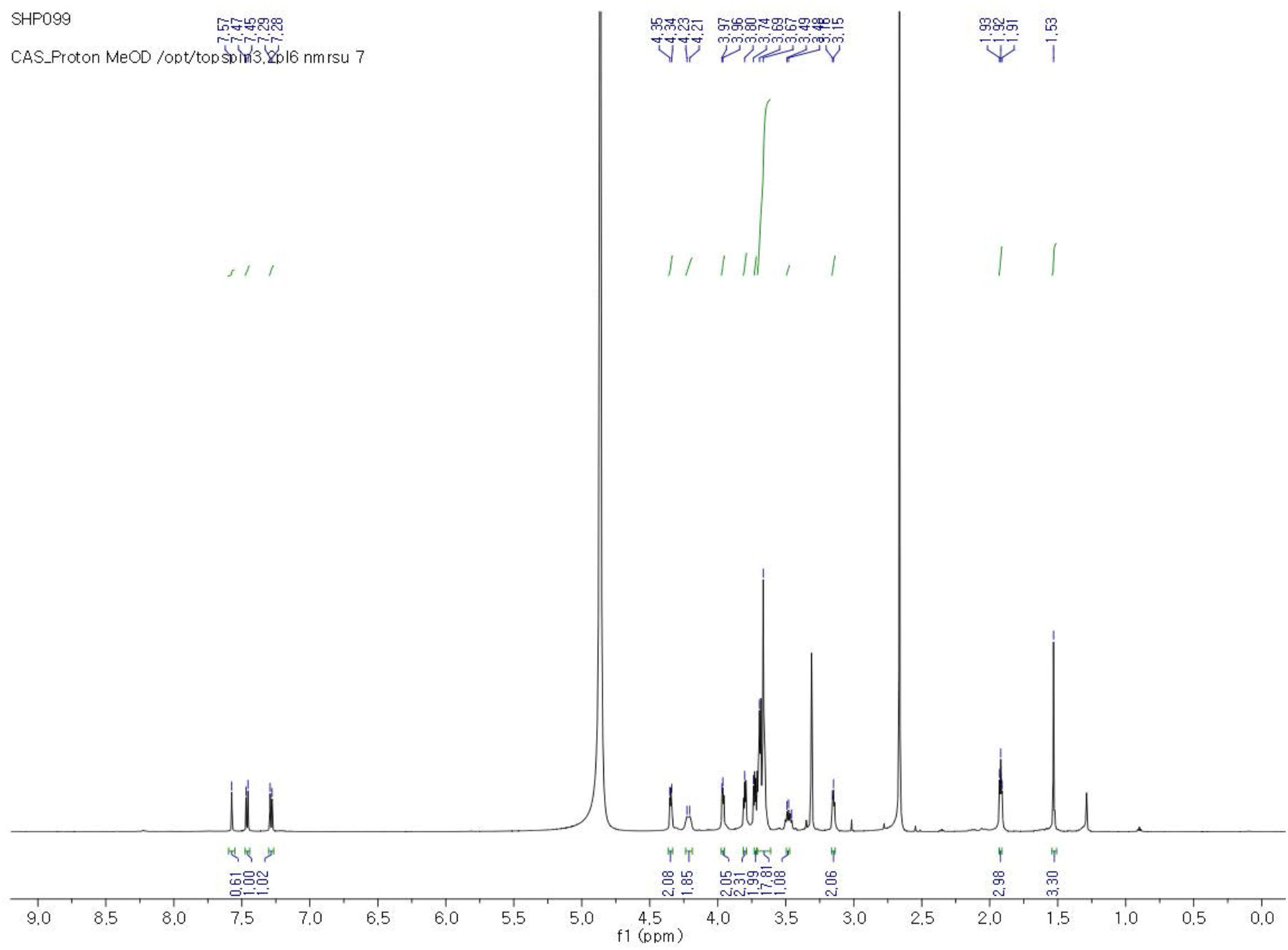
^1^H NMR spectrum of SHP099-I34 in MeOH-*d*4.

**Figure S47.**
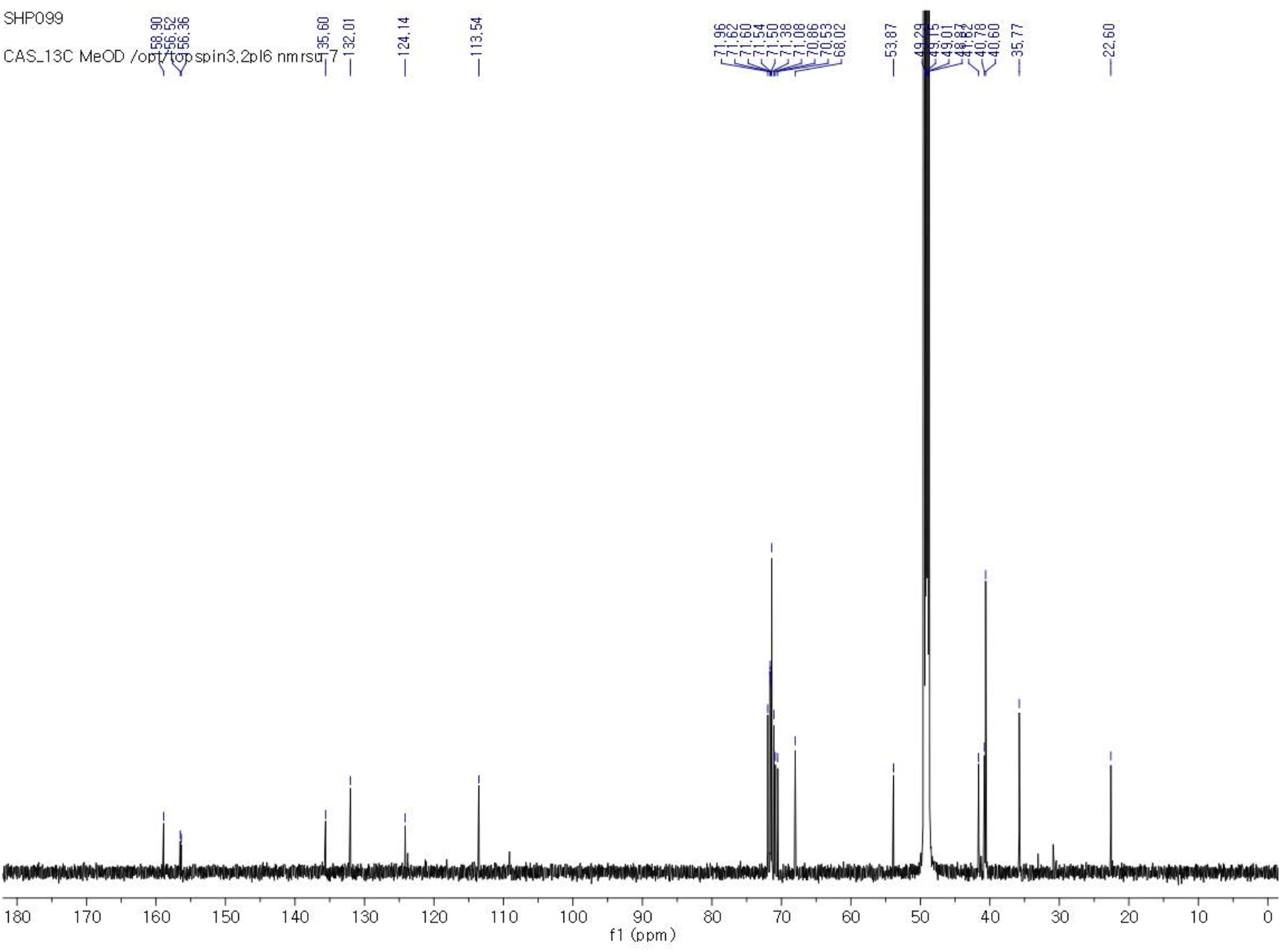
^13^C NMR spectrum of SHP099-I34 in MeOH-*d*4.

**Figure S48.**
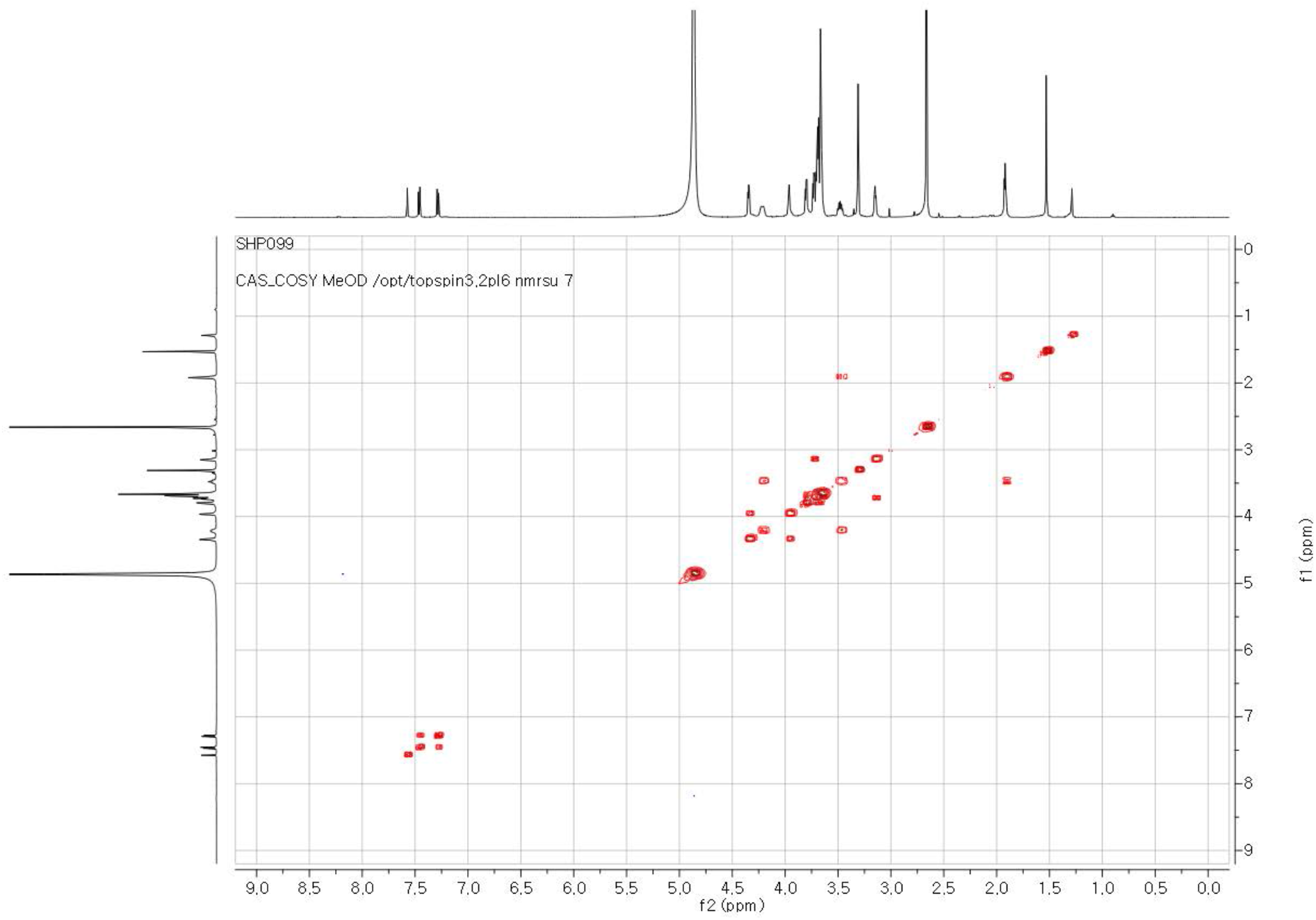
COSY NMR spectrum of SHP099-I34 in MeOH-*d*4.

**Figure S49.**
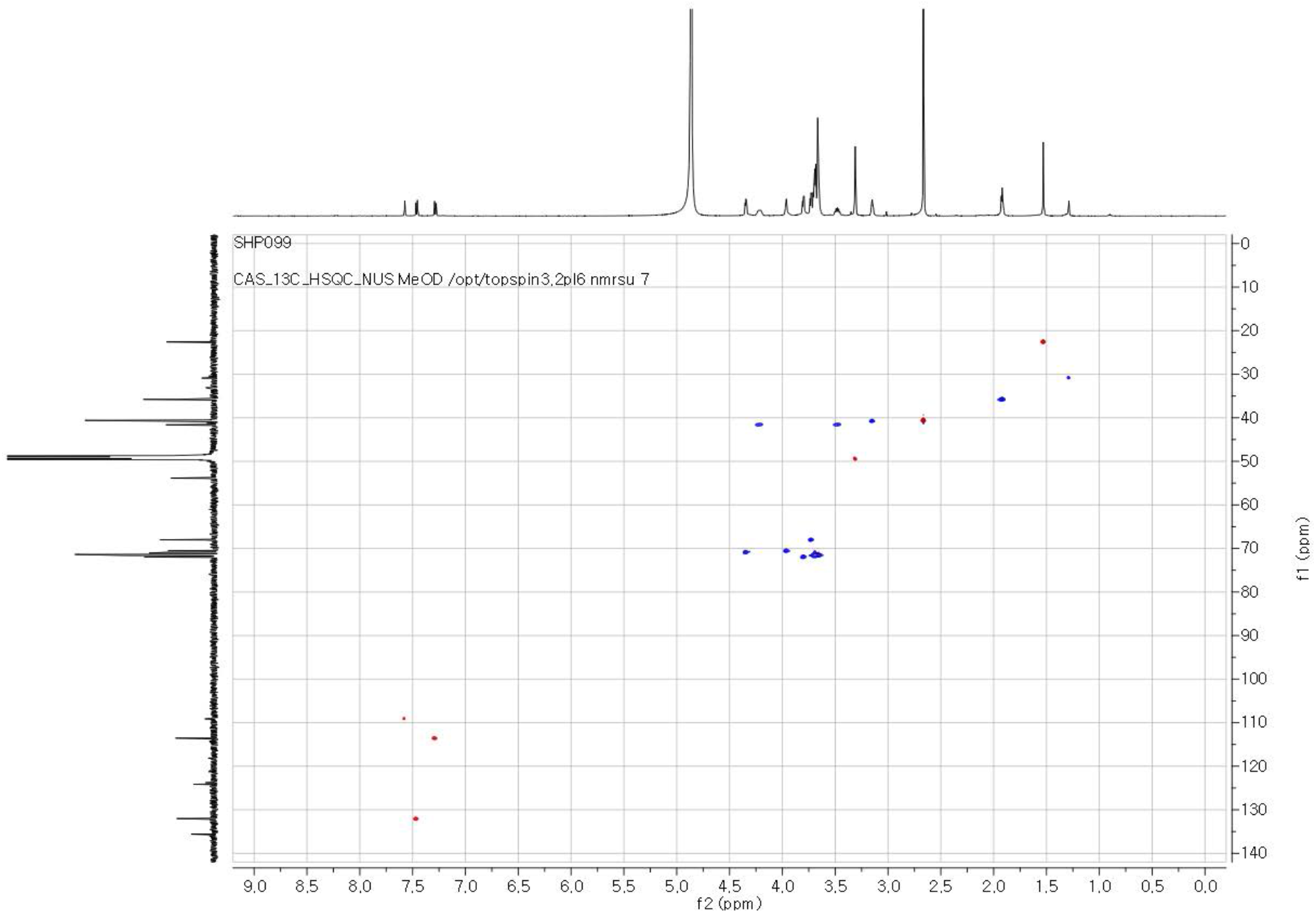
HSQC NMR spectrum of SHP099-I34 in MeOH-*d*4.

**Figure S50.**
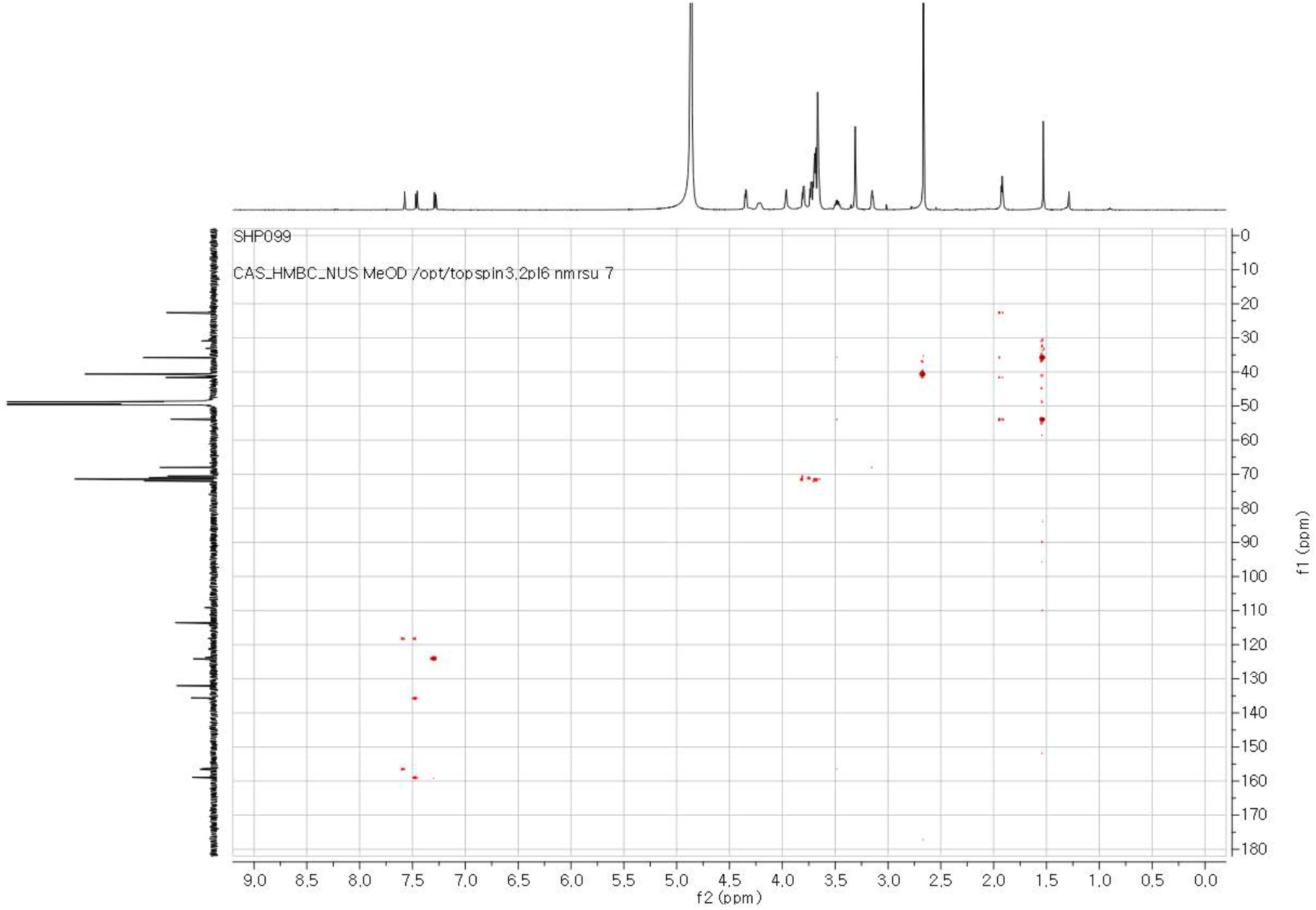
HMBC NMR spectrum of SHP099-I34 in MeOH-*d*4.

**Table S9.**
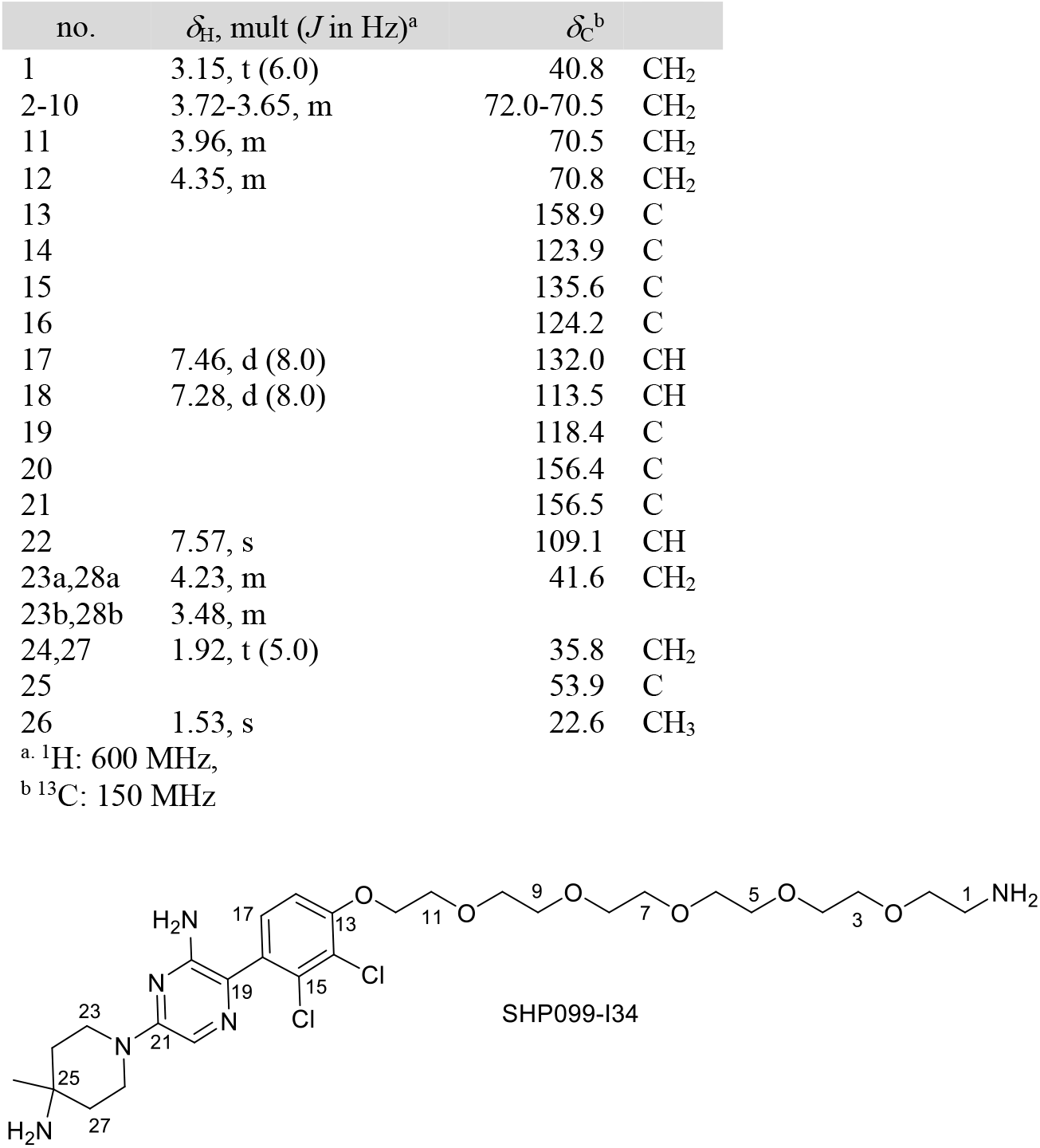
^1^H and ^13^C NMR data of SHP099-I34 in MeOH-*d*4.

**Table S10:** Expected and observed masses of PROTAC compounds.

| Compound | M | [M+H]<sup>+</sup> | [M+2H]<sup>2+</sup> |  |
| --- | --- | --- | --- | --- |
| SHP099-I34 | 630.2699 | 631.277177 | 316.142227 | theoretical |
|  |  | 631.2808 | 316.1446 | observed |
|  |  | -5.7399509 | -7.5076969 | ppm |
| S9-1C | 928.32891 | 929.336187 |  | theoretical |
|  |  | 929.3343 |  | observed |
|  |  | 2.02994355 |  | ppm |
| S9-2C | 928.32891 | 929.336187 |  | theoretical |
|  |  | 929.3354 |  | observed |
|  |  | 0.846303 |  | ppm |
| S9-3C | 944.32382 | 945.331097 |  | theoretical |
|  |  | 945.3299 |  | observed |
|  |  | 1.26569411 |  | ppm |
| S9-4C | 944.32382 | 945.331097 |  | theoretical |
|  |  | 945.3311 |  | observed |
|  |  | -0.0037024 |  | ppm |
| R1-1C | 941.31293 | 942.320207 |  | theoretical |
|  |  | 942.3201 |  | observed |
|  |  | 0.11301891 |  | ppm |
| R1-3C | 1029.36536 | 1030.37264 |  | theoretical |
|  |  | 1030.3721 |  | observed |
|  |  | 0.52068541 |  | ppm |
| R1-5C | 1117.41778 | 1118.42506 |  | theoretical |
|  |  | 1118.4246 |  | observed |
|  |  | 0.40816324 |  | ppm |

**Figure S51:**
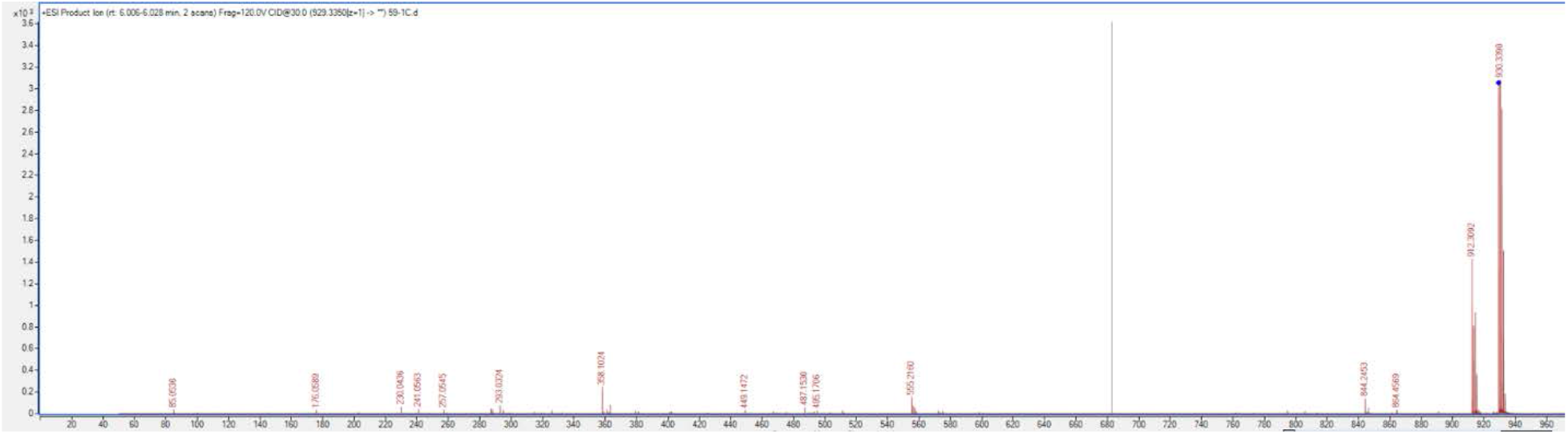
M**a**ss **Spectrum (LC-MS/MS) of S9-1C.**

**Figure S52:**
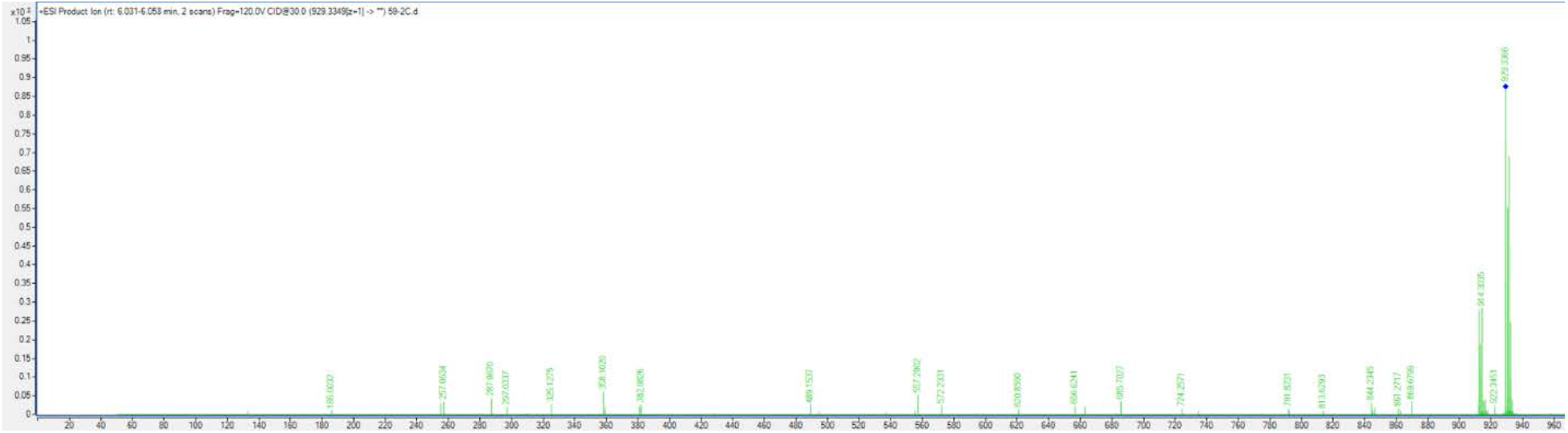
M**a**ss **Spectrum (LC-MS/MS) of S9-2C.**

**Figure S53:**
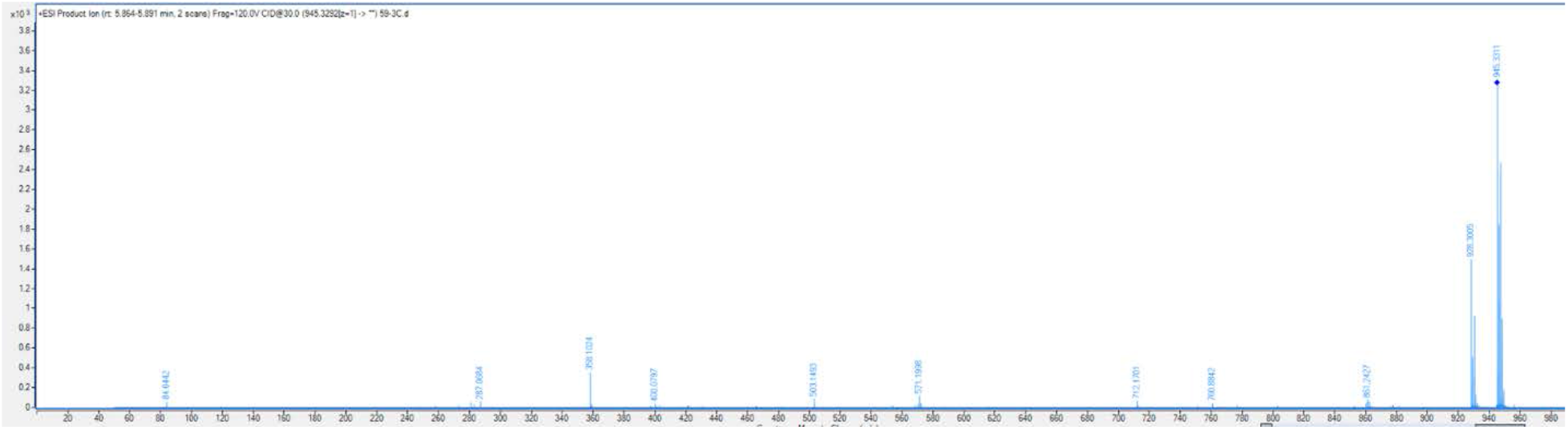
M**a**ss **Spectrum (LC-MS/MS) of S9-3C.**

**Figure S54:**
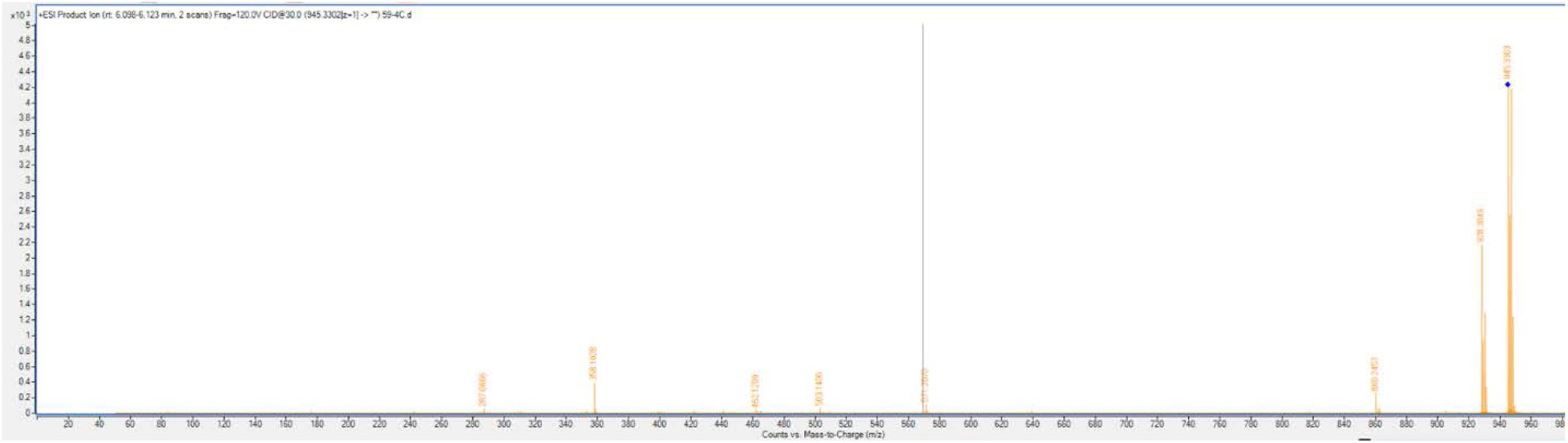
M**a**ss **Spectrum (LC-MS/MS) of S9-4C.**

**Figure S55:**
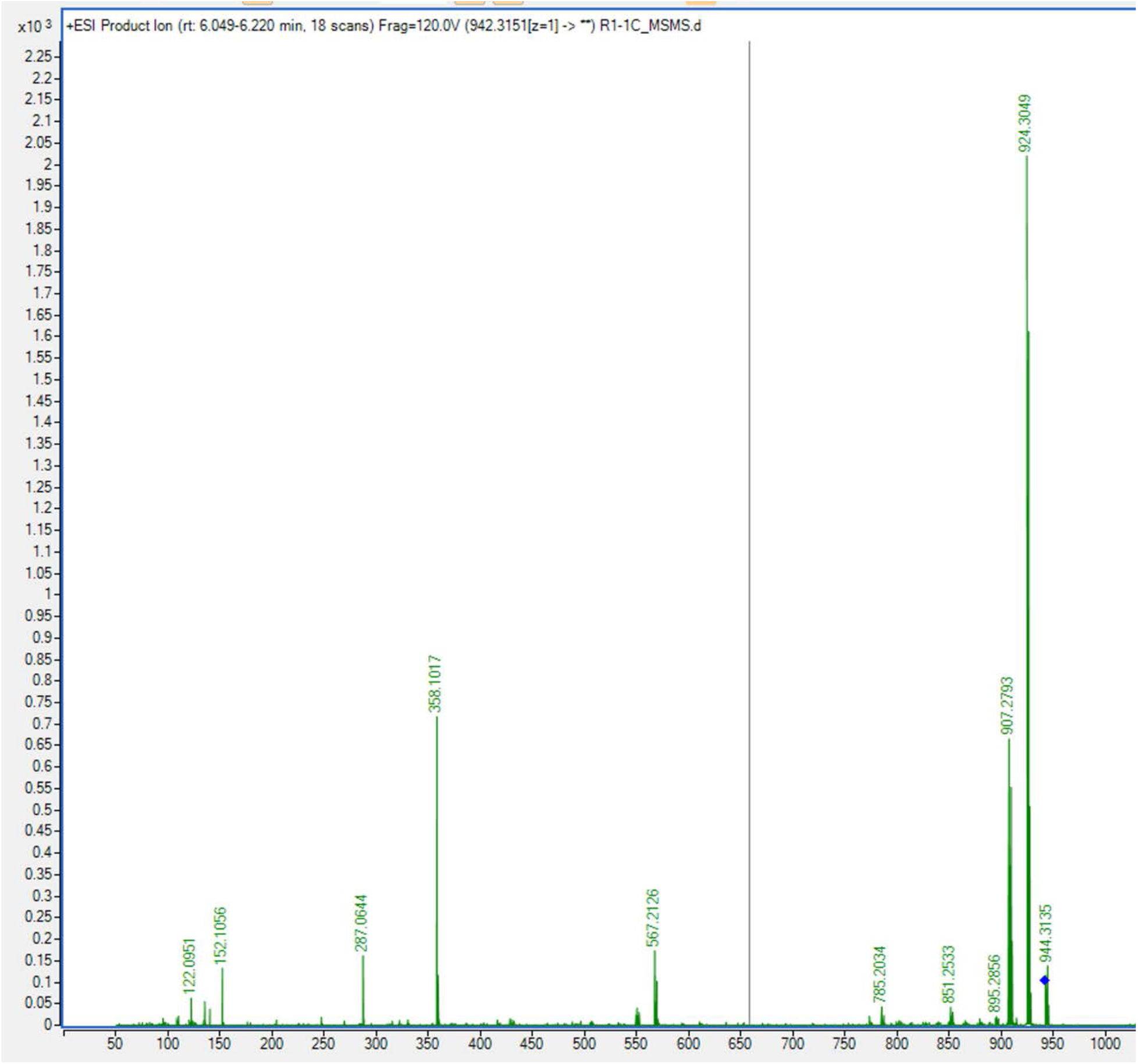
M**a**ss **Spectrum (LC-MS/MS) of R1-1C.**

**Figure S56:**
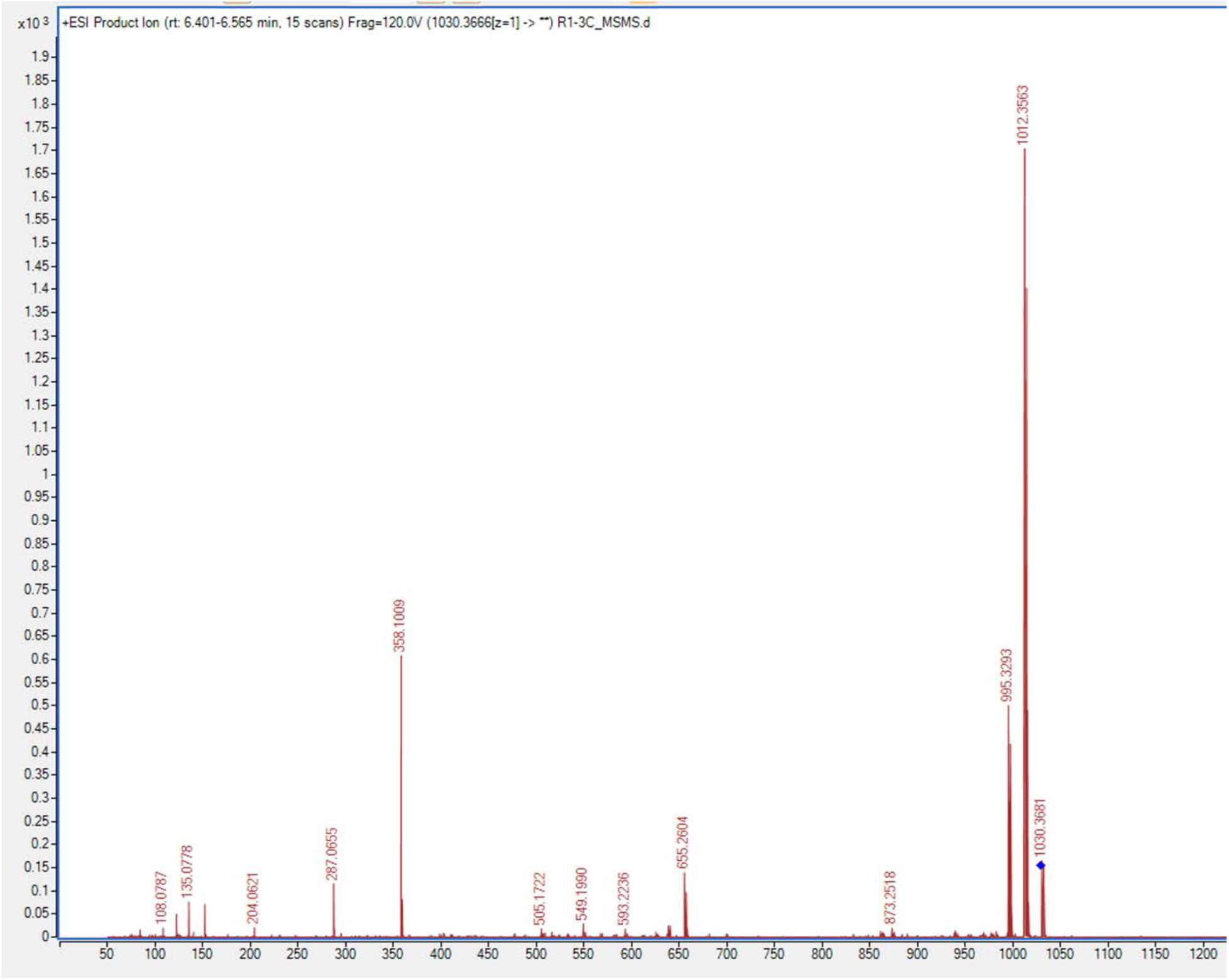
M**a**ss **Spectrum (LC-MS/MS) of R1-3C.**

**Figure S57:**
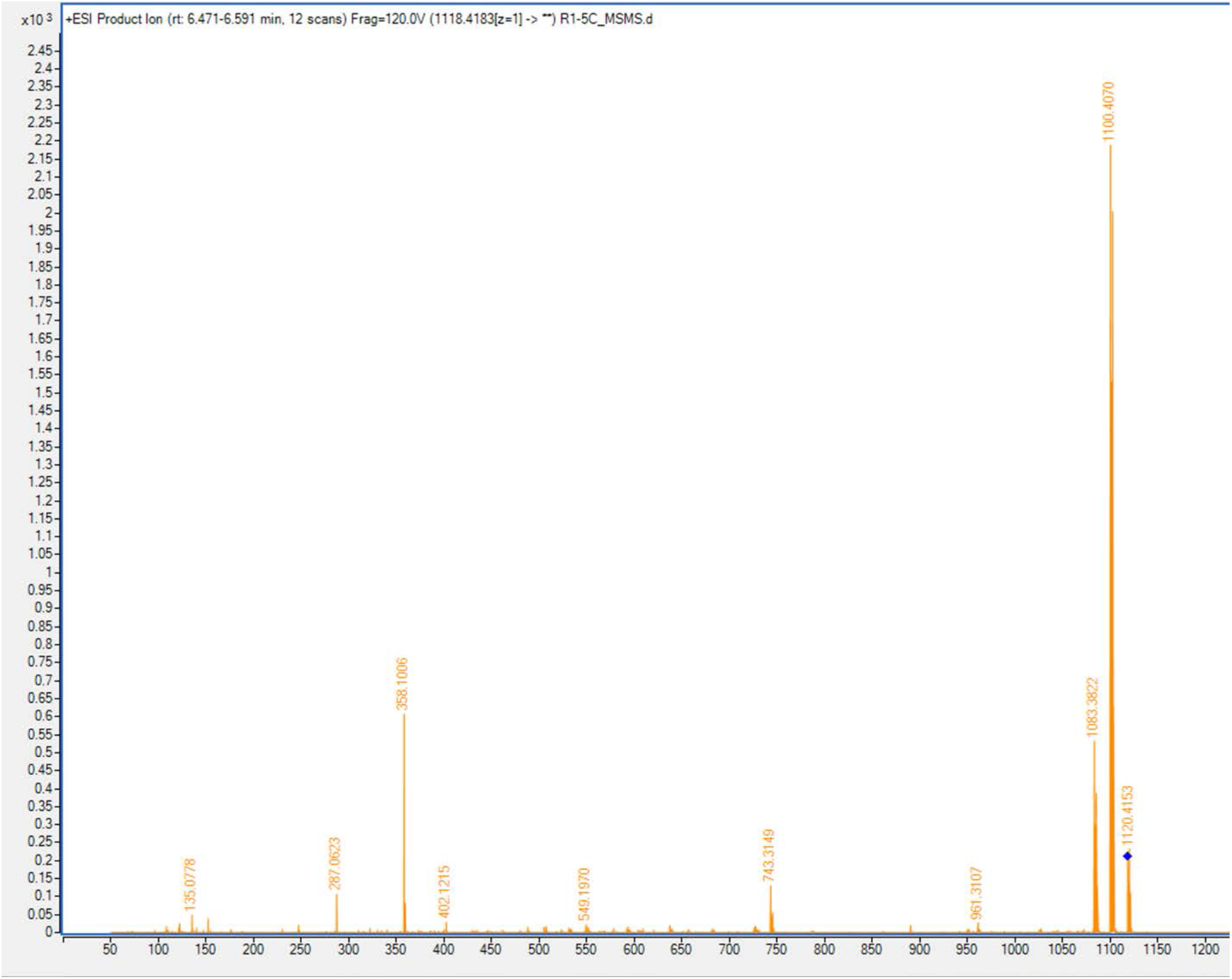
M**a**ss **Spectrum (LC-MS/MS) of R1-5C.**

